# Benchmarking scRNA-seq Copy Number Inference: A Comprehensive Evaluation and Practitioner’s Guide

**DOI:** 10.64898/2026.04.12.718050

**Authors:** Hung-Ching Chang, Yuxin Shi, Haoyu Cheng, Jian Zou, Alexander Chih-Chieh Chang, Brent T. Schlegel, Wenjia Wang, Daniel D. Brown, Fangyuan Chen, Sarah Wang, Danyang Li, Ria Sai, Noelle Michel, Steffi Oesterreich, Adrian V. Lee, George C. Tseng

## Abstract

Accurately inferring copy number variation (CNV) from scRNA-seq data is critical for identifying malignant cells, reconstructing tumor subclonal architecture, and uncovering the genomic drivers that dictate cancer cell biology. However, the performance of existing tools varies significantly, and current benchmarks lack the breadth of datasets and methods necessary to provide definitive guidance. We present a comprehensive benchmark of 12 CNV inference methods across 28 real datasets (>100,000 cells) and diverse synthetic datasets. By evaluating methods based on malignant cell classification accuracy, CNV inference accuracy, scalability, and robustness, we establish a definitive practitioner’s guideline: allele-aware methods like Numbat excel when high-quality allelic inference can be achieved, whereas expression-centric tools such as Clonalscope, CopyKAT, inferCNV, and SCEVAN remain reliable when raw sequencing data are unavailable. Our study provides both a practical decision-making framework for researchers and a public repository of standardized CNV profiles to catalyze further methodological innovation.

## Introduction

Copy number variations (CNVs) are pervasive genomic aberrations across diverse cancer types, including both solid and hematologic malignancies^1^. CNVs often accumulate during tumor evolution, with advanced and metastatic cancers typically exhibiting more complex CNV landscapes. These genomic alterations are associated with poor prognosis^2^, aggressive disease subtypes^3^, and treatment resistance^4^, and have thus emerged as key biomarkers in cancer research. Moreover, CNVs act as important drivers of intratumoral heterogeneity, posing major challenges for precision oncology and therapeutic design.

Single-cell DNA sequencing (scDNA-seq) enables direct detection of CNVs at single-cell resolution and has been instrumental in delineating subclonal architectures. However, its high cost and limited genome coverage restrict its application in large-scale studies. In contrast, single-cell RNA sequencing (scRNA-seq) provides a more accessible and widely adopted platform, offering the ability to jointly profile gene expression and infer genomic alterations at the transcriptomic level. Yet, inferring CNVs from scRNA-seq data remains challenging because gene expression is shaped by multiple layers of regulation — transcriptional control, gene silencing, epigenetic modulation, and post-transcriptional processing. The inherent sparsity and technical noise in single-cell data further complicate accurate CNV inference. Despite these challenges, CNV analysis based on scRNA-seq data offers a unique opportunity to connect genomic alterations with functional transcriptional consequences, providing multi-omics information of each single cell and deep insight into tumor evolution and cellular heterogeneity.

A variety of computational tools have been developed to infer CNVs from scRNA-seq data, broadly categorized into **expression-centric** and **allele-aware** approaches. Expression-centric methods — such as Clonalscope^5^, CONGAS^6^, CONICSmat^7^, CopyKAT^8^, copyVAE^9^, inferCNV^10^, SCEVAN^11^, and sciCNV^12^ — rely solely on gene expression, assuming that amplified or deleted genomic regions lead to corresponding up- or down-regulation of resident genes. These methods are sensitive to expression fluctuations unrelated to CNVs and often require robust normalization and reference-based denoising. In contrast, allele-aware methods — including CaSpER^13^, Chloris^14^, HoneyBADGER^15^, and Numbat^16^ — integrate allelic frequency information derived from raw reads with expression magnitude, providing a more comprehensive view of CNV states and enabling the discrimination of loss-of-heterozygosity (LoH) from copy number alterations.

Several recent studies have benchmarked CNV inference from scRNA-seq data, yet gaps remain. Schmid et al.^17^ evaluated six methods across 20 tumor datasets against paired scDNA-seq or WES-based ground truth, while Chen et al.^18^ and Song et al.^19^ assessed five methods using limited-scale multi-omics datasets. Although these efforts are foundational, their scope is constrained by the number of methods evaluated, the diversity of datasets, and other key elements to be illustrated in the next paragraph (Supplementary Table 1). Consequently, the field still lacks a definitive consensus on tool selection and best practices, particularly regarding the trade-offs between expression-centric and allele-aware approaches.

We present the most extensive benchmarking study of scRNA-seq CNV inference methods to date, providing a rigorous framework to guide tool selection across diverse experimental contexts. Our study advances the field in five key dimensions. First, we utilize a diverse compendium of 28 real-world datasets—spanning primary tumors, metastases, patient-derived organoids (PDOs), and cell lines—comprising over 100,000 cells. Notably, 20 of these datasets include matched scDNA-seq data, enabling high-resolution validation at the subclone level. Second, we employed both de novo and sampling-based simulations to systematically evaluate performance under controlled variables, such as sequencing depth and cell number. Third, we conducted a multi-layered performance assessment across cell, subclone, and bulk levels using complementary metrics including the Youden index, F1 score, and adjusted Rand index (ARI). Fourth, we profiled computational scalability in terms of resource usage efficiency (CPU/memory) to address the demands of large-scale atlas projects. We also evaluated tool usability and documentation quality. Finally, to provide a holistic practitioner’s perspective, the evaluation weights the trade-offs between leading expression-centric and allele-aware tools, culminating in a definitive tree-based roadmap to guide method selection in cancer single-cell research (Supplementary Table 1).

## Results

### Overview of Benchmarking Strategy

To systematically evaluate CNV inference, we benchmarked 12 widely used tools — comprising eight expression-centric and four allele-aware methods (Fig. 1A) — across 28 diverse cancer datasets. Our cohort spans four distinct and diverse biological sources: five primary tumors, six metastases, nine cell lines, and eight PDOs generated from both primary and metastatic tumors (Fig. 1B, Supplementary Data 1). Each source presents distinct challenges for CNV inference. For example, cell lines typically exhibit high tumor purity but lack internal normal diploid references, whereas PDOs often preserve both tumor subclonal diversity and microenvironment. To ensure “gold-standard” evaluation, ground-truth malignant cells were determined via three independent machine learning (ML) approaches^20-22^ (see Methods), while subclonal structures and CNV events were benchmarked against matched scDNA-seq or bulk WGS/WES data. We further augmented these real-world data with two types of synthetic data: de novo simulations by synthesizing scRNA-seq count data from CNV ground truth to assess CNV calling fidelity (see Methods), and more than 50 sampling-based simulations (down- or up-sampling cells or reads) to rigorously profile method scalability and robustness across varying sequencing depths and cell numbers.

**Figure 1:**
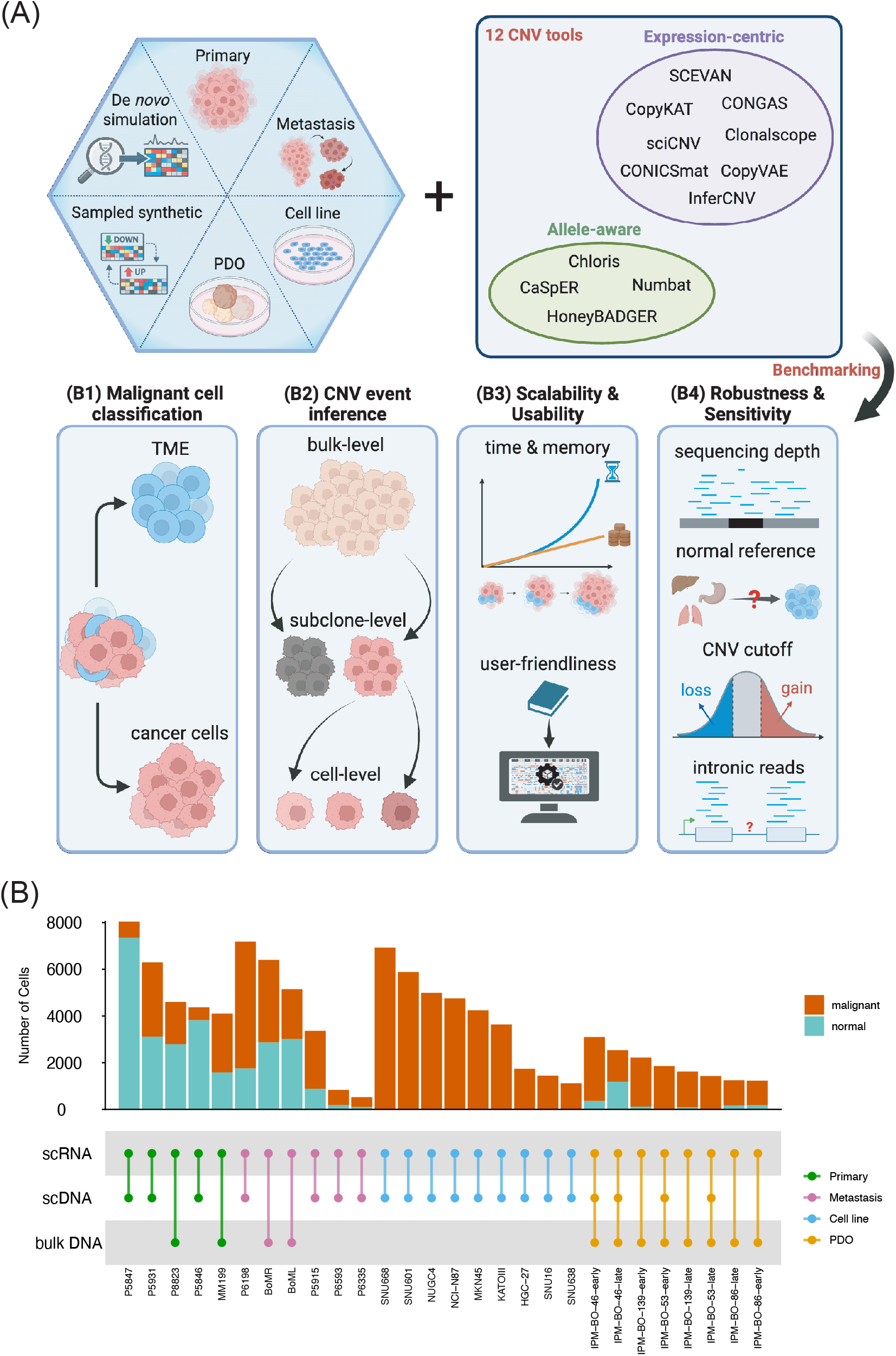
Summary of benchmarking pipeline. (A) Data: We use 4 different data sources (primary tumor, metastasis, cell line, PDO) and two types of synthetic data (de novo simulation and sampling-based synthetic data); Tools: We compare 12 CNV inference tools; Benchmarking tasks: B1-B4 represent four dimensions of evaluation criteria. (B) Summary of 28 cancer samples for evaluation, including number of single cells and data availability of scDNA-seq or bulk DNA-seq for validation.

Our evaluation and practitioner’s guideline are driven by four benchmarking dimensions (Fig. 1A): (B1) **Malignant cell classification**—accuracy of distinguishing malignant from normal cells; (B2) **CNV event inference**—accuracy of CNV detection at three resolution levels (cell, subclone and bulk), quantified by multiple metrics including the Youden index, F1 score, sensitivity, and specificity; (B3) **Scalability and Usability**—assessment of computational efficiency, documentation quality, and tunable parameters; and (B4) **Robustness and Sensitivity**—stability of performance under varying experimental and computational conditions, including sequencing depth, choice of normal diploid reference, CNV state cutoff, and inclusion of intronic reads.

Two major technical challenges in benchmarking—selecting appropriate normal references and standardizing diverse CNV outputs across tools—were addressed through a systematic preprocessing framework (see Methods), ensuring that results from different methods are directly comparable. To visualize benchmarking results, we generated a comprehensive summary table (Fig. 2) integrating all evaluation metrics across the first three benchmarking tasks (B1–B3), where lighter and larger symbols indicate higher performance. In addition, a tree-based guideline (Fig. 6) provides user-friendly recommendations of tool selection for practitioners. All benchmarking data—including performance metrics, parameter settings, and inference outputs—are publicly available in a GitHub repository (see Code Availability), with predicted CNV profiles accessible in Supplementary Fig. 9-36. To promote reproducibility, we also provide a generalizable shell-script workflow that enables users to benchmark existing or newly developed CNV inference tools under their own experimental settings.

**Figure 2:**
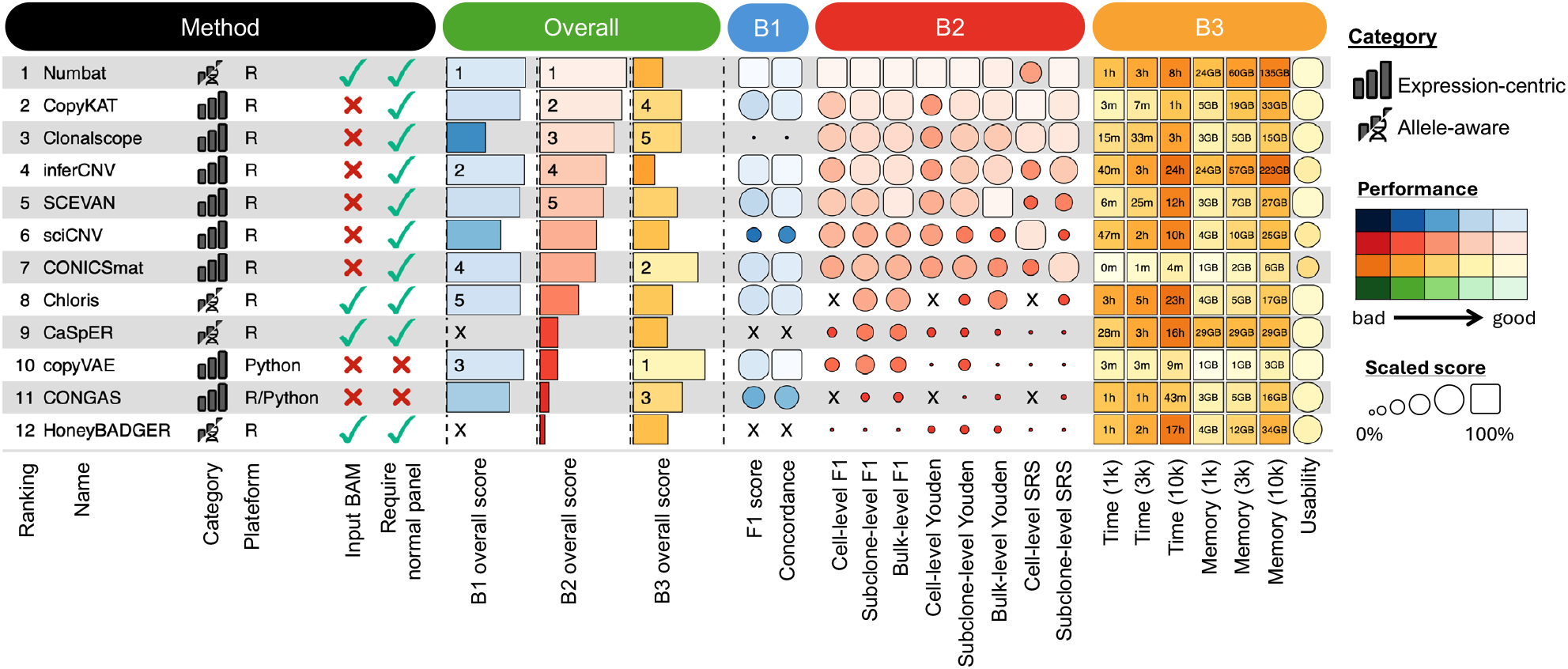
Visualization of performance evaluation results. Visualization of each method’s performance across three benchmarking tasks: B1. Malignant cell classification (blue); B2. CNV event inference (red); B3 Scalability and usability (orange). The final ranking and practitioner’s guide are primarily determined by performance in the B2 task, which represents the core objective of the study.

### Bl: Malignant Cell Classification

Accurate malignant cell identification is a prerequisite for reliable CNV calling, yet inference tools vary significantly in their classification logic. As not all methods natively provide cell-type designations, we categorized the 12 tools into three functional groups based on their capacity for malignant cell discrimination: (1) built-in classifiers, (2) score-based classifiers, and (3) non-classifiable tools. Built-in classifiers natively output malignant or normal labels as a standard pipeline component. In contrast, score-based classifiers generate comprehensive CNV profiles but lack explicit labeling functions. For these methods, following the framework established by Oketch et al.^23^, we inferred malignancy by quantifying the overall “CNV burden” using a per-cell score from gene-level CNV values. Finally, tools that provided neither a classification interface nor a complete CNV profile (e.g., CaSpER and HoneyBADGER) were deemed non-classifiable and were excluded from this specific benchmarking task.

To evaluate the accuracy of malignant cell classification, we quantified performance using F1 score, prediction concordance, accuracy, sensitivity, and specificity (Fig. 3B and Supplementary Fig. 3). The (pseudo-) ground truth labels were defined by a consensus of three state-of-the-art machine-learning-based classifiers: scMalignantFinder^20^, ikarus^21^, scATOMIC^22^. Performance was then assessed by comparing predicted malignant-cell labels against the ML-consensus ground truth. To complement this ground-truth-based evaluation, we further examined prediction concordance for each pair of methods using the Jaccard index (Supplementary Fig. 9B-36B) and average the pairwise Jaccard index for each method in Supplementary Fig. 2. This comparative analysis is particularly vital because ML-based classifiers do not explicitly incorporate CNV information.

**Figure 3:**
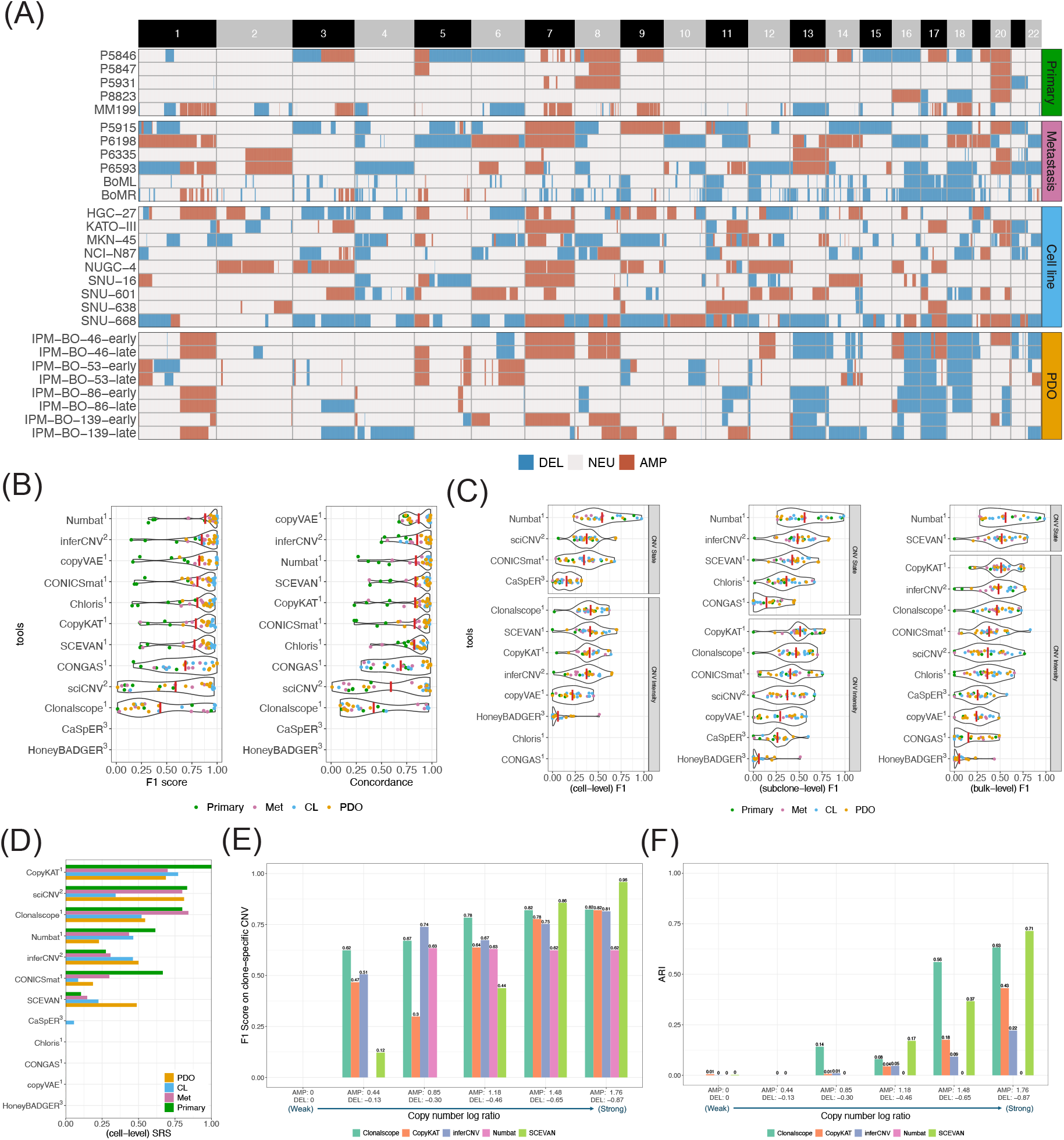
The performance of (B1) malignant cell classification and (B2) CNV inference. (A) Ground truth of CNV profiles derived by scDNA or bulk DNA. (B) Performance of malignant cell classification (C) Performance of CNV inference using Fl score in three hierarchical level: cell, subclone and bulk. (D) Performance of subclone reconstruction using cell-level SRS. (E) Performance of CNV inference in de novo simulations. (F) Performance of subclone reconstruction using ARI in de novo simulations. Note: ^1^built-in classifiers (Numbat, CONICSmat, CopyKAT, SCEVAN, copyVAE, Chloris, CONGAS, Clonalscope), ^2^CNV-score-based classifiers (inferCNV, sciCNV), and ^3^non-classifiable tools (CaSpER, HoneyBADGER).

Overall, several tools - including Numbat, inferCNV, copyVAE, CONICSmat, Chloris, CopyKAT, and SCEVAN (ranked by B1 overall score in Fig. 2) — consistently achieved high accuracy in malignant cell classification (Fig. 3B). In contrast, sciCNV sometimes produced inverted predictions (e.g., MM199, BoML, BoMR, NCI-N87 in the Supplementary Fig. 13A, 18A, 19A, 23A), and CONGAS and Clonalscope required accurate specification of CNV segments and WGS input, respectively, to achieve optimal performance. Notably, specificity was evaluated only in primary and metastatic datasets, where sufficient normal cells were available (Fig. 1B). UMAP visualizations of malignant cell classifications for ML-consensus and all evaluated tools are provided in the Supplementary Fig. 9A-36A.

Although there is not obvious difference among high-accuracy tools, performance in the B1 task varied across data sources. In particular, classification accuracy was generally higher in cell lines than in primary tumor, metastases, and PDOs, likely reflecting differences in tumor purity and genomic complexity. Cell lines are typically homogeneous and dominated by large clonal CNVs, yielding stronger signals for classification in both ML-based and CNV-based methods, whereas primary tumor, metastases, and PDOs preserve subclonal heterogeneity and often include stromal or immune cells, complicating malignant cell identification. Additionally, we observed lower prediction concordance in primary samples with limited genomic instability (e.g., P5846, P5847, P5931, P8823 in Supplementary Fig. 2). These discrepancies likely reflect cases where a low CNV burden in epithelial populations provides insufficient evidence for either CNV-based inference or ML-based classifiers to achieve high-confidence agreement.

### B2.1: Benchmarking CNV Inference via Real-World Datasets

#### Benchmarking strategy and ground truth validation

Following malignant cell classification (B1), gene-level CNV inference is evaluated using F1 score and Youden index to account for the inherent imbalance between CNV and copy-neutral regions. Performance was assessed at three resolutions -- cell, subclone and bulk - focusing on amplification (AMP) and deletion (DEL) events. Copy-neutral loss of heterozygosity (LoH) was excluded from the primary comparison due to limited support across tools (available only in allele-aware methods). To ensure a fair comparison, tools lacking built-in malignant-cell classifiers were evaluated using the ML-consensus malignant cells (Supplementary Fig. 1). Notably, while Clonalscope includes a built-in malignant-cell classifier, we still utilized the ML-consensus for its evaluation, as its native performance heavily depends on availability of matched WGS input, which was not the focus of this study.

A major challenge in this benchmark is comparing intensity-based methods (reporting continuous values) versus copy-state-based methods (outputting discrete states via hidden Markov models^10,16^ or mixture models^24^). Since discrete states are directly comparable and more intuitive for biological interpretation, we prioritized state-based evaluation. For intensity-based tools, we converted continuous values into discrete states using a semi-supervised cutoff based on the global proportion of CNV events in the DNA-seq inferred ground truth. This approach avoids the over-optimization of supervised cutoffs (e.g., maximizing F1 in Schmid et al.^17^) or the bias of arbitrary fixed thresholds (e.g., copyKAT cutoff in Guo et al.^16^).

Ground-truth CNV profiles are derived from scDNA using CopyKit^25^ or bulk DNA using CNVkit^26^ (see Fig 3A) and we evaluated their reliability using three complementary criteria: (1) Cross-modality concordance: Bulk-level profiles derived from scDNA showed strong Pearson correlations with matched bulk DNA in four PDO samples. (Supplementary Fig. 47); (2) Independent analysis validation: Several datasets in our study have been extensively characterized in previous publications (e.g., NCI-N87, MKN-45, and P5931^27,28^), allowing validation of major CNV events from the same samples; (3) Literature validation: Our results are consistent with established recurrent events from the literature, including 1q amplification and 16q deletion in breast cancer^29^ and chromosome 7 amplification in gastric cancer^30^ (Supplementary Fig. 48). Together, these evaluations establish the high fidelity of our ground-truth CNV profiles, confirming their suitability for benchmarking scRNA-seq inference tools.

#### Performance variability, tool-specific strengths and subclonal reconstruction

In Fig. 3C (F1 score) and Supplementary Fig. 6 (Youden index), comparison against ground truth revealed substantial variability in method performance. Among all tools, Numbat, an allele-aware method, achieved the highest overall performance, with average F1 scores at the cell and subclone levels exceeding other tools by at least 0.1. Top expression-centric methods (Clonalscope, CopyKAT, inferCNV, and SCEVAN) followed with generally reliable and comparable accuracy. Intriguingly, although Numbat dominated in most samples, its performance was unstable, with clear inferiority observed in a subset of samples (black arrows in Supplementary Fig. 4). This trend is further explored in the Practitioner’s Guide section.

We observed that event-specific sensitivities can be distinct in different methods: SCEVAN demonstrated superior performance in detecting deletions, while Numbat and Clonalscope were more sensitive to amplifications (Supplementary Fig. 7). Notably, accuracy remained high in technically challenging datasets, such as the eight multiplexed breast cancer PDOs, suggesting that these methods are robust across diverse library-preparation protocols (Supplementary Fig. 4). Although several intensity-based approaches achieved comparable performance to the top state-based methods (inferCNV, Numbat, SCEVAN), the advantage of state-based methods were still obvious, particularly in their interpretability and independence from cutoff selection. By contrast, intensity-based methods require thresholding to distinguish CNV from neutral regions, yet optimal cutoffs are rarely generalizable in practice. The robustness to CNV cutoff in intensity-based methods will be further evaluated in section B4.

We next assessed subclonal reconstruction by quantifying how well predicted CNV profiles under cell or subclone level matched to scDNA-defined ground-truth subclones. For tools without built-in clustering, we applied hierarchical clustering and matched labels to the ground-truth subclone yielding the highest F1 score (threshold at >0.6 to exclude spurious assignments). CopyKAT, sciCNV, Clonalscope, and Numbat emerged as the most effective methods for subclone reconstruction based on the cell-level subclone reconstruction score (SRS) (Fig. 3D). However, we observed subclone-level SRS (Supplementary Fig. 8) significantly underperformed cell-level SRS, especially sciCNV and CopyKAT. This discrepancy likely stems from clustering algorithms failing to preserve rare subclones or the inherent subclonal differences between cells sequenced in scDNA vs. scRNA modalities. We note that SRS is generally moderate performance, which is expected since the subclone result can be impacted by many intermediate steps. And the accuracy of (pseudo-) ground truth can also impact the performance (see discussion).

### B2.2: Benchmarking CNV Inference via De Novo Simulation

To precisely quantify CNV-inference sensitivity and assess factors influencing subclonal reconstruction, we evaluated all methods using de novo synthetic datasets with pre-defined subclone structures and a board range of CNV effect sizes (see Supplementary Methods). Although scDNA-seq provides ground-truth profiles in real-world datasets, discrepancies arise from two factors: (1) modality-specific subclones, as scDNA and scRNA are sequenced from different individual cells; and (2) signal ambiguity in genomic regions with high technical noise. De novo simulations overcome these limitations by providing an absolute reference for both CNV boundaries and subclonal assignments.

We simulated synthetic scRNA-seq datasets using scDesign3^31^, a generative simulation framework that learns gene-level expression distributions and gene-gene dependency structures from real scRNA-seq data. Specifically, normal tissue expression from P5931 sample was used as the transcriptional backbone, while the subclonal CNV structure of the SNU638 gastric cancer cell line was used as a template (see Methods). With a definite ground truth, we evaluated clone-specific CNV accuracy using the F1 score and clustering performance via ARI. Under weak CNV signals (log-ratios near 0), Clonalscope exhibited the highest detection sensitivity, followed by CopyKAT and inferCNV (Fig. 3E). As the effect size increased, SCEVAN emerged as the top performer, largely due to its superior precision in CNV breakpoint detection (Supplementary Fig. 42). Regarding subclonal clustering performance, SCEVAN, Clonalscope, and CopyKAT achieved the highest ARI values (0.71, 0.63, 0.43, respectively) under strong CNV signals (Fig. 3F). Notably, inferCNV tended to over-partition the cell populations, suggesting that the default Leiden clustering parameters may require further optimization for specific datasets.

We note that our de novo simulation primarily modulated gene expression magnitude without altering heterozygous allele frequencies. The utility of allele-specific signals was diminished, and it inherently disadvantages allele-aware methods like Numbat, which rely on B-allele frequency (BAF) shifts for refinement. Taken together, these results suggest that SCEVAN provides the most reliable CNV breakpoint detection and subclone reconstruction under high-signal conditions, while Clonalscope and CopyKAT remain robust under weaker signals.

### B3: Scalability and Usability

Beyond malignant cell classification accuracy and CNV inference fidelity, computational overhead is a critical factor for large-scale applications. To systematically benchmark runtime and memory usage, we generated a series of synthetic datasets based on the SNU601 cell line (*n =* 5,885 Cells), by down-sampling or up-sampling to vary in size from *n =* 1,000 *to* 20,000 cells (1k, 2k, 3k, 4k, 5k, 10k, and 20k, with three replicates each). All methods, including required preprocessing steps, were executed using default parameters to record total compute time (Fig. 4A) and peak memory consumption (Fig. 4B). To quantify scaling trends, we applied log-log regression models to runtime and memory usage as a function of cell count (Fig. 4C for five top performers; Supplementary Fig. 44 for others). To maintain practical relevance, we imposed a 24-hour runtime limit; tasks exceeding these limits were right-censored.

**Figure 4:**
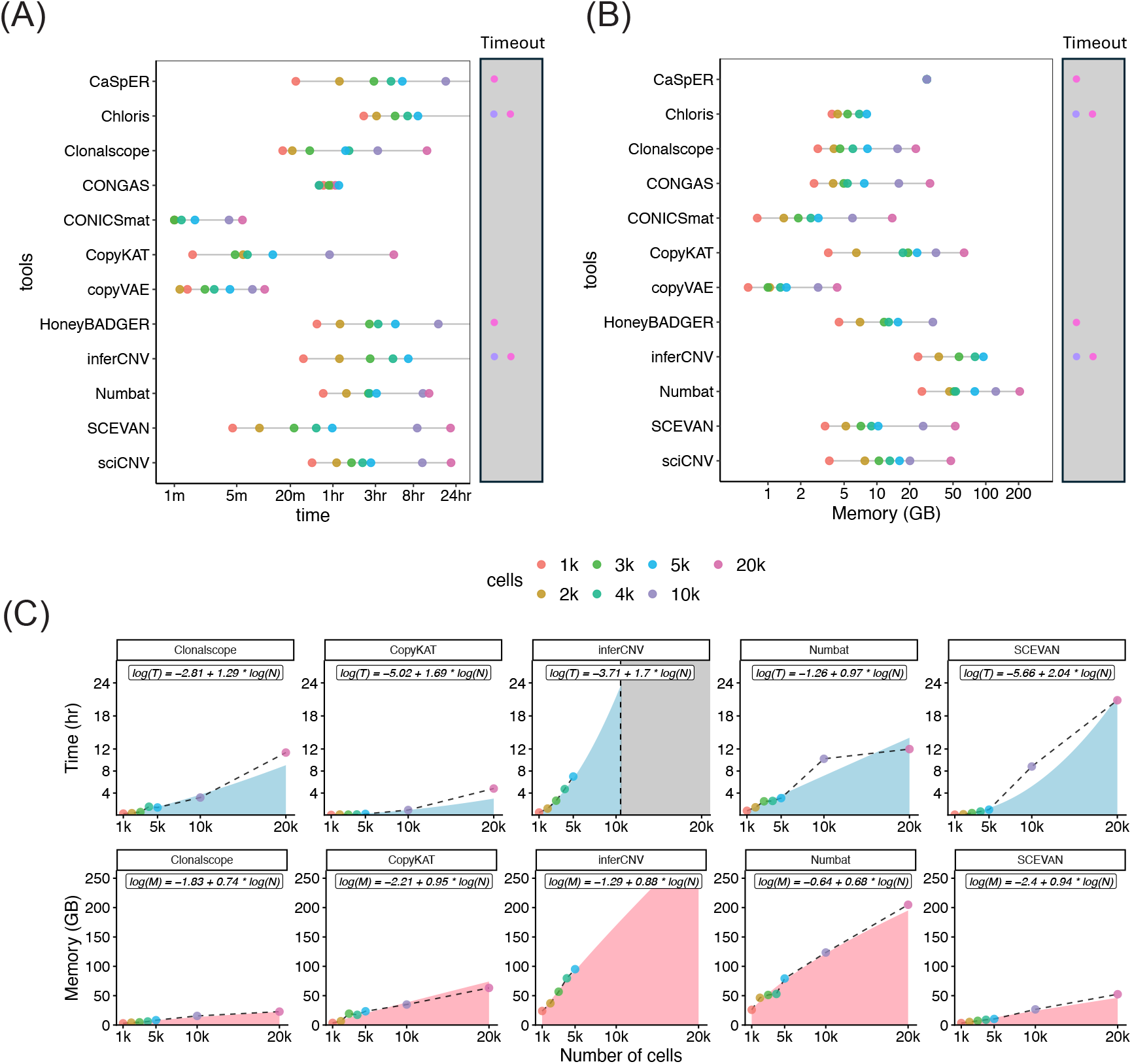
Runtime and memory benchmark. (A) Computing time benchmark and (B) memory usage benchmark for CNV calling algorithms across increasing number of cells. Running time is right censored at 24 hours, respectively. (C) Computing time for the five top-performing tools. Dash lines represent the observed running time/memory usage, and the curve areas indicate predicted values from log-log regression models. Figures are plotted in the original scale. *Note: For each setting, 10 replicates were generated if the running time was less than 30 minutes; otherwise, 3 replicates were generated. The median value is reported in the figure*.

Our analysis revealed significant variability in scalability across tools (Fig. 4A-B). While CONGAS and CaSpER maintained nearly constant runtime and memory footprints, respectively, most methods exhibited predictably increasing demands as datasets grew. Notably, Chloris, HoneyBADGER, inferCNV, and CaSpER failed to complete within the 24-hour window for the 20k cell dataset. Chloris and inferCNV were the least scalable, failing at the 10k cell threshold.

The scaling of runtime efficiency—represented by the slopes of our log—log models-varied widely among the top five performing methods (Fig. 4C). SCEVAN proved the least scalable, with a runtime slope of ::2.04, indicating a steep quadratic increase. In contrast, Clonalscope (slope = 1.29) and Numbat (slope = 0.97) demonstrated superior scalability, with Numbat achieving approximately linear growth. CopyKAT and inferCNV fell between these extremes, with slopes of 1.69 and 1.70, respectively. Memory usage for all five top-performing tools scaled more efficiently than runtime, exhibiting sub-linear to linear growth (slopes = 0.68-0.95). We also assessed the usability of each method using standardized criteria previously established for evaluating single-cell data integration tools^32^ (see Methods and Supplementary Fig. 45). Overall, most tools demonstrated high usability, supported by well-maintained open-source repositories and clear tutorials.

### B4: Robustness and Sensitivity Analysis

In practical applications, several technical factors—including CNV-state thresholds, sequencing depth, the inclusion of intronic reads, and reference panel selection—can significantly influence inference performance. We systematically evaluated the sensitivity of all 12 tools to these parameters to determine their robustness in real-world workflows, with the five top-performing methods from B2 evaluation presented in Fig. 5.

**Figure 5:**
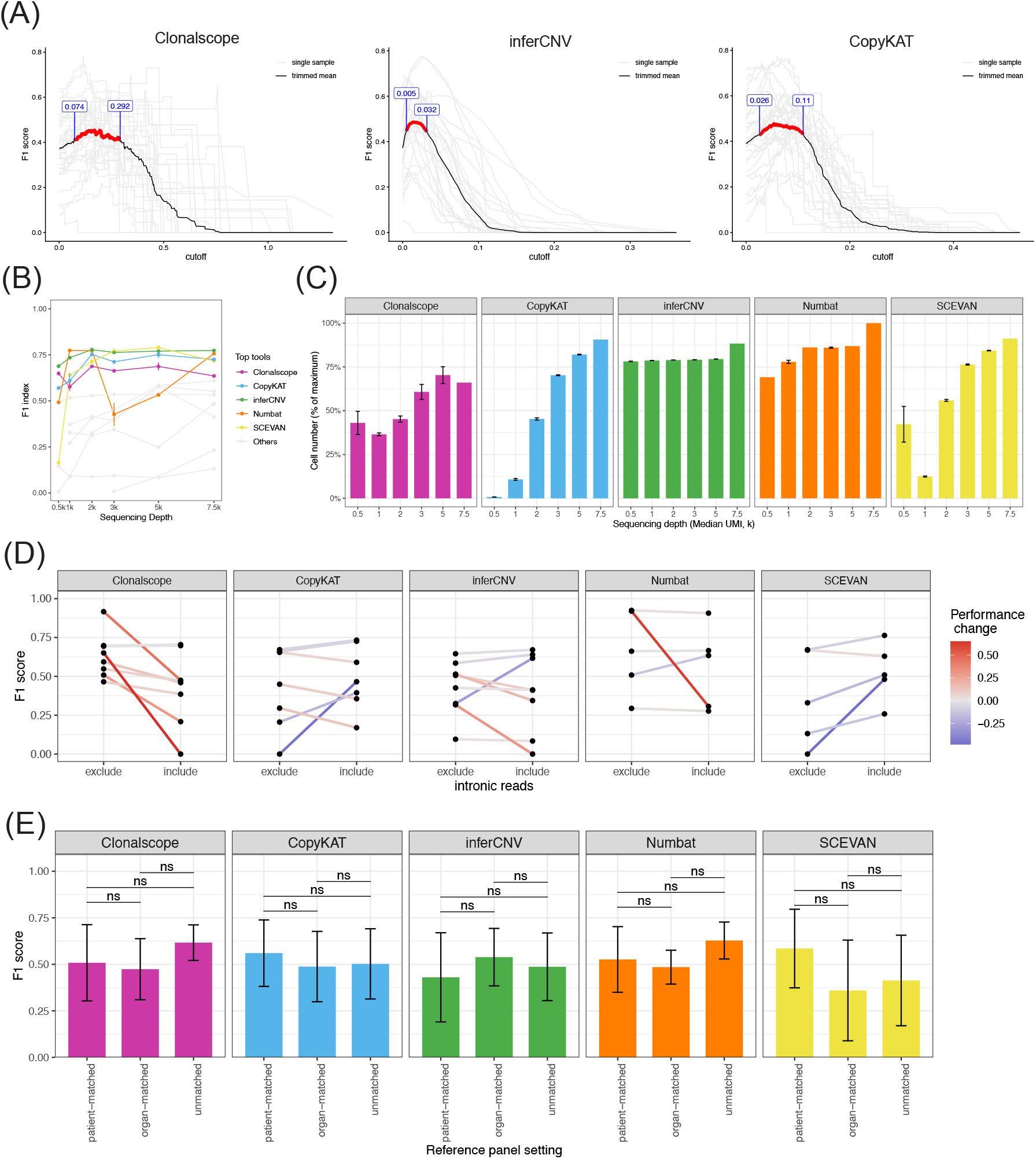
Robustness and sensitivity analysis. (A) CNV-state cutoff of three top-performing intensity-based approaches. Red curves achieve ≥ 90% of the maximum trimmed mean Fl score. (B) Effect of sequencing depth on CNV inference performance. (C) The number of retained cells under different sequencing depth after quality filtering. (D) Impact of intronic-read inclusion on CNV inference performance (E) Robustness of CNV inference accuracy under different reference-panel choices.

**Figure 6:**
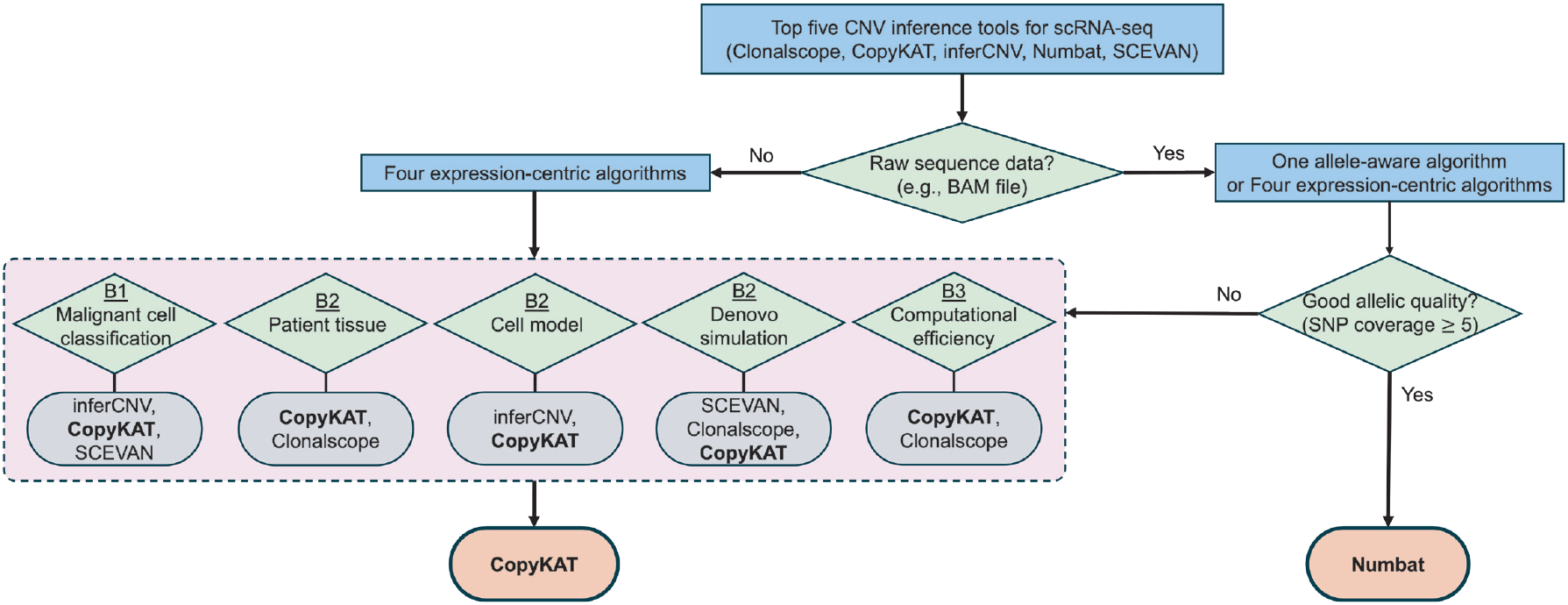
Tree-based guideline for users. Two methods (CopyKAT and Numbat) are recommended for different scenarios in the tree-based guideline.

For intensity-based methods among the top five (CopyKAT, Clonalscope, and inferCNV), we found that the performance of CopyKAT and Clonalscope was more robust to the choice of CNV-state cutoff than that of inferCNV (Fig. 5A). In contrast, several other tools, such as sciCNV and CaSpER, exhibited high sensitivity to these thresholds (Supplementary Fig. 46).

Regarding sequencing depth, most top-performing methods tolerate moderate library-size reductions. However, performance degrades sharply under extremely low-depth conditions (Fig. 5B). Furthermore, as library size decreases, the number of cells surviving standard quality-control filters drops substantially, which can severely limit the resolution of downstream clonal analysis (Fig. 5C).

While recent updates to the 10x Genomics pipeline (Cell Ranger v7.0+) recommend including intronic reads to increase usable data (22% increase of mapped reads; Supplementary Table 2), we found this had a negligible effect on most tools. The notable exception was Clonalscope, which exhibited a significant decline in performance when intronic reads were included (median F1 decrease = 0.16, Wilcoxon P-value < 0.039; Fig. 5D).

Finally, we assessed the impact of reference panel selection. Although patient-matched normal references are the gold standard for minimizing subject-specific biases, they are frequently unavailable. We evaluated five tumor samples under three scenarios: patient-matched, organ-matched, and unmatched (external) references. Across all top-performing methods, we observed no significant difference in F1 scores, suggesting that these tools are sufficiently robust to leverage external reference panels without compromising accuracy (Fig. SE).

### Practitioner’s Guide: A Decision-Making Framework

To translate our benchmarking results into actionable insights, we established a tree-based decision framework (Fig. 6) tailored to common experimental scenarios. While five tools (Clonalscope, CopyKAT, inferCNV, Numbat, and SCEVAN) consistently emerged as top performers, the optimal selection depends on data modality and specific analytical objectives.

A key finding of our study is the performance variability of Numbat; it often outperforms all other methods by large margin but significantly underperforms in the other occasions (Supplementary Fig. 4). Unlike expression-centric tools, Numbat integrates gene expression with B-allele frequency (BAF), providing a significant advantage when allelic signal quality is high. Our analysis of 28 datasets confirms that samples with higher SNP-level depth yielded significantly higher normalized F1 scores (median difference = 0.2, Wilcoxon P-value <0.01; Supplementary Fig. 43A-B). We therefore recommend Numbat as the primary choice when raw sequencing data (e.g., BAM files) are available and SNP coverage is sufficiently high (≥5). However, if SNP-level sequencing depth is insufficient (< 5), CopyKAT serves as the most reliable alternative, offering superior stability under low-coverage conditions (Fig. 6; Supplementary Fig. 43C).

In many real-world scenarios, researchers are limited to processed gene expression matrices due to privacy regulations, data storage constraints, or limited computational resources for processing raw FASTQ/BAM files. In these cases, only expression-centric methods are viable, and their strengths vary by task: (B1) Malignant Cell Classification: inferCNV, CopyKAT, and SCEVAN demonstrated the most consistent performance; (B2) CNV Inference Accuracy: CopyKAT and Clonalscope stand out across patient tissue samples, while inferCNV and CopyKAT outperform in cell model samples (supplementary Fig. 5). In additional, de novo simulations show that SCEVAN, Clonalscope, and CopyKAT achieved higher F1 scores and ARI values; (B3) Computational Efficiency: CopyKAT and Clonalscope are the most scalable options, requiring significantly less runtime and memory, which is critical for large-scale atlas projects. Taking all aspects together, CopyKAT provides the best overall balance of accuracy, interpretability, and efficiency when raw sequencing data are unavailable (Fig. 6). Once a tool is selected, users can further refine their results using our robustness analysis (B4) to optimize parameters such as CNV-state cutoffs, normal diploid reference selection, and the inclusion of intronic reads.

## Discussion

We have presented the most extensive benchmarking analysis of scRNA-seq CNV inference to date, evaluating 12 computational tools across 28 diverse tumor datasets with over 100,000 cells. By systematically assessing four benchmarks - malignant cell classification, CNV inference accuracy, scalability, and robustness (B1-B4) -, our study provides a comprehensive framework for conclusively navigating the current landscape of single-cell transcriptomics. To bridge the gap between algorithmic development and biological application, we translated these findings into a practical, decision-tree-based recommendation system (Fig. 6) that accounts for real-world data constraints encountered by practitioners.

Our results revealed substantial variability in tool performance across datasets and experimental conditions, yet several key principles emerged. First, the allele-aware method, Numbat, frequently outperformed expression-centric tools when allelic inference quality was high (≥5), as reflected by SNP-level depth. However, its performance dramatically decreases when averaging sequencing depth on SNPs is low (< 5). Second, the expression-centric method, CopyKAT, exhibited efficient runtime and modest memory requirements, allowing it to run on a standard personal computer, while maintaining robust performance across datasets. Third, we found that copy-state-based methods (e.g., inferCNV, Numbat, SCEVAN) provide a more rigorous and interpretable alternative to intensity-based methods, as they mitigate the biases associated with arbitrary threshold selection and offer discrete, biologically meaningful outputs. However, intensity-based outputs retain continuous signal information, which can be useful for assessing relative CNV signal strength and detecting subtle copy-number changes. Therefore, the choice between these two frameworks depends on whether interpretability or quantitative signal resolution is prioritized.

Despite these strengths, certain inherent limitations merit consideration. To begin with, although we utilized scDNA-seq and bulk WGS as “gold standards”, these modalities do not profile the exact same cells as scRNA-seq, potentially introducing modality-specific subclonal discrepancies. While the advanced de novo simulations provided an absolute ground truth to mitigate this, no simulation can fully encapsulate the longitudinally dynamic and complex transcriptional system. Beyond this, our study focused primarily on droplet-based scRNA-seq; while these represent the vast majority of current datasets, the performance of these tools on alternative platforms (e.g., plate-based methods) warrants further investigation. Furthermore, we observed suboptimal performance across methods in subclonal reconstruction, which may stem from both algorithmic constraints in clustering approaches and imperfectness in the ground truth used to benchmark subclonal structure. Future work could focus on improving clustering approaches and developing more reliable gold-standard references. Finally, while our 100,000-cell cohort is the largest to date, the expanding landscape of rare cancer types and emerging multi-omic technologies will continue to provide new data for future evaluations.

As the field of single-cell CNV analysis develops rapidly, we envision two major future directions, in which our comprehensive evaluation and decision-making framework will provide a solid foundation for method development and comparison. (1) Pan-cancer and multi-sample learning: Much like the recent advancements in bulk RNA-seq using large-scale pan-cancer data (e.g., deep-learning-based RCANE^33^), scRNA-seq CNV inference is poised to move toward joint-learning models. By training on vast, pan-cancer and cross-institutional cohorts, future algorithms could learn tissue-specific expression baselines and recurrent subclonal signatures, significantly improving the signal-to-noise ratio in challenging primary tumor samples. (2) Spatial integration: The rise of spatial transcriptomics presents an unprecedented opportunity for CNV inference. Incorporating spatial coordinates can serve as a powerful prior, as cells in close physical proximity are statistically more likely to share clonal origins. Spatial-aware modeling will be essential for accurately mapping the physical architecture of tumor evolution.

In summary, this study provides a foundational resource for the single-cell community, offering clear guidance on preprocessing, reference selection, and tool optimization. By providing our reproducible pipelines and predicted CNV profiles through our public repository, we aim to facilitate the standardization of CNV analysis and provide a rigorous baseline for the next generation of computational tool development.

## Data Availability

For the eight gastric primary and metastatic samples, scRNA-seq data were obtained from dbGaP under accession phs001818, and scDNA-seq data were obtained from dbGaP under accessions phs001711 and phs001818. For the nine gastric cancer cell lines, scDNA-seq and scRNA-seq data were downloaded from the SRA repository under accessions PRJNA498809 and PRJNAS98203, respectively. Two breast cancer bone metastasis samples (BoML and BoMR^34^) and eight breast cancer PDO samples were generated in the Lee-Oesterreich lab at the University of Pittsburgh (Under submission to dbGAP, contact AVL for resource). For samples without patient-matched normal references, two external normal references were used. A 2000-cells normal breast dataset was subset from the Human Breast Cell Atlas (HBCA)^35^, which were accessed through the CZ CELLxGENE Discover Census (version 202S-04-20) for breast cancer bone metastasis and PDO samples. The rectum, marrow, and stomach datasets^36^ were downloaded from GEO (under accession number GSE159929; the rectum dataset GSM48S0S86, the marrow dataset GSM4909954, and the stomach dataset GSM4909960) and used for the analysis of primary CRC P8823, primary multiple myeloma MM199, and gastric cancer cell lines, respectively.

## Acknowledgement

This work is supported in part by NIH Award Number R01CA285337 (HCC, VS, HC, SW, RS, SO, AL and GCT), R01LM014142 (GCT), R01CA252378 (SO, AVL), Susan G. Komen award SAB220213 (AVL, SO) and Breast Cancer Research Fund award BCRF SPEC-22-014 (SO and AVL). This study used the University of Pittsburgh Center for Research Computing and Data, RRID:SCR_022735, through the resources provided. Specifically, this work used the HTC cluster, which is supported by NIH award number S10OD028483.

## Methods

### Preprocessing the single-cell RNA-Seq datasets

We collected 28 scRNA-seq datasets from previous studies and in-house experiments. All datasets were preprocessed using Cell Ranger (v9.0.1) with default parameters to generate gene—cell count matrices and BAM files from FASTQ inputs. Because Cell Ranger v9.0.1 includes intronic reads by default, we additionally generated count matrices excluding intronic reads for robustness analysis by setting include-introns as false. For datasets where only BAM files were available (e.g., P5915 and P6335), FASTQ files were recovered using the bamtofastq tool provided by Cell Ranger. For breast cancer bone metastasis and PDO samples processed with sample multiplexing, we demultiplexed hashed samples by extracting Antibody Capture counts and classifying cells using HTO signals through Seurat’s HTODemux workflow.

In addition to expression matrices and BAM files, cell-type annotations were essential for (1) preparing Numbat input, (2) identifying internal normal reference cells, and (3) validating the ground truth malignant cells. Inspired by GPTCelltype^37^, we performed automated annotation using a combined Seurat and ChatGPT strategy. We first clustered cells and identified the top 10 marker genes per cluster, then used then used ChatGPT to assign cell-type labels. Notably, we did not apply standard cell-filtering steps (e.g., low UMI count, high mitochondrial fraction) because our analysis required retaining all cells when validating ML-based malignant-cell predictions. Interestingly, ChatGPT reliably annotated clusters enriched for low-count or high-mitochondrial cells as low-quality populations, demonstrating robustness even in the presence of problematic cells.

### Generating ground-truth malignant cells

To fairly benchmark malignant cell classification, we defined ground-truth malignant cells using three recent ML-based classifiers (scMalignantFinder^20^ (v1.1.7), ikarus^21^ (v0.0.3), and scATOMIC^22^ (v2.0.3)), which do not rely on CNV information. This choice avoids a possible bias in the comparison. Because these methods differ in model design and decision boundaries, we adopted a majority-vote strategy: a cell was labeled as malignant if at least two of the three classifiers predicted it as malignant. All cells from cancer cell line datasets were treated as malignant by default. We applied a relaxed threshold (tumor probability ≥ 0.3) for ikarus^21^, as we observed that ikarus tended to be more conservative than the other two methods.

### Generating and Verifying ground-truth CNV from single-cell and bulk DNA

Copy number variation profiles derived from single-cell and bulk DNA-seq were used as ground truth. For single-cell DNA-seq, we generated per-cell BAM files using subset-bam (v1.1.0) and performed preprocessing, quality control, and copy number inference with CopyKit (v0.1.2). For bulk DNA, we used CNVkit (v0.9.12) to estimate copy number profiles. Both CopyKit and CNVkit employ the circular binary segmentation (CBS) algorithm and estimate sample ploidy. The estimated ploidy for cell line samples from CopyKit was consistent with original publication^27^, which further strengthens our confidence in the ground-truth copy number profiles. For bulk DNA, copy number states (gain and loss) were assigned using the default threshold (0.5) recommended in the CNVkit documentation

### Postprocessing of CNV inference output

The resolution and output format of predicted CNVs vary across tools. For resolution, some methods report CNVs at the gene level, while others output CNVs at the bin, segment or chromosome-arm level (e.g., copyVAE, Clonalscope, and CONGAS). Additionally, different tools produce either discrete CNV states (e.g., copy-state-based methods) or continuous CNV intensity scores (e.g., intensity-based methods). To enable consistent comparison, we standardized all outputs to gene-level discrete CNV states. First, CNVs reported at the segment, bin, or chromosome-arm level were mapped to gene coordinates. Second, for methods producing continuous CNV intensity scores, we converted these values into discrete CNV states (amplification, neutral, and deletion) using a semi-supervised cutoff based on the global proportion of CNV events inferred from matched DNA-seq ground truth.

### CNV inference methods

All 12 selected CNV inference methods were executed using their default parameter settings, following the recommendations from available tutorials, original publications, or communication with the authors. Detailed descriptions of methods and any adjustments made to address software-related issues are provided in the Supplementary Methods.

### De novo simulations

To generate synthetic datasets with known CNV profiles and subclonal structures, we performed de novo simulations using scDesign3^31^. The simulation procedure consisted of three steps: (1) learning the distribution of gene expression from diploid scRNA-seq data, (2) imposing artificial CNV effects on gene expression means, and (3) generating synthetic cells from the modified generative model.

First, we used a diploid reference scRNA-Seq dataset from normal tissue of patient P5931 (995 cells) to estimate the baseline expression distribution. Gene expression for each gene was modeled using a negative binomial distribution, and gene-gene dependence was captured using a Gaussian copula framework implemented in scDesign3. Second, CNV effects were introduced based on the subclonal CNV structure derived from scDNA sequencing of the SNU638 gastric cancer cell line. Three subclones were defined from the scDNA CNV profile. Synthetic cells were assigned to these subclones with proportions of 30%, 40%, and 30%. For genes located in amplified or deleted regions (copy-number ratio >1.25 or <0.75), the mean expression parameter was adjusted according to the corresponding CNV dosage effect. To mimic varying signal strengths, we simulated six levels of CNV effect size ranging from weak to strong. Finally, synthetic scRNA-seq data were generated using the modified mean parameters together with the remaining parameters learned from the reference dataset. This framework produced datasets with realistic gene-expression distributions while preserving predefined CNV structures and subclone compositions.

All of our data simulation scripts are publicly available as R scripts at github.com/hung-ching-chang/scCNVBench. Additional implementation details of the simulation procedure are provided in the Supplementary Methods.

### Benchmark metrics

For cell-level or subclone-level CNV accuracy evaluation, we assign each cell or subclone to the most similar ground-truth subclone derived from scDNA-seq and calculate the Youden index to evaluate performance, as it jointly considers sensitivity and specificity.

To comprehensively evaluate the performance of the tested tools, the main benchmark metrics were concordance score, event-level F1 score, event-level Youden index, subclone recovery score, trimmed mean F1 score, accuracy, sensitivity, specificity, and ARI. The definitions of these metrics are provided below.

#### Prediction concordance score

The concordance score was designed to quantify the consistency of malignant-cell predictions across CNV-based methods as well as ML-based classifiers. Because the ground-truth malignant-cell labels were defined using ML-based classifiers that do not explicitly incorporate CNV information, agreement among CNV-based methods provides an additional evaluation of prediction reliability.

Let *M*_*i*_ and *M*_*j*_ denote the malignant cells predicted by method *i* and *j*. The pairwise concordance between two methods was measured using the Jaccard index, defined as

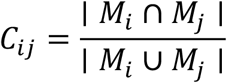

where | *M*_*i*_ ∩ *M*_*j*_ | represents the number of cells predicted as malignant by both methods and | *M*_*i*_ ∪ *M*_*j*_ | represents the total number of cells predicted as malignant by either method. Because not all methods produce reliable predictions, we defined the concordance score of method *i* as the average of the three highest pairwise concordance values between method *i* and the others:

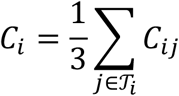

where 𝒯_*i*_ denotes the set of five methods with the highest concordance values with method The concordance score ranges from 0 to 1, where higher values indicate stronger agreement with other high-performing methods.

#### Event-level F1 score

The event-level F1 score was proposed to quantify how accurately a method detects CNV events (amplifications and deletions). Specifically, for gene-based CNV, each gene was assigned one of three states: deletion (-1), neutral (0), or amplification (+1). CNV events were defined as non-neutral states (-1 or +1), and a gene was counted as correctly predicted only when its predicted state exactly matched the ground truth. Let *y* and *y* denote the ground-truth and predicted CNV states for gene *g*, respectively. The true positive, true negative, false positive, and false negative are defined as follows: *TP =* ∑_*g*_ 1 (*y* = 1 Λ *ŷ*_*g*_ *=* 1) + ∑_*g*_ 1 (*y*_*g*_ *=* − 1 Λ *ŷ*_*g*_ *=* − 1), *FP =* ∑_*g*_ 1 (*y*_*g*_ ≠ 1 Λ *ŷ*_*g*_ *=* 1) + ∑_*g*_ 1 (*y*_*g*_ ≠ 1 Λ *ŷ*_*g*_ *=* − 1), and *FN =* ∑_*g*_ 1 (*y*_*g*_ ≠ 0 Λ *ŷ*_*g*_ *=* 0). Event-level precision and event-level recall were defined as

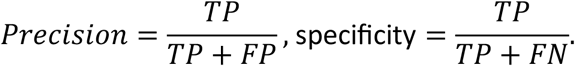

and the event-level F1 score was computed as the harmonic mean of precision and recall given by the equation:

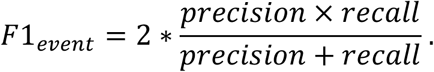

It returns a value between 0 and 1, where 1 indicates perfect recovery of all true CNV events (both amplification and deletion) with no false positives.

#### Event-level Youden index

While the event-level F1 score focuses on the accuracy of detecting CNV events, we additionally evaluated performance using the event-level Youden index, which accounts for both correct detection of CNV events and correct identification of neutral regions. Using the same confusion matrix defined above, sensitivity and specificity were computed as

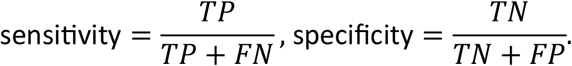

The event-level Vouden index was then defined as *Youden*_*event*_ *=* sensitivity + specificity – 1. The index ranges from − 1 to 1, where 1 indicates perfect classification performance, 0 indicates performance equivalent to random prediction, and negative values indicate systematic misclassification.

#### Subclone recovery score (SRS)

The subclone recovery score quantifies how well predicted subclonal structures correspond to the ground-truth subclones derived from scDNA-seq data. Each predicted subclone was first matched to the ground-truth subclone that achieved the highest CNV similarity, measured using the event-level F1 score. To avoid counting spurious or weakly supported matches, only assignments with event-level F1 score greater than 0.6 were considered valid matches. Let *K*_1_ denote the number of ground-truth subclones successfully identified by at least one predicted subclone, *K*_2_ denote the total number of predicted subclones produced by a method, and *K*_3_ denote the total number of ground-truth subclones. The subclone recovery score was defined as

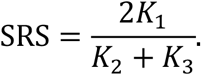

This formulation penalizes both missing subclones (when *K*_1_ *< K*_3_) and over-segmentation (when *K*_2_ *> K*_3_). The score ranges from 0 to 1, where 1 indicates perfect recovery of the ground-truth subclonal structure.

#### Trimmed mean Fl score

To evaluate the robustness of CNV-state cutoff selection, we summarized performance using a trimmed mean of event-level F1 scores across samples. Specifically, for each cutoff value, the top 2S% and bottom 2S% of F1 scores were excluded to reduce the influence of extreme values and outlier samples, and the remaining S0% were averaged. To define a recommended range of cutoff values, we identified those achieving at least 90% of the maximum trimmed mean F1 score, representing configurations that maintain near-optimal and robust performance across samples.

#### Other standard metrics

For malignant cell classification, we also evaluated classification performance using accuracy, sensitivity, and specificity. Given the ground-truth malignant-cell labels defined by ML-based classifiers, accuracy was defined as the proportion of correctly classified cells among all cells, sensitivity as the proportion of malignant cells correctly identified, and specificity as the proportion of normal cells correctly identified.

For evaluating subclonal structure in synthetic datasets generated by *de novo* simulation, where the ground-truth clustering labels are known, we also measured clustering accuracy using the Adjusted Rand Index (ARI), which quantifies the similarity between predicted and true cluster assignments.

To facilitate comparison across methods under varying sample difficulty, we additionally computed a normalized F1 score by scaling each method’s performance relative to the best achievable performance within each sample. This normalization accounts for sample-specific upper bounds in CNV inference accuracy and enables clearer assessment of relative method performance. In particular, it allows us to observe the impact of allelic information on Numbat’s performance compared with other methods.

### Normal reference selection

To reflect the real-world usage, we implemented a hierarchical strategy for normal reference selection at three levels: patient-matched normal (non-tumor tissue from the same patient), organ-matched normal, and unmatched normal (Supplementary Fig. 1). Patient-matched normals were considered the most reliable and were used whenever available (e.g., in gastrointestinal cancer datasets^38^ such as P5846, P5931, P5915). When patient-matched normals were unavailable, such as in cell lines or PDOs (e.g., NCI-N87, SNU601, KATOIII), we used organ-matched external samples as reference. When neither patient-matched nor organ-matched normals were available, we selected the closest available organ as an unmatched reference. For example, for sample P8823 (colon cancer tissue), we used normal rectum tissue as the reference (Supplementary Data 1).

### Scalability assessment

To quantify computational efficiency, we recorded runtime and peak memory usage for each method across datasets with different numbers of cells (from 1k to 20k cells). Runtime (*T*, in hours) and memory usage (*M*, in GB) were measured while running each method using default parameters under the same computational environment. To characterize scalability with increasing dataset size, we modeled runtime and memory usage as power-law functions of the number of cells (*N*). Specifically, we fitted the following log-log linear regression models: log (*T*) *= a*_*T*_+ *b*_*T*_ lo*g* (*N*) and log(*M*) *= a*_*M*_+ *b*_*M*_ (*N*), where *T* and *M* denote runtime and peak memory usage, and *N* denotes the number of cells. The intercept parameters (a_*T*_, a_*M*_) reflect the baseline computational cost, whereas the slope parameters (b_*T*_,b_*M*_) quantify how rapidly runtime and memory requirements increase with dataset size. Larger slope values indicate poorer scalability with increasing numbers of cells.

### Usability assessment

We assessed the usability using an objective scoring system adapted from Luecken et al.^32^. A set of ten categories were defined to comprehensively evaluate the user-friendliness of each method. The first six categories were grouped into a package score, including open-source availability, version control, unit testing, GitHub repository, tutorial availability, and function documentation, which assess code accessibility, software maintenance, and ease of use for new users. The remaining four categories were grouped into a paper score, including peer review status, evaluation of accuracy, evaluation of robustness, and benchmarking against other methods, which assess the level of methodological validation and transparency reported in the original publication. Scores from the ten categories were first averaged within the package and paper groups and then summed to obtain an overall usability score (range 0-2) for each method. Supplementary Data 2 reports the scores, references, and reasons for any lost points.

### Visualization

Inspired by Saelens et al.^39^, we implemented the R package *funkyheatmap* to visualize overall method performance (Fig. 2). Aggregated scores from the B1-B3 tasks are represented by bars to highlight differences in overall performance across methods, whereas individual metric scores are displayed as colored circles or squares. This visualization allows simultaneous comparison of task-level summaries and metric-specific performance. The overall ranking was determined solely based on the B2 task (CNV event inference), which represents the central objective of this study. B1 (malignant-cell classification) was not used for ranking because not all methods provide built-in functionality for malignant-cell identification. B3 (scalability and usability) was also excluded from the ranking because these factors depend largely on available computational resources and user-specific constraints.

To visualize CNV profiles and subclonal structure, we generated genome-wide CNV heatmaps from gene-level CNV estimates. Gene positions were mapped to genomic coordinates and aggregated into equally spaced genomic bins to produce smoothed CNV signals across chromosomes. To illustrate the correspondence between scRNA-derived and scDNA-derived subclones, we implemented ribbon-style plots linking predicted RNA subclones to their matched scDNA subclones. The widths of the blocks represent the relative proportion of cells in each subclone, and curved ribbons connect RNA subclones to the corresponding DNA subclones based on the event-level F1 score. Subclones with insufficient CNV concordance (i.e. below the predefined F1 threshold of 0.6) were excluded from the mapping visualization. This representation provides an intuitive view of both subclone composition and the agreement between RNA- and DNA-derived subclonal structures.

