## Supplemental Methods and Figures for "Benchmarking scRNA-seq Copy Number Inference: A Comprehensive Evaluation and Practitioner’s Guide"

*Supplementary Information*

Hung-Ching Chang[1], Yuxin Shi[1], Haoyu Cheng[2], Jian Zou[1], Alexander Chih-Chieh Chang[3,4,5], Brent T. Schlegel[3,4,5], Wenjia Wang[1], Daniel D. Brown[5], Fangyuan Chen[3,6], Sarah Wang[1], Danyang Li[1], Ria Sai[7], Noelle Michel[3], Steffi Oesterreich[3,4], Adrian V. Lee[3,4], George C. Tseng[1]

[1]Department of Biostatistics, University of Pittsburgh Graduate School of Public Health, Pittsburgh, PA, USA.

[2]Department of Computational Biology, Carnegie Mellon University School of Computer Science, Pittsburgh, PA, USA.

[3]Women's Cancer Research Center, UPMC Hillman Cancer Center, Magee-Womens Research Institute, Pittsburgh, PA, USA.

[4]Department of Pharmacology and Chemical Biology, University of Pittsburgh, Pittsburgh, PA, USA.

[5]Institute for Precision Medicine, University of Pittsburgh, Pittsburgh, PA, USA.

[6]Dana Farber Cancer Institute, Boston, MA, USA.

[7]Department of Chemistry, University of Pittsburgh School of Arts and Sciences, Pittsburgh, PA, USA.

#### SUPPLEMENTARY METHODS 4

**De *novo* simulation of CNV-affected scRNA-seq data 4**

**CNV inference methods 7**
- CaSpER 7
- Chloris 7
- Clonalscope 7
- CONGAS 7
- CONICSmat 7
- CopyKAT 8
- copyVAE 8
- HoneyBADGER 8
- inferCNV 8
- Numbat 8
- SCEVAN 9
- sciCNV 9

#### SUPPLEMENTARY FIGURES 10

**Benchmarking workflow 10**

**Malignant cell classification results 11**
- Sample-wise concordance score 11
- Accuracy, Sensitivity, and Specificity of B1 task 12

**CNV inference results 13**
- Sample-wise F1 score 13
- Patient tissues & cell models 14
- Youden index of B2 task 15
- Sensitivity of CNV events 16
- Subclone-level SRS 17

**Primary sample results 18**
- P5846 18
- P5847 19
- P5931 20
- P8823 21
- MM199 22

**Metastases results 23**
- P5915 23
- P6198 24
- P6335 25
- P6593 26
- BoML 27
- BoMR 28

**Cell lines results 29**
- HGC-27 29
- KATOIII 30
- MKN-45 31
- NCI-N87 32
- NUGC-4 33
- SNU-16 34
- SNU-601 35

SNU-638 36
SNU-668 37

**PDOs results** **38**
IPM-BO-46-early 38
IPM-BO-46-late 39
IPM-BO-53-early 40
IPM-BO-53-late 41
IPM-BO-86-early 42
IPM-BO-86-late 43
IPM-BO-139-early 44
IPM-BO-139-late 45

**De novo simulation results** **46**
Ground-truth CNV profile and $Cg$ distribution 46
Clonalscope 47
CopyKAT 48
inferCNV 49
Numbat 50
SCEVAN 51

**Association between SNP-level depth and CNV inference performance** **52**

**Scalability** **53**

**Usability** **54**

**Robustness** **55**
CNV-state cutoff 55

**Validation of ground-truth CNV profiles** **56**
Cross-modality concordance 56
Literature validation of recurrent CNV events 57

**SUPPLEMENTARY TABLE 1: STUDIES COMPARISON** **58**

**SUPPLEMENTARY TABLE 2: INTRONS INCLUSION AND EXCLUSION** **59**

**SUPPLEMENTARY REFERENCES** **60**

### Supplementary Methods

#### De *novo* simulation of CNV-affected scRNA-seq data

Synthetic scRNA-seq datasets with predefined CNV structures were generated using a three-step procedure: (1) learning the expression distribution of diploid cells, (2) imposing artificial CNV effects on gene-expression means, and (3) generating synthetic expression data using the modified parameters. Steps (1) and (3) were implemented using the simulator scDesign3[1], while step (2) was designed to introduce CNV effects with controlled signal strength.

**<u>Step 1: Learning the baseline expression distribution.</u>** To model diploid gene-expression patterns, we used the normal reference scRNA-seq dataset from patient P5931 (995 cells). All cells were assumed to belong to a single cell type and batch. Using this dataset, scDesign3 first fitted a negative binomial distribution for each gene to model gene-level expression variability. To capture dependencies between genes, scDesign3 further learned the joint gene-expression structure using a Gaussian copula model. These estimated parameters form a generative model that can later produce synthetic gene-expression data with the same marginal distributions and gene-gene dependency structure as the original dataset.

**<u>Step 2: Introducing CNV effects.</u>** To introduce CNV signals, we modified the mean expression parameters learned in **Step 1**. The CNV structure was derived from scDNA-seq of the SNU638 gastric cancer cell line, where three subclones were identified (Supplementary Fig. 36). The raw CNV signals from scDNA-seq, named as CNV dosage $D$, was estimated by CopyKit and calculated by the ratio between the copy number of a gene in the tumor genome and that in a diploid reference genome (baseline is 1). Considering deletion events is much less than amplification events, we expanded the only one deletion segment (on chromosome 14) by double the size of segment (from 461 genes to 856 genes). This step ensured that deletion events were not be oblivious given the focal CNV is still challenge for most of CNV inference methods.

To mimic CNV subclonal structure, the 995 synthetic cells were divided into three groups representing the three subclones (30%, 40%, and 30%; n = 298, 398, and 299 cells, respectively). Each group was assigned the CNV profile of a SNU638 subclone. We defined 0.25 as the effective dosage threshold, such that gene expression was modified only when the CNV dosage exceeded 1.25 (amplification) or was below 0.75 (deletion). Accordingly, the modified mean expression parameter of cell $i$ on gene $g$ was defined as

$$\mu_{ig}^{*} = \begin{cases} \mu_{ig} \times D_{s(i)g} \times C_{ig}, & D_{s(i)g} > 1.25 \ or \ D_{s(i)g} < 0.75 \\ \mu_{ig} \quad , & otherwise \end{cases}$$

where $s(i)$ is the subclone assignment of cell $i$, $\mu_{ig}$is the baseline mean expression, $D_{s(i)g}$ is the CNV dosage, and $C_{ig}$ is the DNA-to-RNA translation strength, describing how strongly DNA copy-number changes influence RNA expression.

To simulate data with varying translation strength $C$, we first estimated the empirical distribution of translation strength using the SNU638 dataset and then sampled $C_{ig}$ with pre-defined parameter for each simulation scenario. For each gene $g$, estimated translation strength was defined as

$$\tilde{C}_g = \frac{R_g}{D_g}$$

where $R_g$ is the averaged RNA expression ratio across cells, and $D_g$ is the averaged DNA CNV dosage across tumor cells. This parameter basically reflects, on average, the magnitude of gene-expression change associated with a unit change in DNA copy number. The RNA expression ratio ($R_g$) was calculated as $\bar{X}_{g,tumor}/\bar{X}_{g,normal}$, where $\bar{X}_{g,tumor}$ and $\bar{X}_{g,normal}$ are the mean gene expression levels with CPM normalization in tumor and normal cells, respectively. Following the same criteria for selecting normal reference cells described in the manuscript, we used normal cells from a normal stomach tissue sample of an adult male (GSM4850590) as the reference for tumor cells from the SNU638 cell line.

Because $\tilde{C}_g$ is a positive ratio ranging from 0 to infinity, we modeled the translation strength $log(1 + \tilde{C}_g)$ using a Gamma distribution. The logarithmic transformation was applied to stabilize the distribution and reduce the influence of extreme values when translation strength is large. To estimate the distribution parameters, we focused on five homogeneous CNV segments shared across the three subclones (located on chromosomes 2, 9, 11, 14, and 17). Maximum likelihood estimation (MLE) was used to fit the following model:

$$log(1 + \tilde{C}_g) \sim Gamma(\hat{\alpha}, \hat{\beta})$$

The estimated $\hat{\alpha}$ values across all segments were close to 1, while the estimated $\hat{\beta}$ values centered around 0.8 but showed substantial variation (Supplementary Fig. 36). Based on these observations, we defined six parameter settings to simulate CNV effects with increasing signal strength:

$$\hat{\alpha} = 1$$

$$\hat{\beta}_{Amp} = 0.4, 0.6, 0.8, 1, 1.2, 1.4$$

$$\hat{\beta}_{Del} = 1.1, 1, 0.9, 0.8, 0.7, 0.6$$

, where $\hat{\beta}_{Amp} = 0.4$ and $\hat{\beta}_{Del} = 1.1$ can generate the $\log(\mu_{ig}^*/\mu_{ig})$ around zero. We note that the empirical estimates $\tilde{C}_g$ were used only to infer the underlying distribution of translation strength. During simulation, the cell-level translation strength $C_{ig}$ for each gene $g$ and cell $i$ was then sampled as

$$log(1 + C_{ig}) \sim \begin{cases} Gamma(1, \hat{\beta}_{Amp}), & if\ D_{ig} > 1\ (Amplification) \\ Gamma(1, \hat{\beta}_{Del}), & if\ D_{ig} < 1\ (Deletion) \end{cases}$$

**<u>Step 3: Generation of synthetic scRNA-seq data.</u>** Synthetic gene-expression matrices were generated using the scDesign3 generative model. For each synthetic cell, scDesign3 first samples a cumulative probability vector from the learned Gaussian copula distribution. These cumulative probabilities are then transformed through the quantile function of the negative binomial distributions fitted in **Step 1** to obtain gene-expression counts. By combining the modified mean parameters from **Step 2** with the learned copula structure and dispersion parameters from **Step 1**, this procedure generates realistic scRNA-seq datasets that preserve gene-expression dependencies while incorporating predefined CNV structures and subclone compositions.

#### CNV inference methods

We ran each method according to defaults provided by the authors and contacted them when errors were encountered. These default parameterizations and corresponding method descriptions are detailed below.

##### CaSpER

CaSpER infers copy number variations from scRNA-seq data by jointly leveraging gene expression signals and B-allele frequency (BAF) information derived from sequencing reads. It applies multi-scale smoothing and integrates expression and allelic signals through a hidden Markov model framework to detect CNV events across genomic regions, improving robustness to noise and enabling identification of both large-scale and focal alterations.

##### Chloris

Chloris infers copy number variations from scRNA-seq data by modeling gene expression along genomic coordinates while accounting for transcriptional variability across cells. It employs a probabilistic segmentation framework that leverages spatial continuity of gene expression to detect CNV regions and reconstruct subclonal structures with improved resolution and robustness.

##### Clonalscope

Clonalscope infers copy number variations from scRNA-seq data by integrating expression signals with genomic segmentation to identify coherent CNV regions. It employs a probabilistic clustering framework that jointly models CNV profiles across cells to reconstruct subclonal structures and improve robustness to noise.

##### CONGAS

CONGAS infers copy number variations from scRNA-seq data by integrating gene expression with prior copy number information derived from bulk DNA sequencing. It employs a Bayesian probabilistic model that jointly infers segment-level CNVs and clusters cells into subclones, leveraging variational inference to capture uncertainty and improve robustness.

##### CONICSmat

CONICS infers copy number variations from scRNA-seq data by integrating gene expression with genomic information derived from orthogonal DNA sequencing data or reference controls. It identifies CNV-associated subclones by quantifying large-scale expression shifts and mapping cells to tumor clones, enabling downstream analyses such as clonal assignment and differential expression.

#### CopyKAT

CopyKAT infers copy number variations from scRNA-seq data by leveraging genome-wide gene expression patterns to distinguish aneuploid tumor cells from diploid normal cells. It uses integrative Bayesian segmentation and clustering to identify large-scale CNV profiles and classify cells into tumor and normal populations, enabling downstream subclonal analysis.

#### copyVAE

copyVAE infers copy number variations from scRNA-seq data using a variational autoencoder that jointly models gene expression and latent CNV states. By learning a low-dimensional representation of genomic bins, it captures large-scale CNV patterns and enables subclonal structure identification through clustering in the latent space.

#### HoneyBADGER

HoneyBADGER infers copy number variations from scRNA-seq data by integrating gene expression with allele-specific expression signals derived from heterozygous SNPs. It employs a Bayesian framework to jointly model expression shifts and allelic imbalance, enabling detection of CNV events and identification of malignant subpopulations.

#### inferCNV

inferCNV infers large-scale copy number variations from scRNA-seq data by comparing gene expression intensity across genomic positions between tumor cells and a set of reference normal cells. It applies smoothing across ordered genes to reduce noise and optionally uses a hidden Markov model (HMM) to assign discrete CNV states and identify subclonal structure.

#### Numbat

Numbat infers copy number variations from scRNA-seq data by jointly modeling gene expression and allele-specific counts at heterozygous SNPs. It uses a probabilistic framework with haplotype-aware phasing to detect CNV events and reconstruct subclonal structure with high resolution.

#### SCEVAN

SCEVAN infers copy number variations from scRNA-seq data using a variational segmentation framework applied to smoothed gene expression profiles along genomic coordinates. It jointly segments multiple cells under the assumption that cells within the same subclone share breakpoints, enabling accurate identification of CNV states, malignant cells, and subclonal structure.

#### sciCNV

sciCNV infers copy number variations from scRNA-seq data by comparing gene expression profiles of tumor cells against a reference set of normal cells, using smoothing across genomic regions to reduce noise. It then classifies CNV states based on relative expression shifts and leverages these profiles to distinguish malignant from normal cells and characterize subclonal structure.

### Supplementary Figures

#### Benchmarking workflow

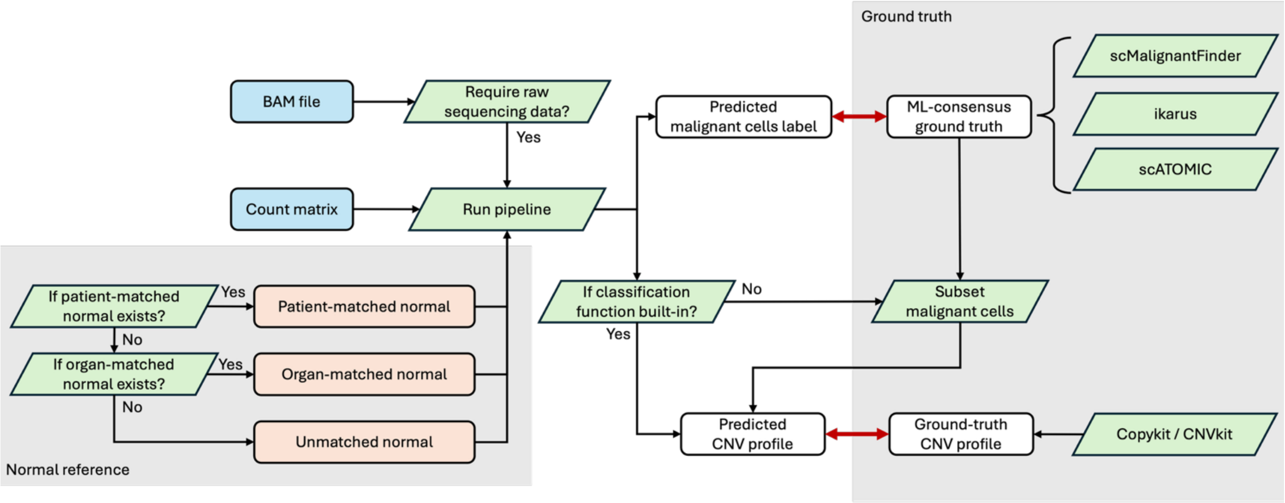


**Supplementary Figure 1: Overview of benchmarking workflow.** Raw sequencing data (e.g., FASTQ and BAM files) are processed to generate count matrix and construct normal reference using a hierarchical strategy (patient-matched, organ-matched, or unmatched). Malignant cell classification performance is assessed by comparison with ML-based consensus ground truth. CNV inference performance is then evaluated against ground-truth CNV profiles.

### Malignant cell classification results

#### Sample-wise concordance score

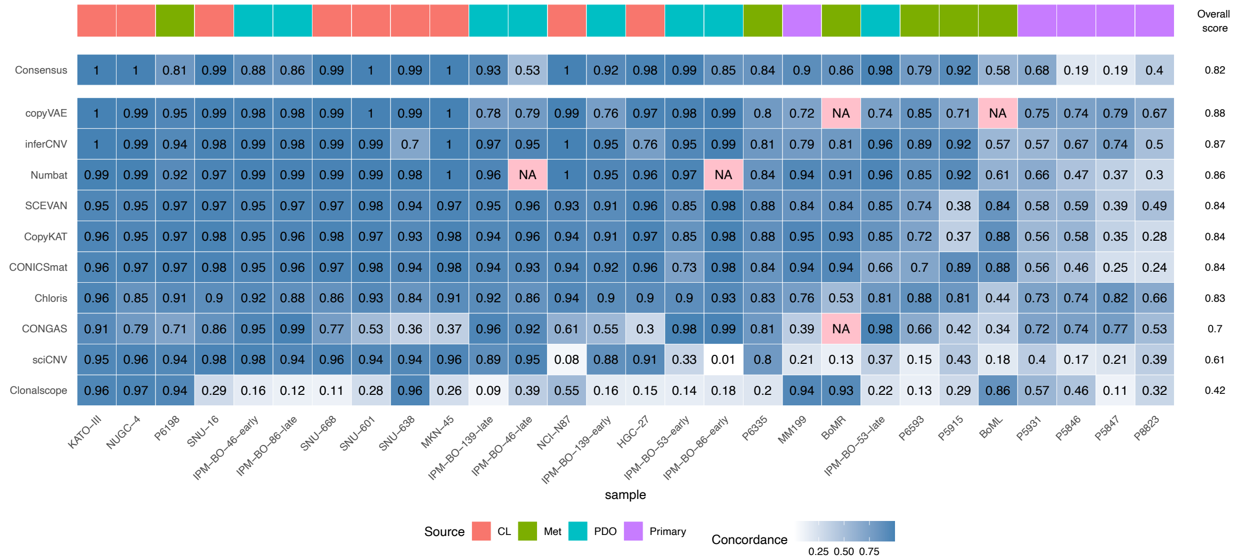


**Supplementary Figure 2: Benchmarking results by sample-wise concordance score for malignant cell classification task.** Concordance was computed as the Jaccard index between CNV profiles inferred by different methods (including scRNA-seq tools and ground truth) for each sample. . Tools are sorted by their overall score, defined as the average concordance across samples. Similarly, samples are sorted by their average concordance (i.e., the average concordance score across methods for each sample), reflecting a spectrum from easier to more challenging cases.

#### Accuracy, Sensitivity, and Specificity of B1 task

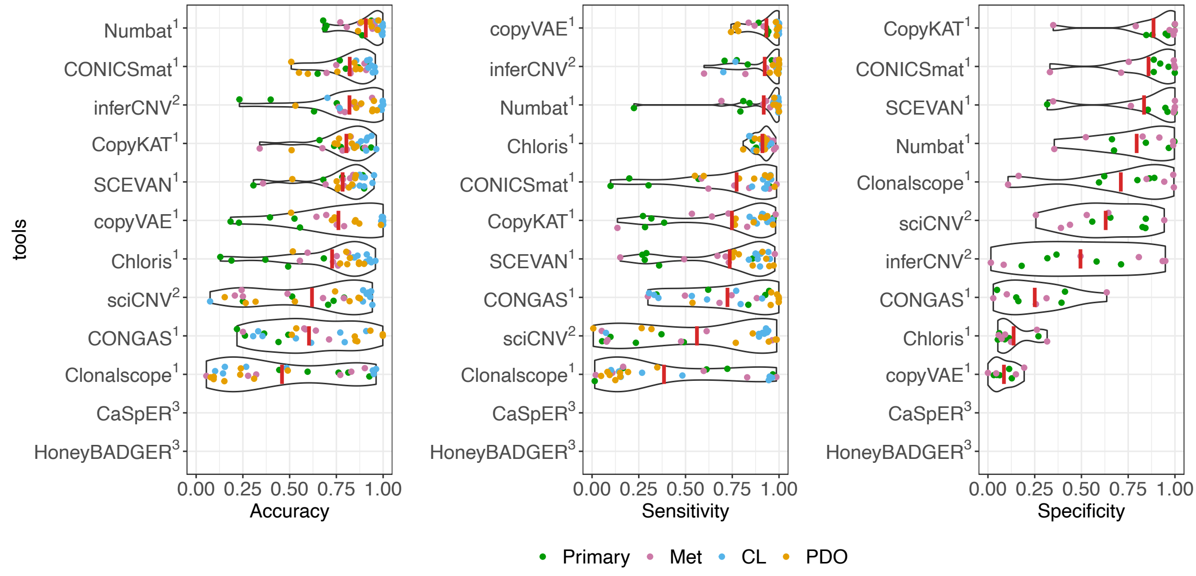


**Supplementary Figure 3: Benchmarking results by accuracy, sensitivity, and specificity for malignant cell classification task.** The plots show overall performance using accuracy, sensitivity, and specificity. Samples are colored by distinct biological sources.

#### CNV inference results

#### Sample-wise F1 score

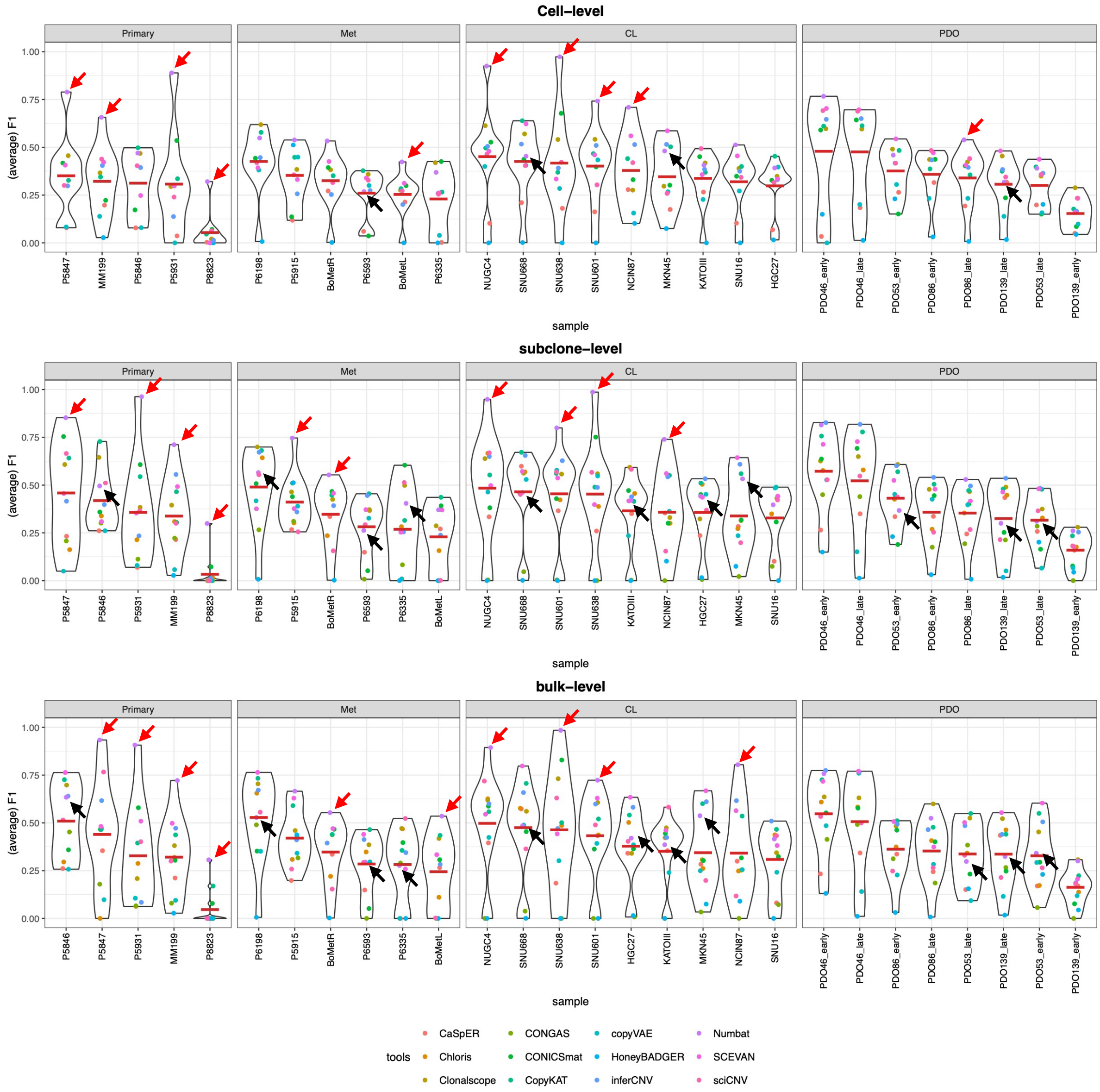


**Supplementary Figure 4: Benchmarking results by sample-wise F1 score for CNV event inference task.** The plots show the accuracy of CNV detection at three resolution levels (cell, subclone and bulk) using event-level F1 score. Samples are devided into four groups (primary, metastasis, cell line, and PDO). Within each group, samples were further sorted by the median F1 score across tools. Methods that failed to run are omitted. The red arrow indicates superior cases where Numbat outperforms other methods (i.e., exceeds the second-best method by at least 0.1 in F1 score), and the black arrow indicates inferior cases where Numbat does not outperform other methods (i.e., falls behind the best methods by at least 0.1 in F1 score).

### Patient tissues & cell models

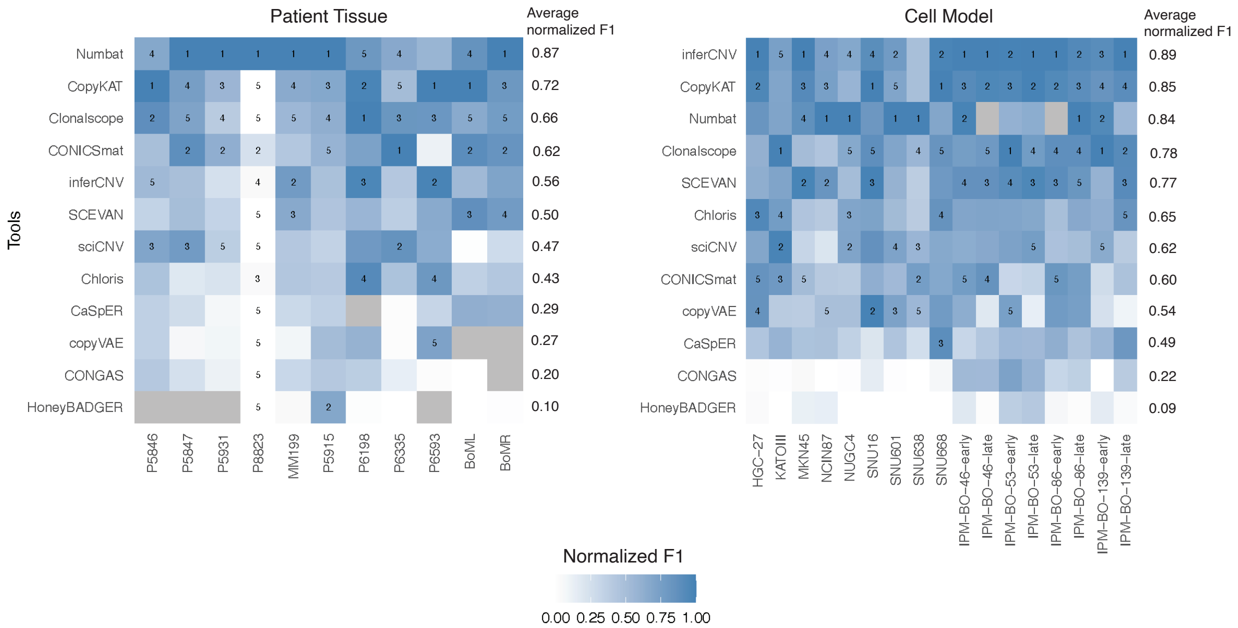


**Supplementary Figure 5: Benchmarking results of patient tissues and cell models by normalized F1 score for CNV event inference task.** Samples are grouped into patient tissues (such as primary samples and metastases) and cell models (such as cell lines and PDOs). The top five methods for each sample are labeled. Methods are then sorted by overall performance.

#### Youden index of B2 task

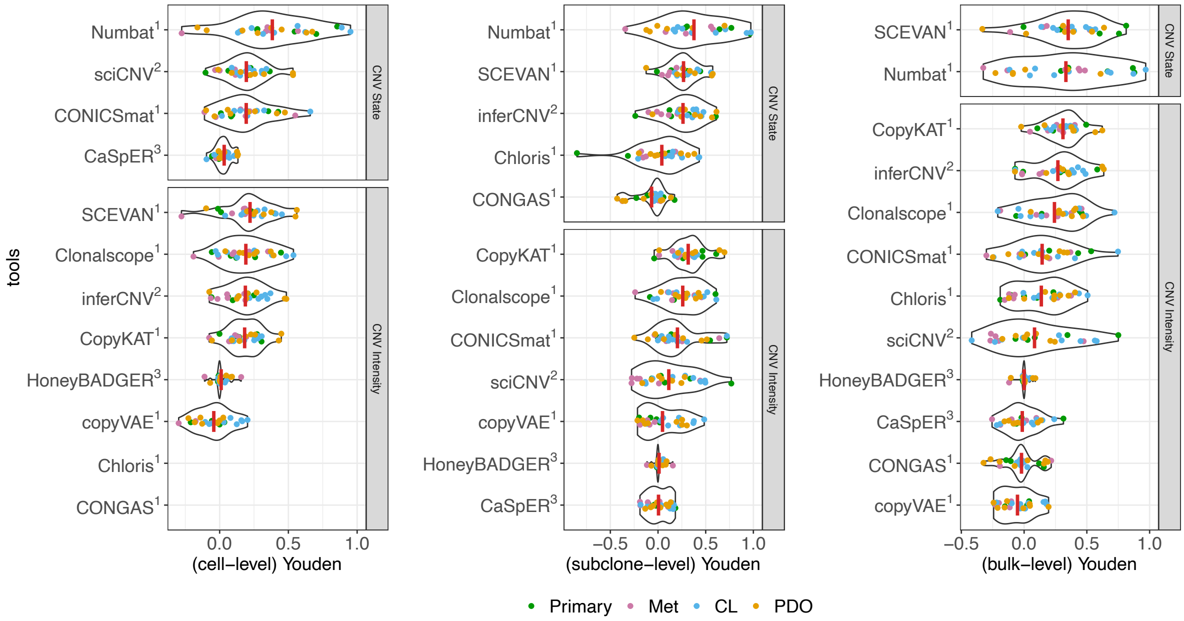


**Supplementary Figure 6: Benchmarking results by event-level Youden index for CNV event inference task.** The plots show overall performance using Youden index. Samples are colored by distinct biological sources.

#### Sensitivity of CNV events

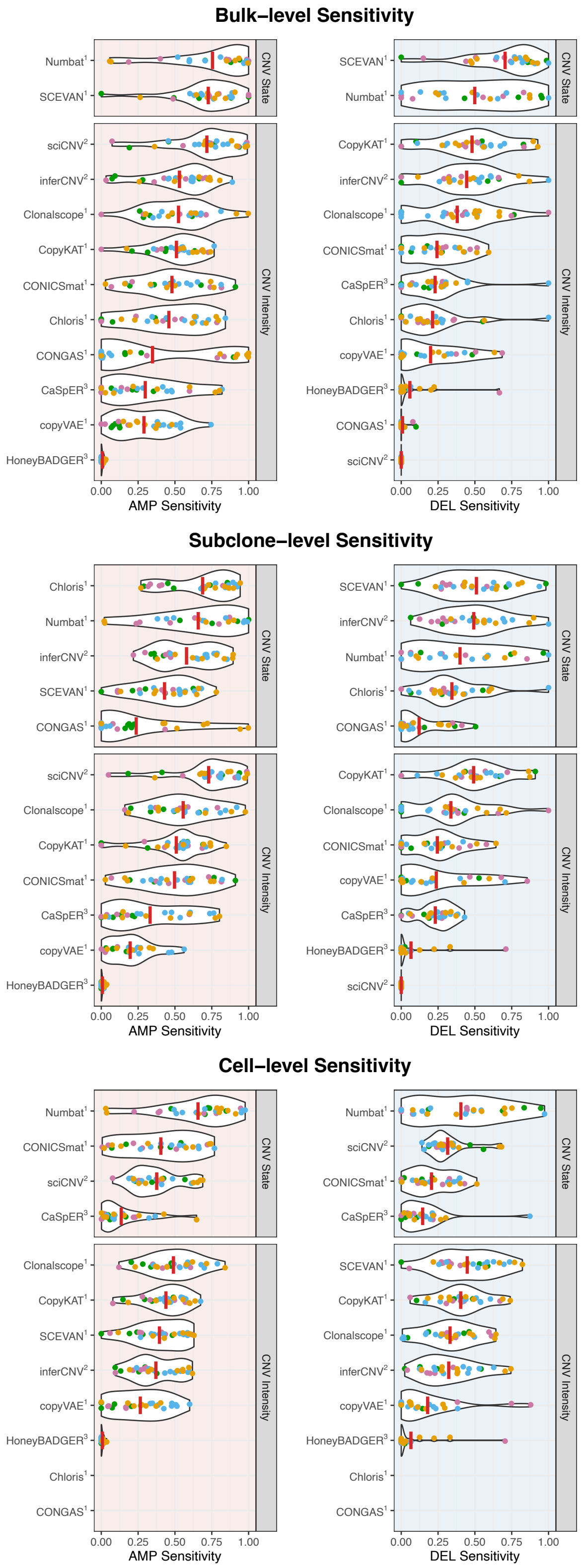


**Supplementary Figure 7: Benchmarking results by event-specific sensitivity for CNV event inference task.** Left panel shows sensitivity of CN amplification, and right panel shows the sensitivity of CN deletion.

#### Subclone-level SRS

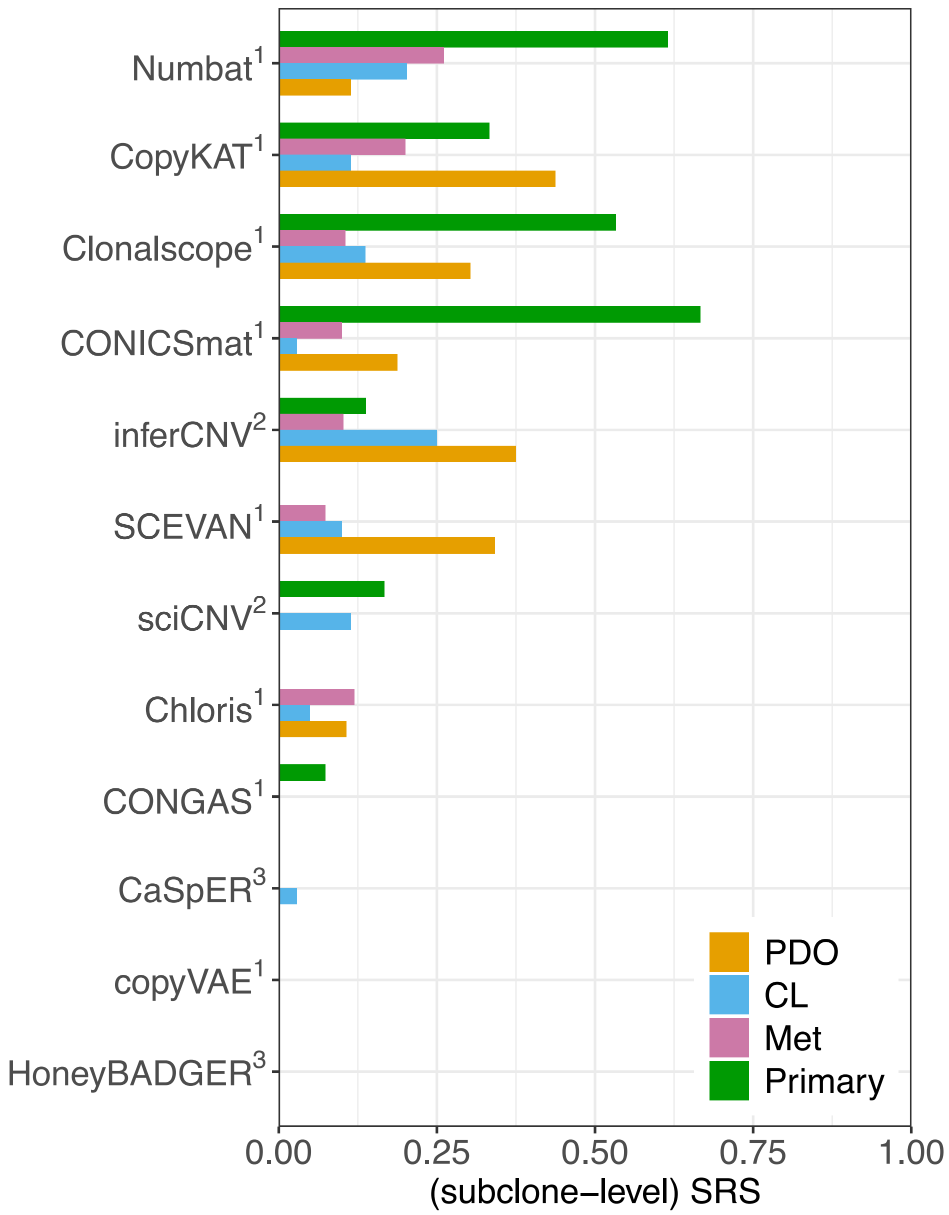


**Supplementary Figure 8: Benchmarking results by cell-level subclone reconstruction score (SRS) for CNV event inference task.** Cell-level SRS evaluates each method's ability to recover ground-truth subclonal structure at single-cell resolution. Compared with subclone-level SRS, higher cell-level scores suggest that CNV signals are preserved at the individual cell level, but clustering fails to accurately group cells into subclones (e.g., CopyKAT). Methods are ordered by subclone-level SRS (see Fig. 3).

### Primary sample results

## P5846

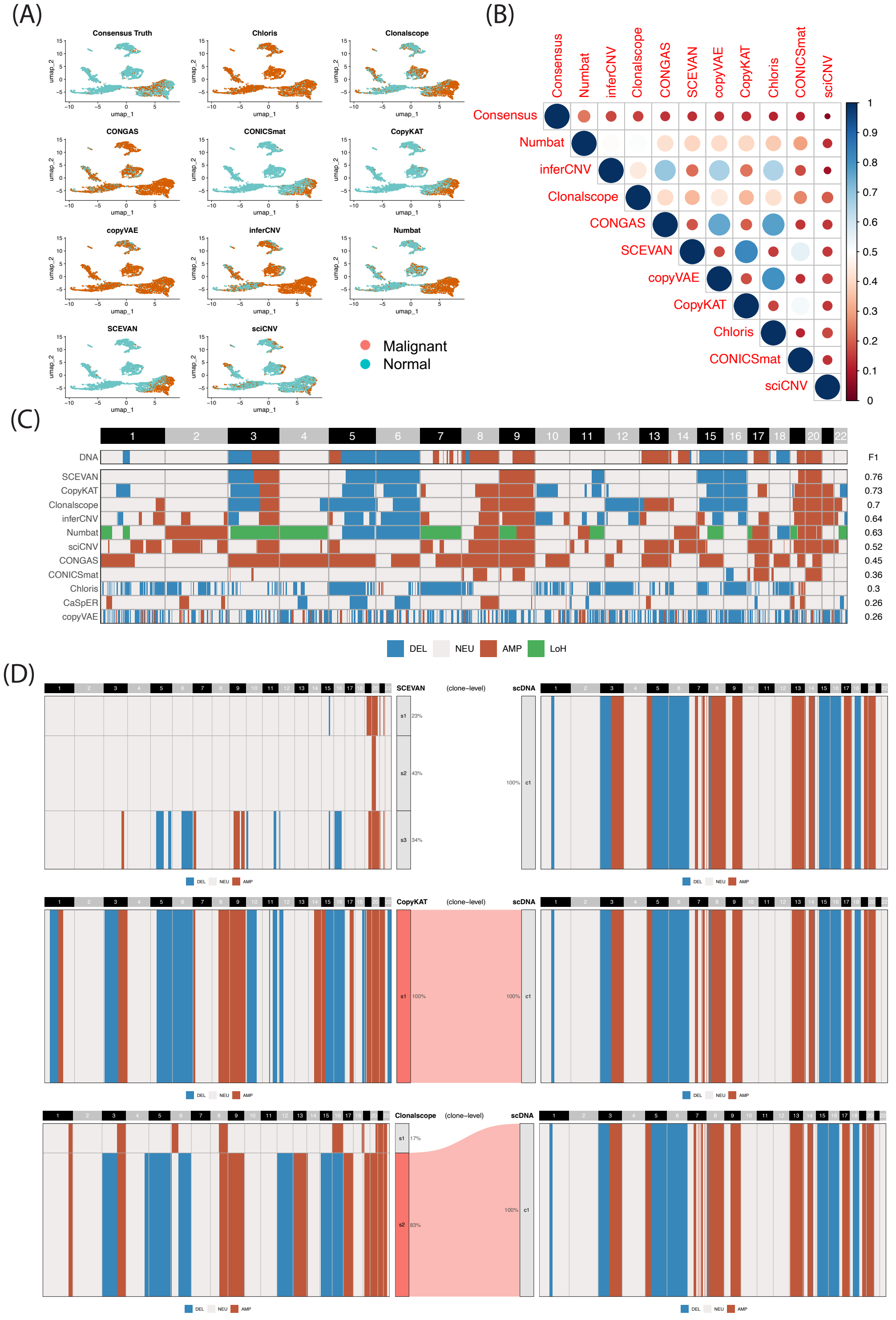


**Supplementary Figure 9: Visualization of P5846 result for B1 and B2 tasks.** (A) UMAP of malignant cell classification. (B) Cross-method concordance score based on the Jaccard index. (C) Bulk-level CNV profiles inferred from scRNA-based CNV methods, with ground truth derived from single-cell DNA sequencing. (D) Subclonal assignments of the top three methods at the bulk level. Only subclones with F1 > 0.6 are assigned.

# P5847

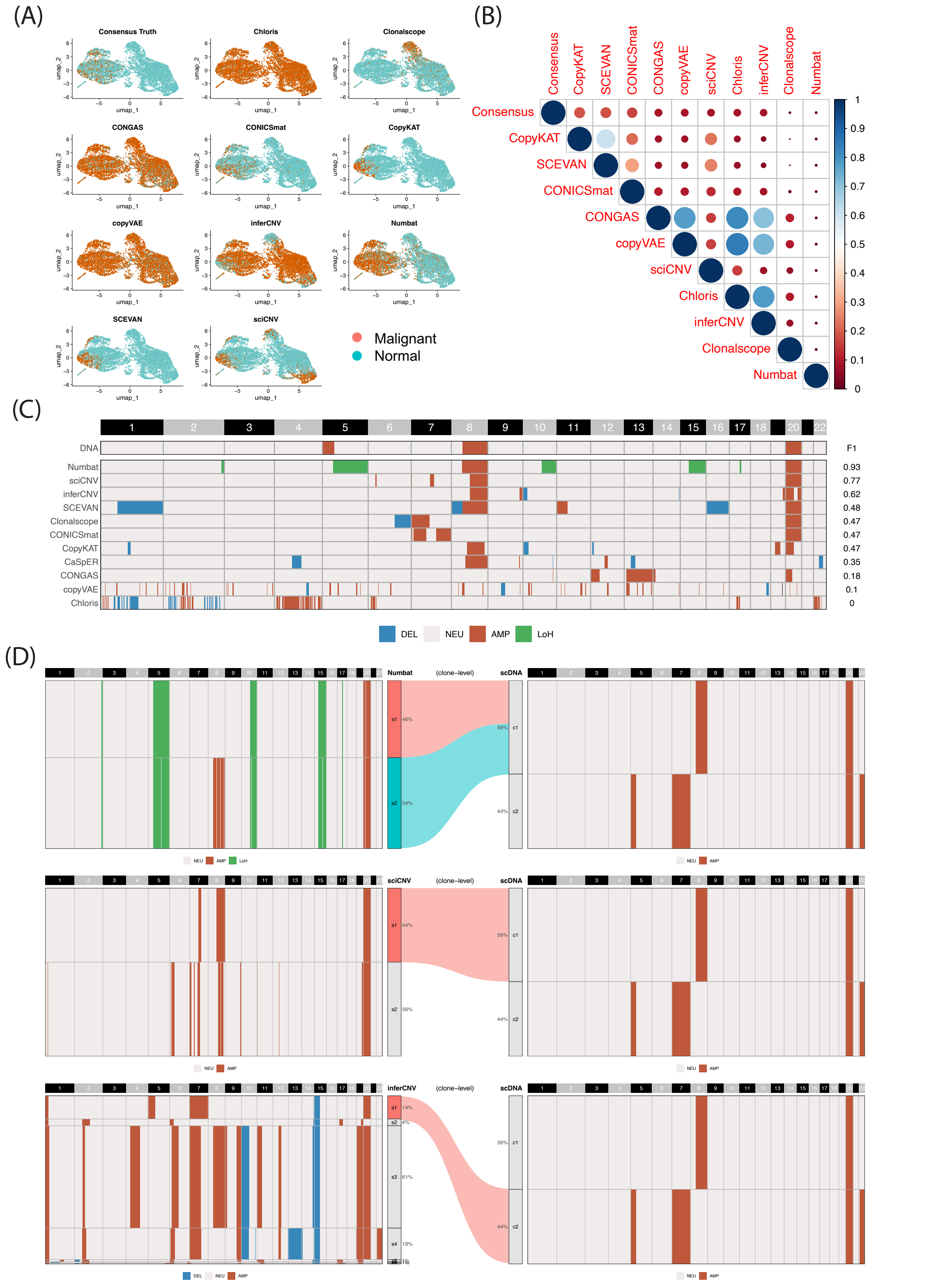


**Supplementary Figure 10: Visualization of P5847 result for B1 and B2 tasks.** (A) UMAP of malignant cell classification. (B) Cross-method concordance score based on the Jaccard index. (C) Bulk-level CNV profiles inferred from scRNA-based CNV methods, with ground truth derived from single-cell DNA sequencing. (D) Subclonal assignments of the top three methods at the bulk level. Only subclones with F1 > 0.6 are assigned.

## P5931

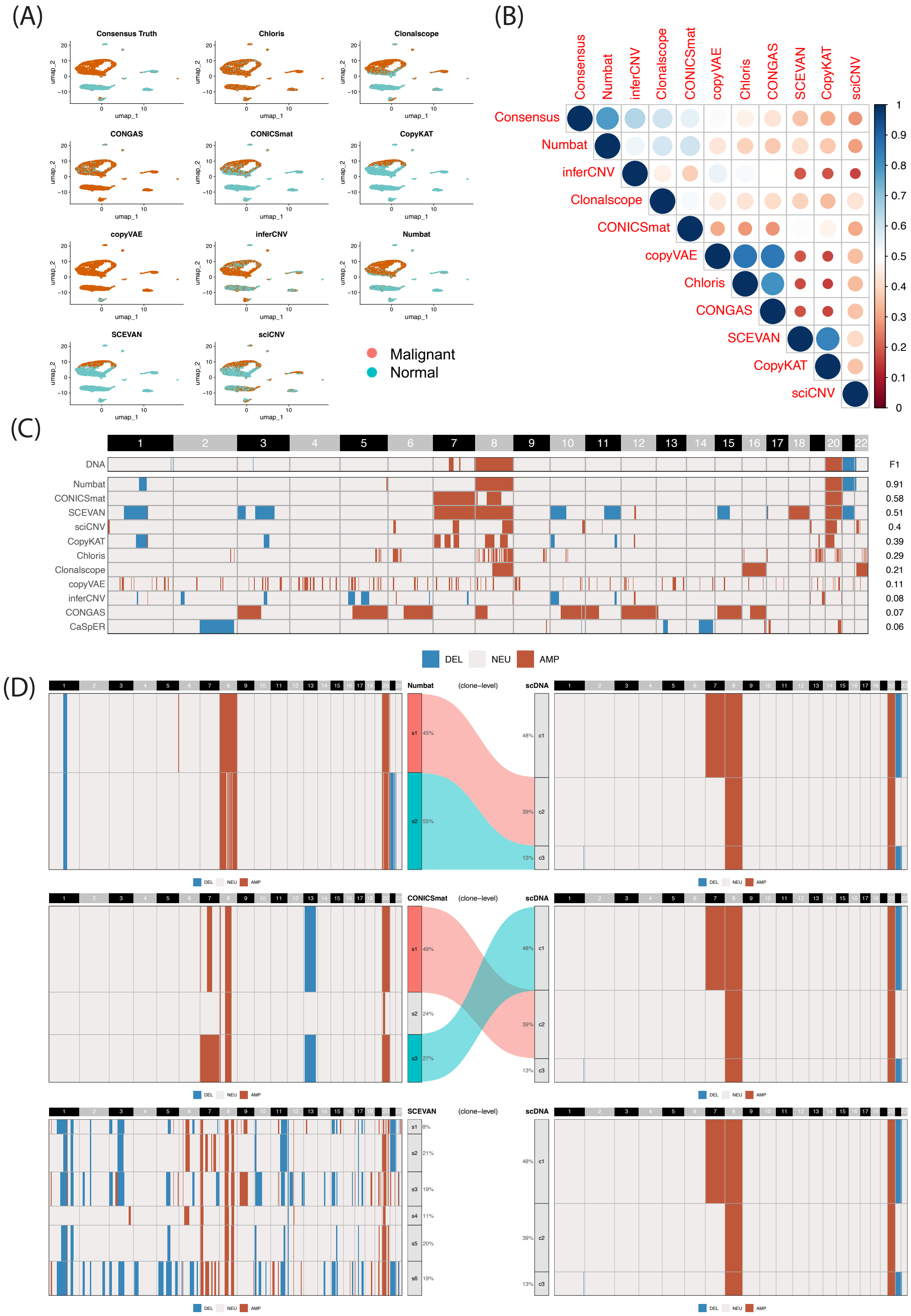


**Supplementary Figure 11: Visualization of P5931 result for B1 and B2 tasks.** (A) UMAP of malignant cell classification. (B) Cross-method concordance score based on the Jaccard index. (C) Bulk-level CNV profiles inferred from scRNA-based CNV methods, with ground truth derived from single-cell DNA sequencing. (D) Subclonal assignments of the top three methods at the bulk level. Only subclones with F1 > 0.6 are assigned.

## P8823

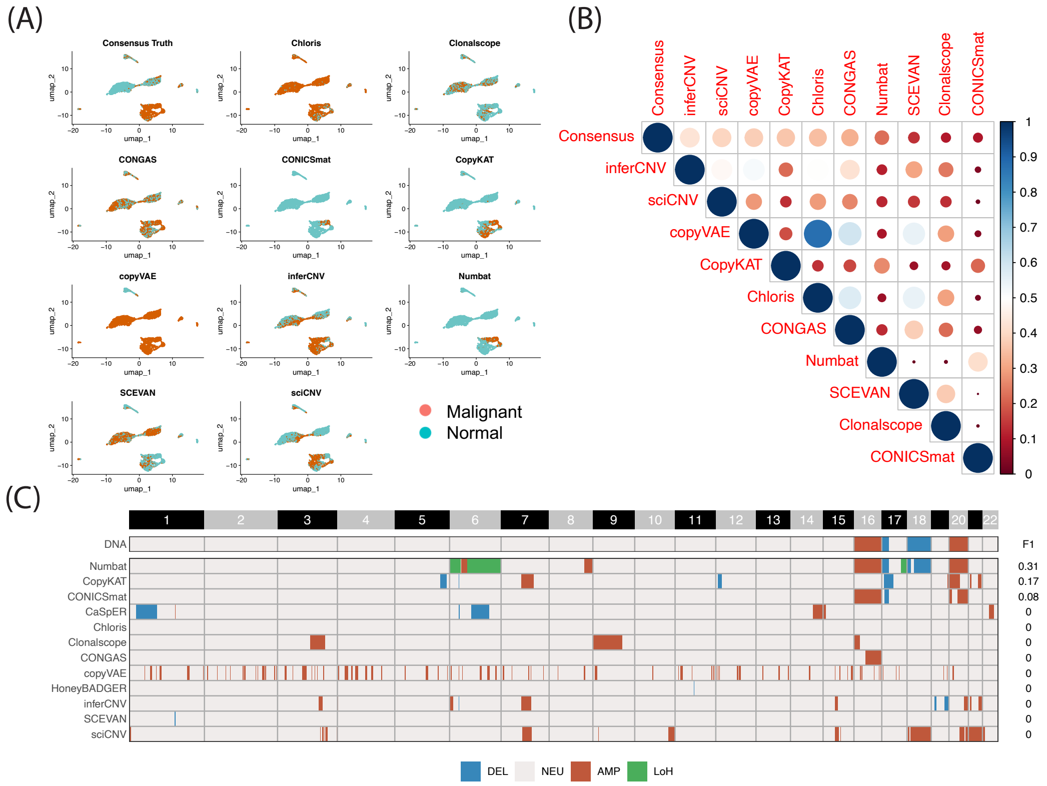


**Supplementary Figure 12: Visualization of P8823 result for B1 and B2 tasks.** (A) UMAP of malignant cell classification. (B) Cross-method concordance score based on the Jaccard index. (C) Bulk-level CNV profiles inferred from scRNA-based CNV methods, with ground truth derived from bulk DNA sequencing.

## MM199

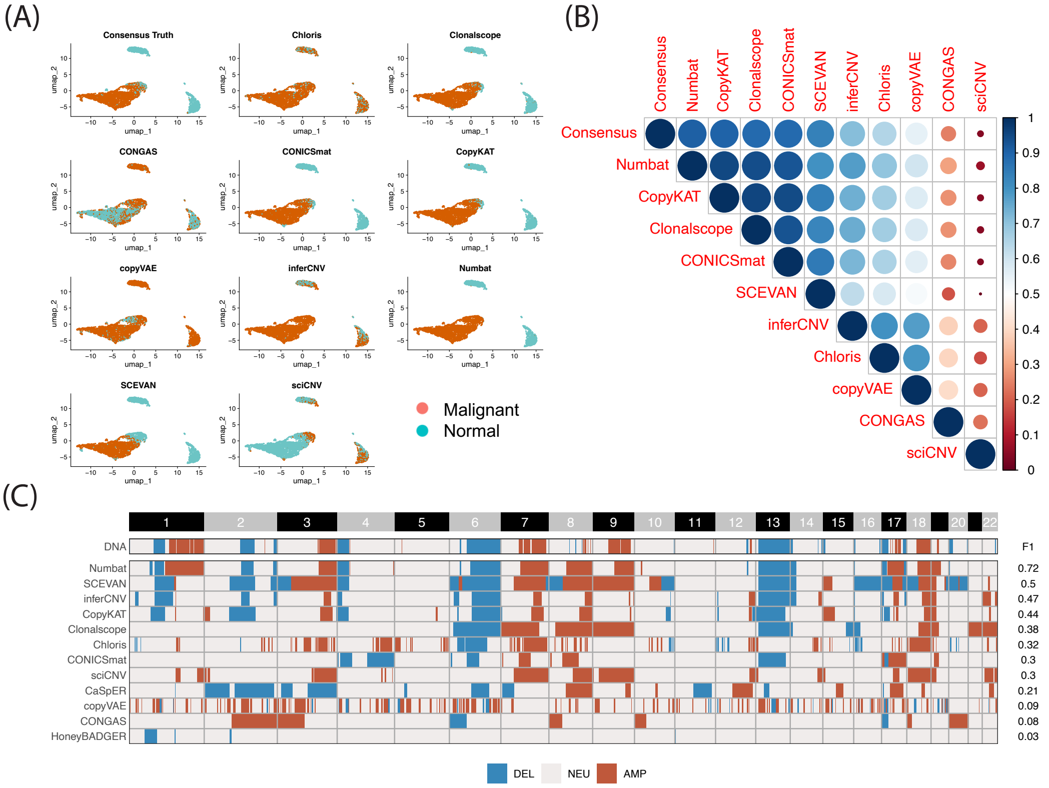


**Supplementary Figure 13: Visualization of MM199 result for B1 and B2 tasks.** (A) UMAP of malignant cell classification. (B) Cross-method concordance score based on the Jaccard index. (C) Bulk-level CNV profiles inferred from scRNA-based CNV methods, with ground truth derived from bulk DNA sequencing.

### Metastases results

## P5915

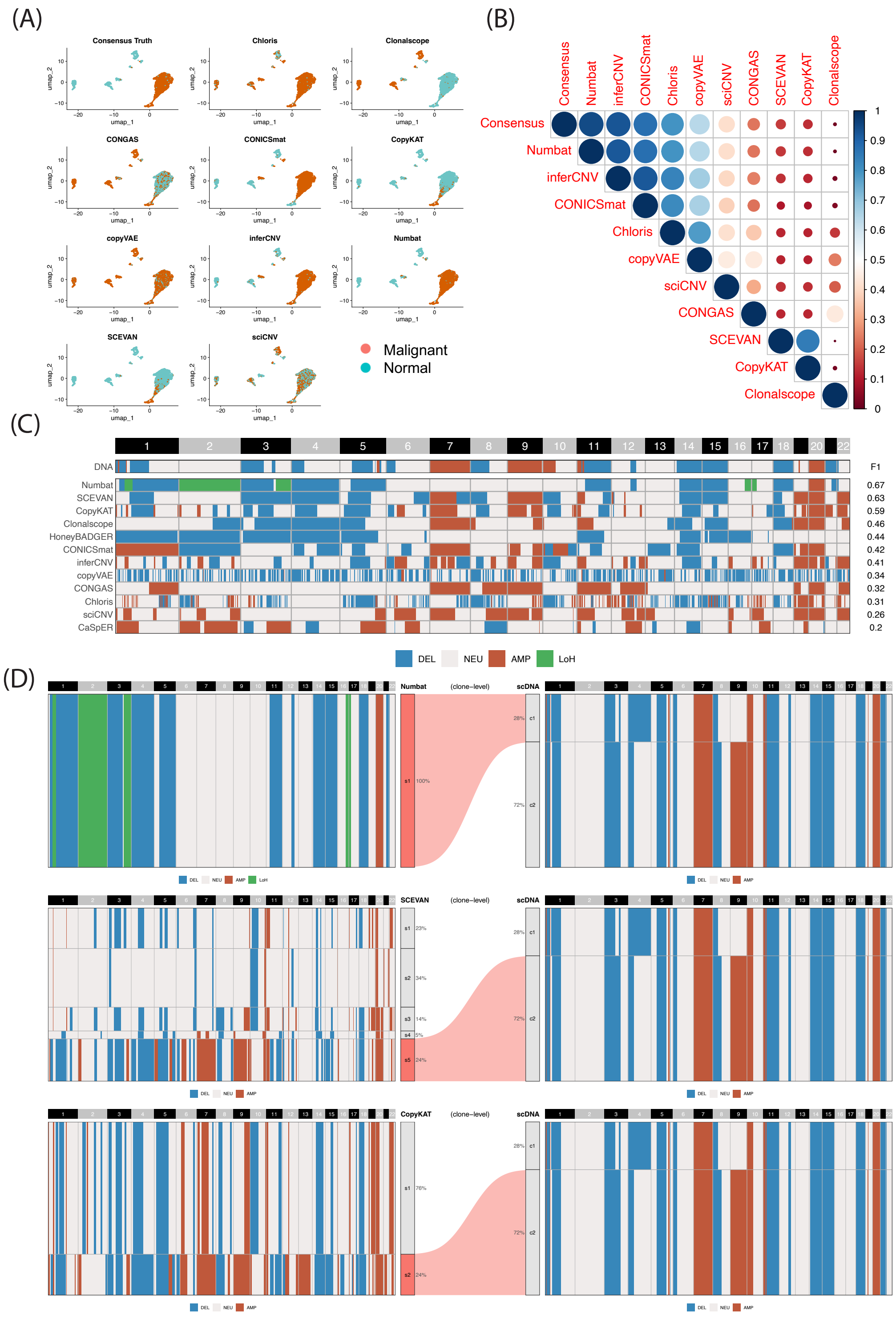


**Supplementary Figure 14: Visualization of P5915 result for B1 and B2 tasks.** (A) UMAP of malignant cell classification. (B) Cross-method concordance score based on the Jaccard index. (C) Bulk-level CNV profiles inferred from scRNA-based CNV methods, with ground truth derived from single-cell DNA sequencing. (D) Subclonal assignments of the top three methods at the bulk level. Only subclones with F1 > 0.6 are assigned.

## P6198

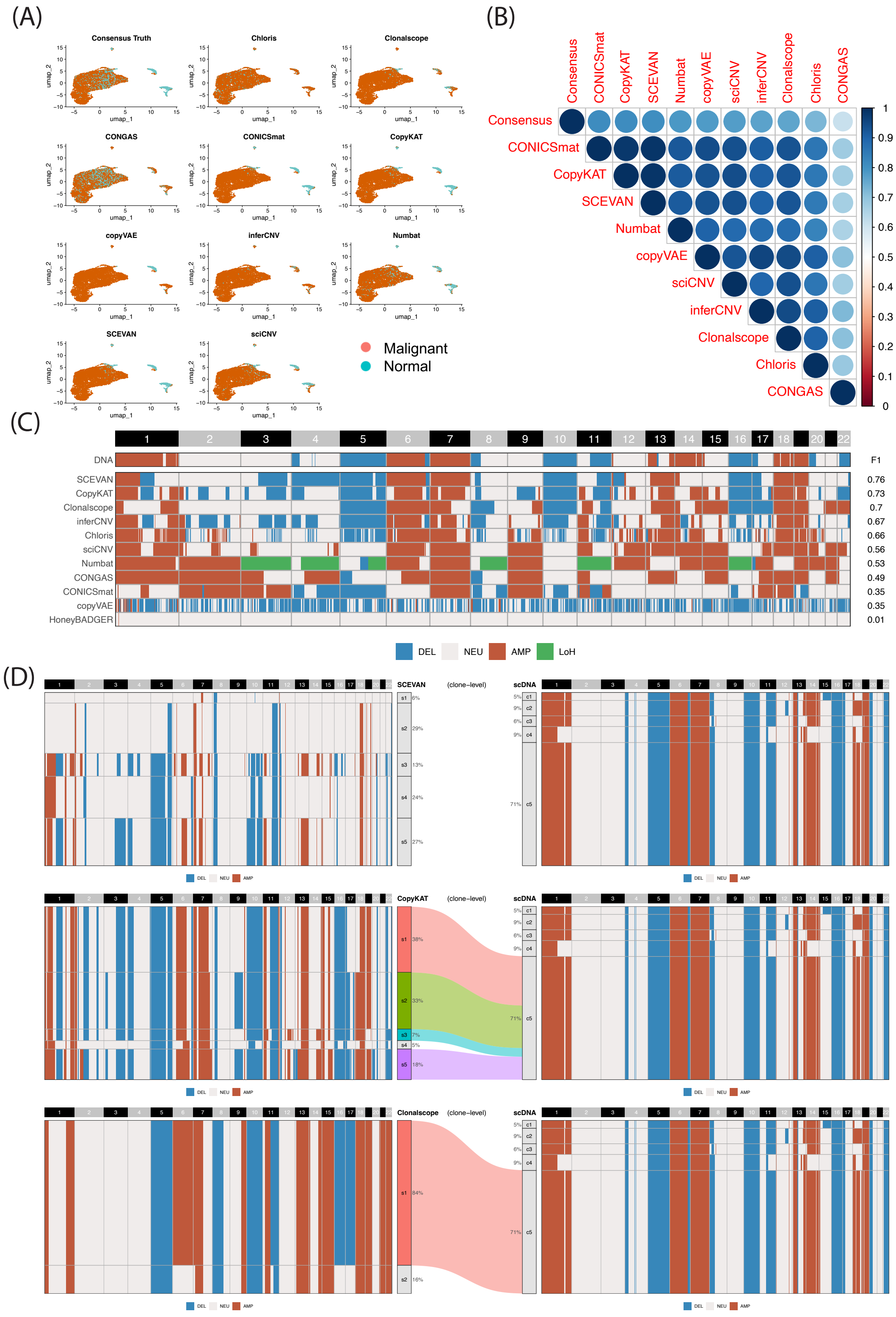


**Supplementary Figure15: Visualization of P6198 result for B1 and B2 tasks.** (A) UMAP of malignant cell classification. (B) Cross-method concordance score based on the Jaccard index. (C) Bulk-level CNV profiles inferred from scRNA-based CNV methods, with ground truth derived from single-cell DNA sequencing. (D) Subclonal assignments of the top three methods at the bulk level. Only subclones with F1 > 0.6 are assigned.

## P6335

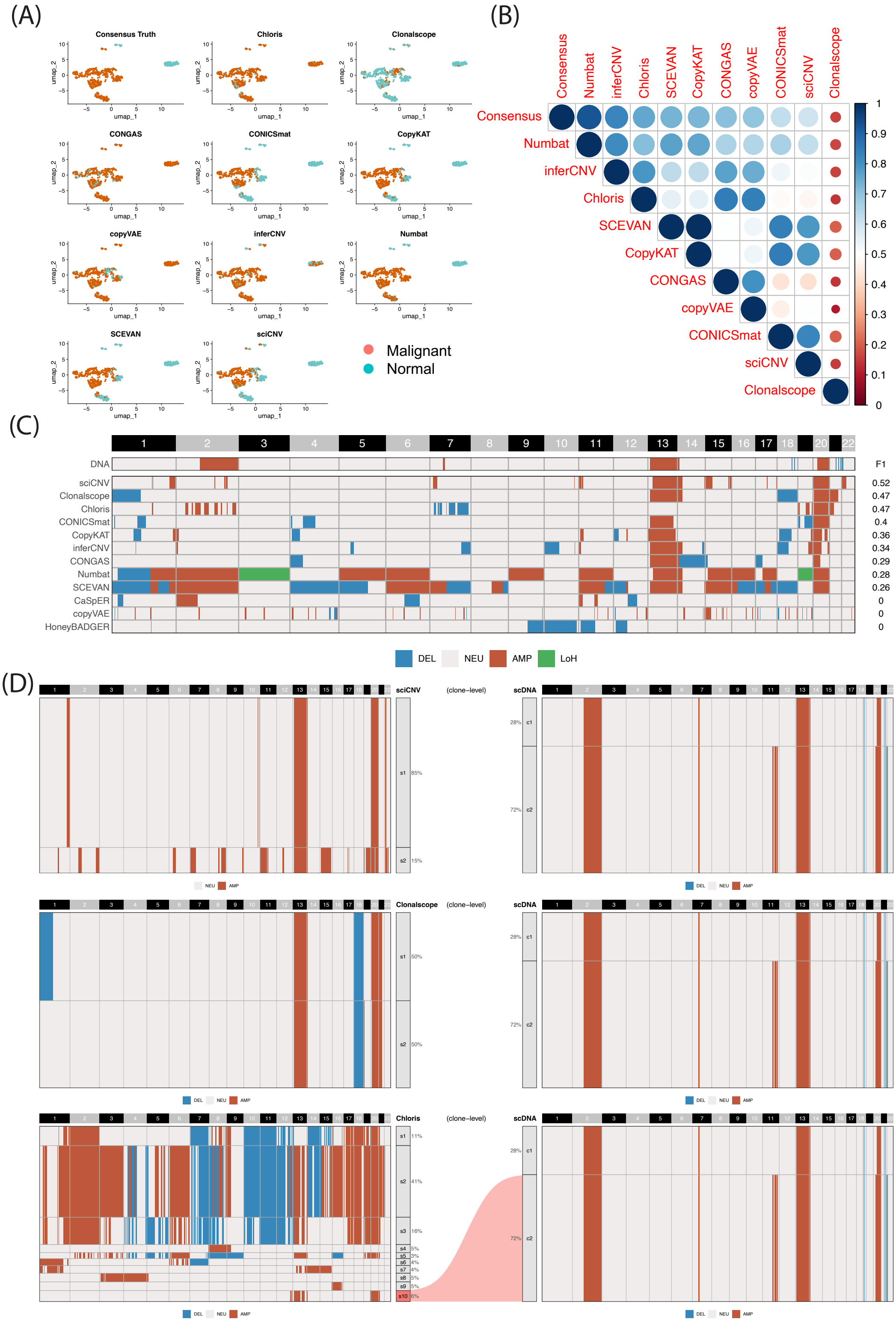


**Supplementary Figure 16: Visualization of P6335 result for B1 and B2 tasks.** (A) UMAP of malignant cell classification. (B) Cross-method concordance score based on the Jaccard index. (C) Bulk-level CNV profiles inferred from scRNA-based CNV methods, with ground truth derived from single-cell DNA sequencing. (D) Subclonal assignments of the top three methods at the bulk level. Only subclones with F1 > 0.6 are assigned.

# P6593

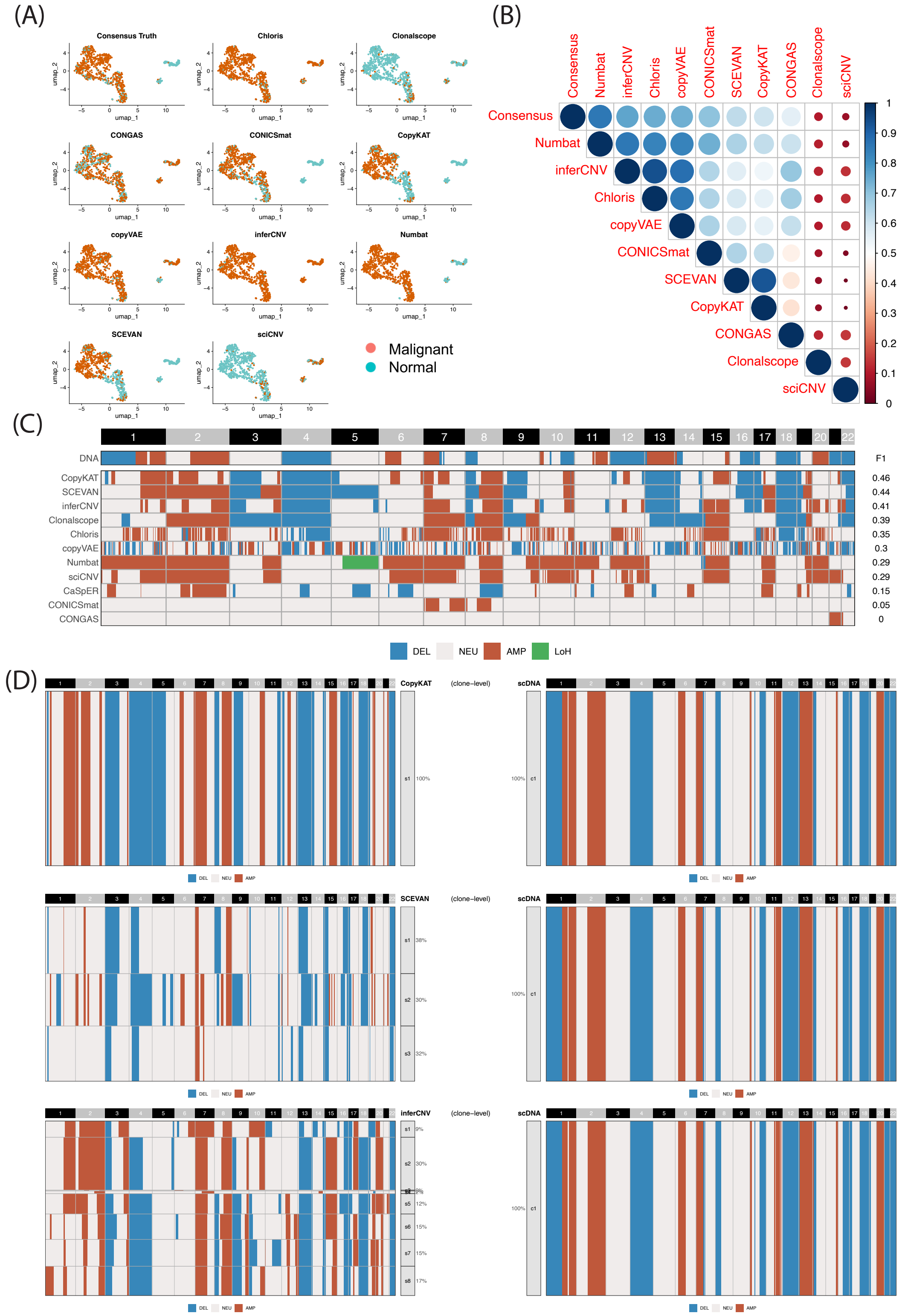


**Supplementary Figure 17: Visualization of P6593 result for B1 and B2 tasks.** (A) UMAP of malignant cell classification. (B) Cross-method concordance score based on the Jaccard index. (C) Bulk-level CNV profiles inferred from scRNA-based CNV methods, with ground truth derived from single-cell DNA sequencing. (D) Subclonal assignments of the top three methods at the bulk level. Only subclones with F1 > 0.6 are assigned.

#### BoML

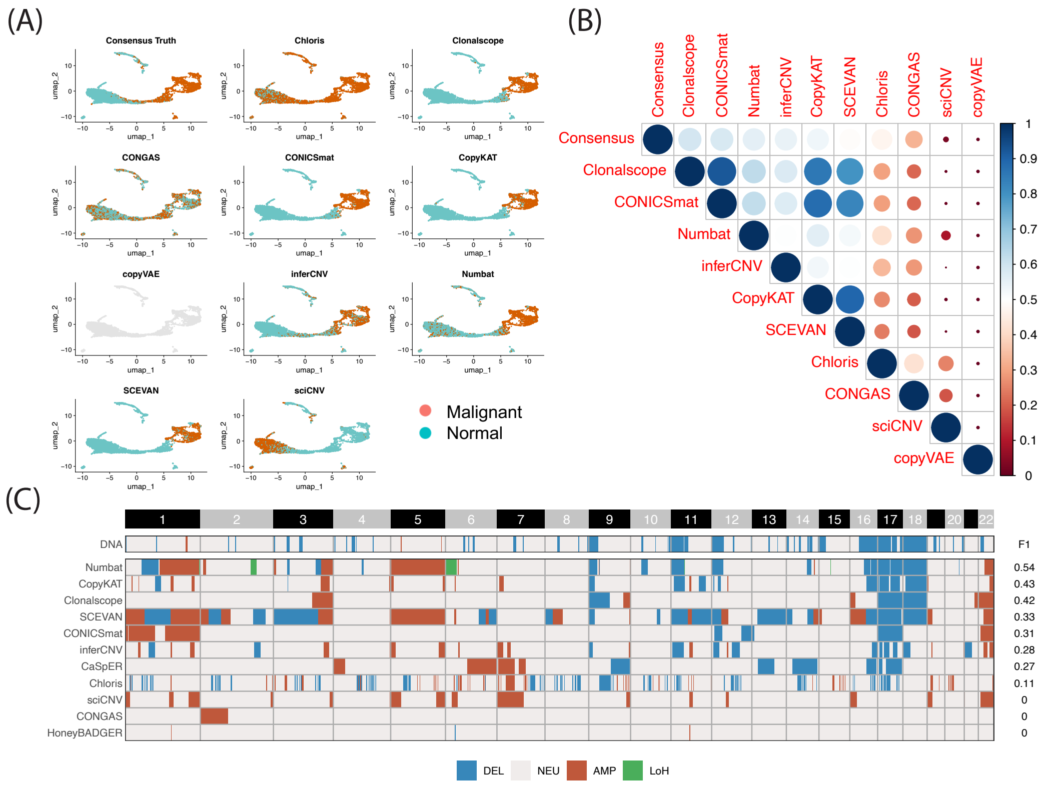


**Supplementary Figure 18: Visualization of BoML result for B1 and B2 tasks.** (A) UMAP of malignant cell classification. (B) Cross-method concordance score based on the Jaccard index. (C) Bulk-level CNV profiles inferred from scRNA-based CNV methods, with ground truth derived from bulk DNA sequencing.

#### BoMR

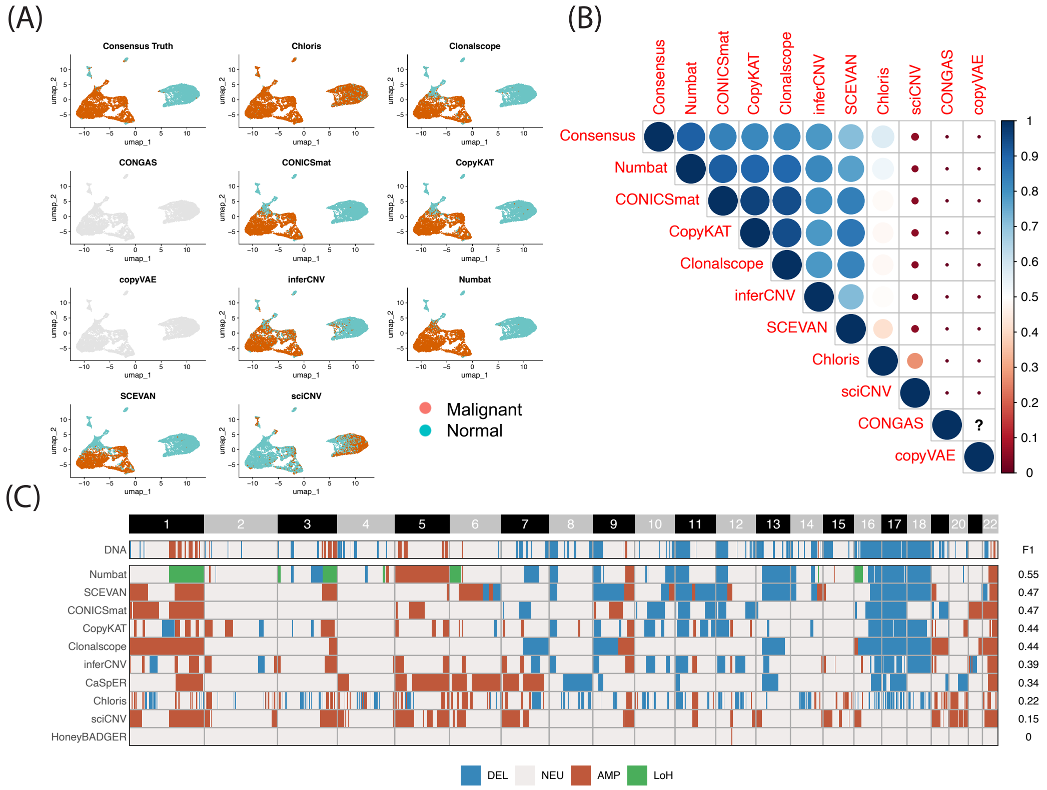


**Supplementary Figure 19: Visualization of BoMR result for B1 and B2 tasks.** (A) UMAP of malignant cell classification. (B) Cross-method concordance score based on the Jaccard index. (C) Bulk-level CNV profiles inferred from scRNA-based CNV methods, with ground truth derived from bulk DNA sequencing.

### Cell lines results

#### HGC-27

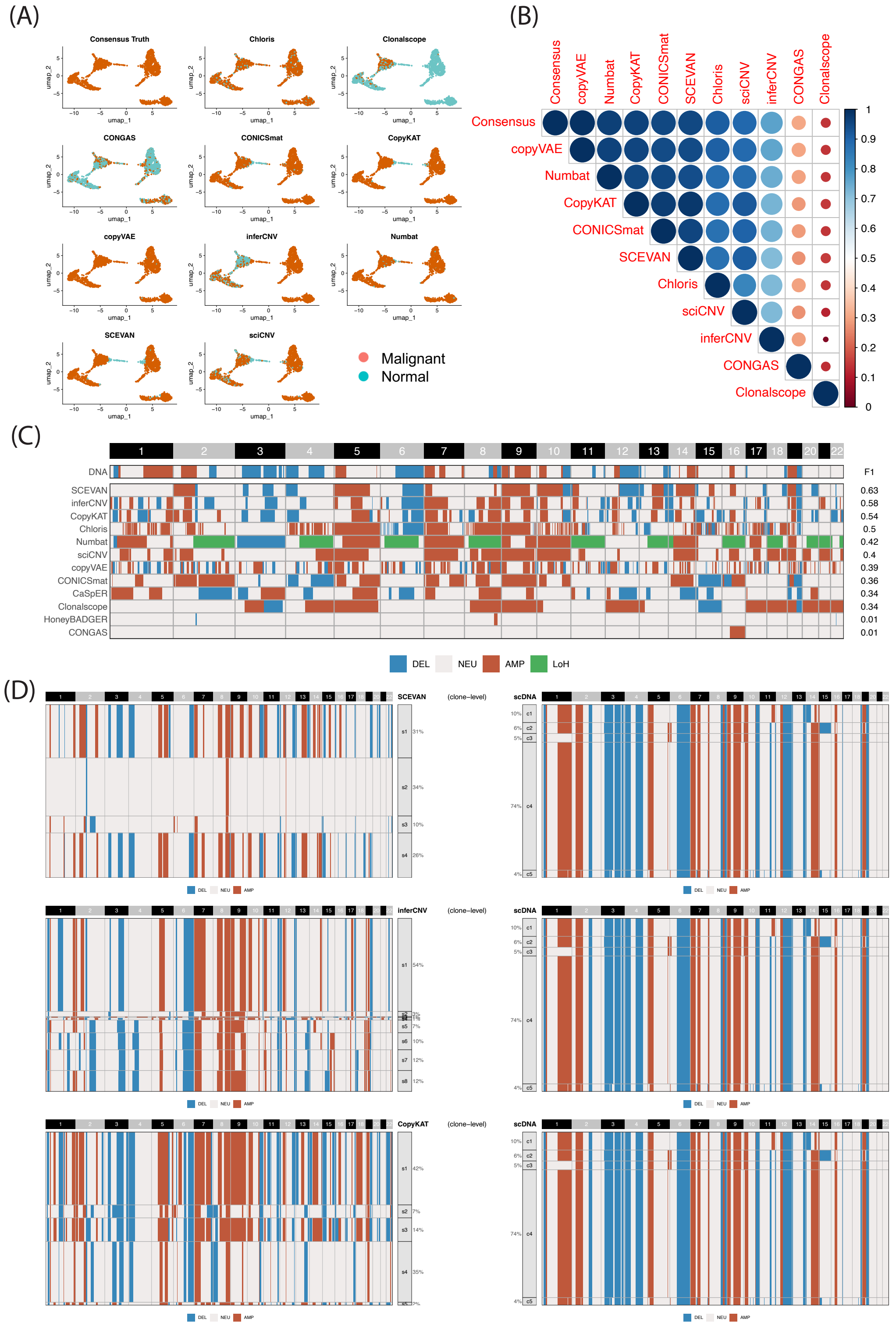

**Supplementary Figure 20: Visualization of HGC-27 result for B1 and B2 tasks.** (A) UMAP of malignant cell classification. (B) Cross-method concordance score based on the Jaccard index. (C) Bulk-level CNV profiles inferred from scRNA-based CNV methods, with ground truth derived from single-cell DNA sequencing. (D) Subclonal assignments of the top three methods at the bulk level. Only subclones with F1 > 0.6 are assigned.

#### KATOIII

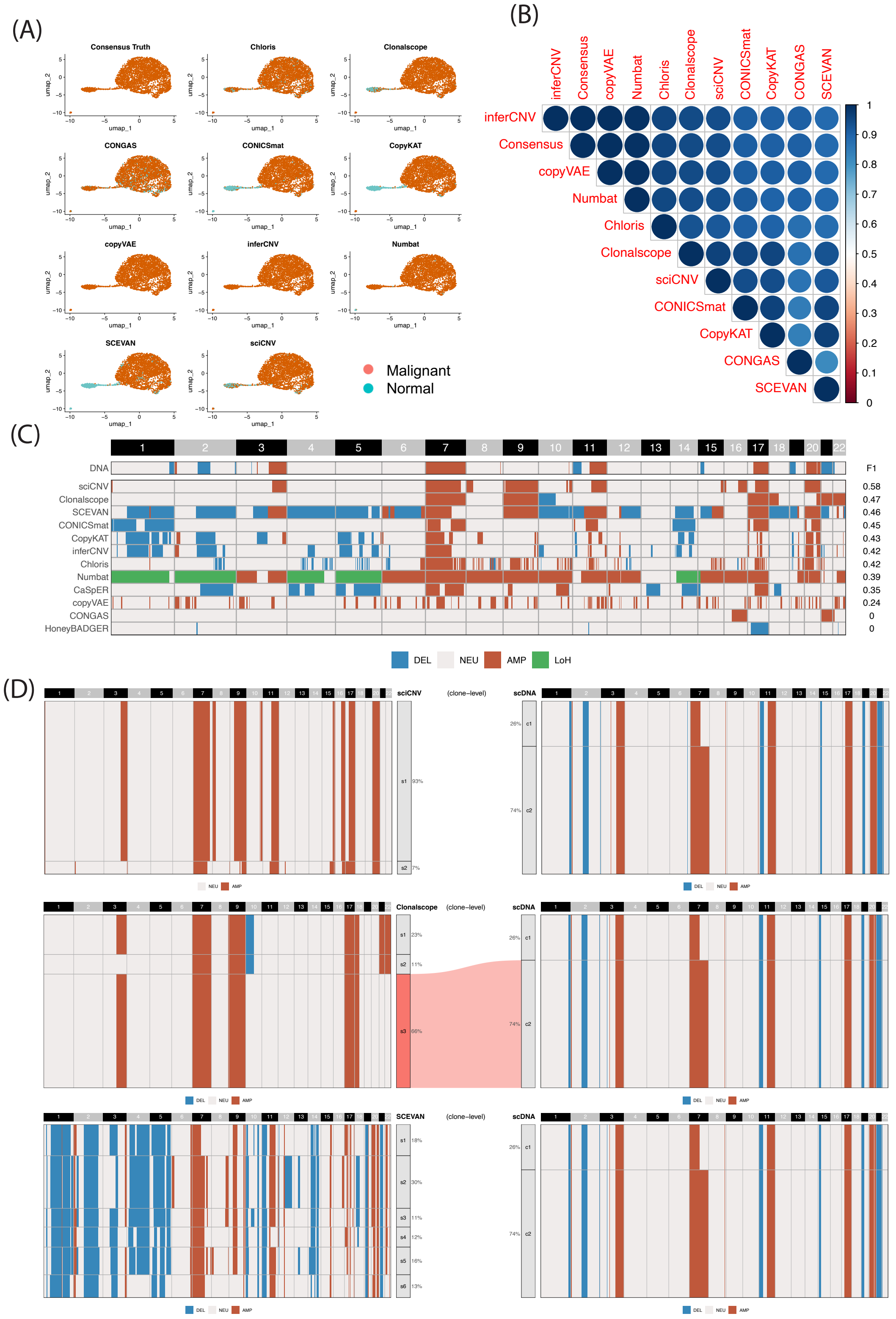


**Supplementary Figure 21: Visualization of KATOIII result for B1 and B2 tasks.** (A) UMAP of malignant cell classification. (B) Cross-method concordance score based on the Jaccard index. (C) Bulk-level CNV profiles inferred from scRNA-based CNV methods, with ground truth derived from single-cell DNA sequencing. (D) Subclonal assignments of the top three methods at the bulk level. Only subclones with F1 > 0.6 are assigned.

### MKN-45

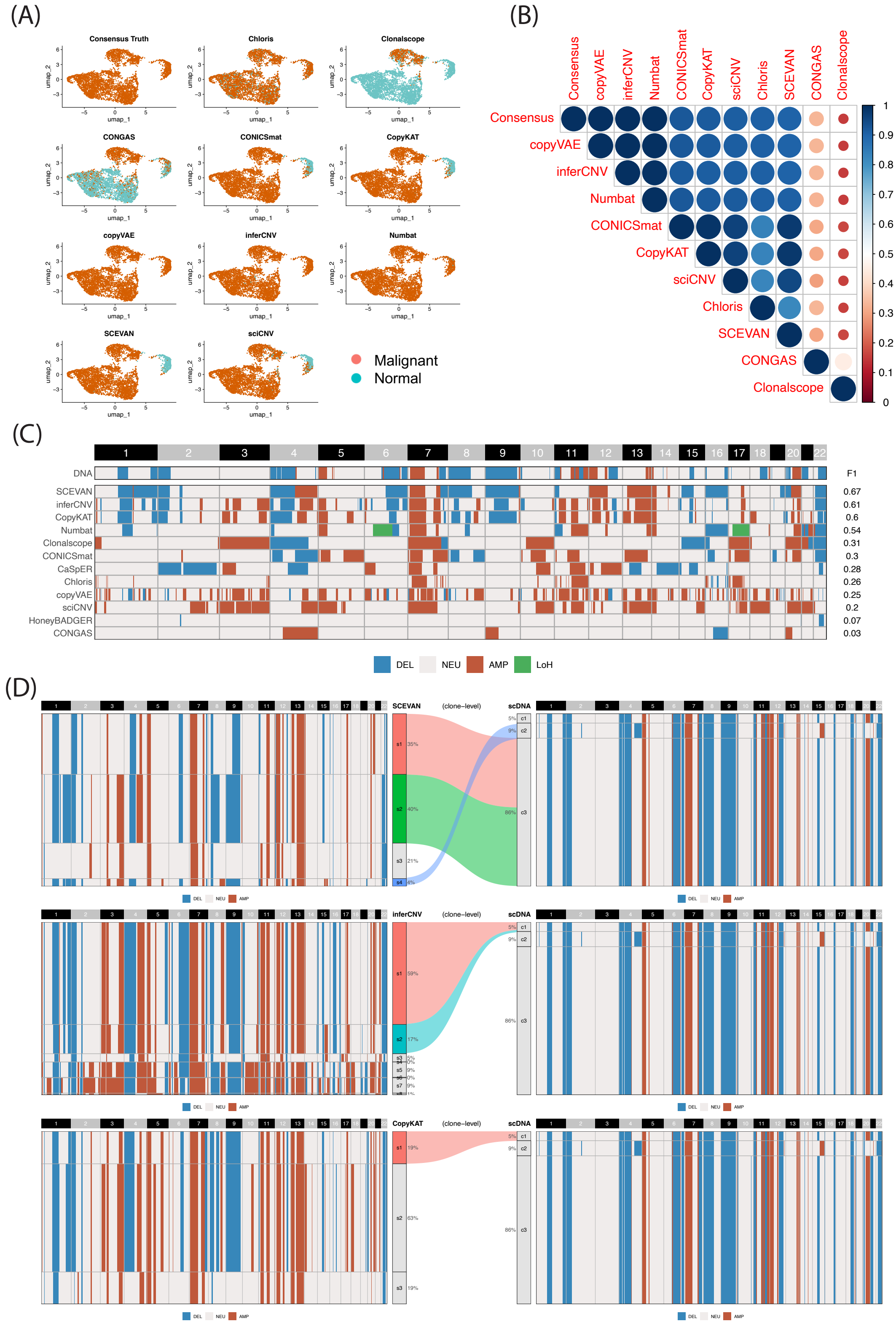


**Supplementary Figure 22: Visualization of MKN-45 result for B1 and B2 tasks.** (A) UMAP of malignant cell classification. (B) Cross-method concordance score based on the Jaccard index. (C) Bulk-level CNV profiles inferred from scRNA-based CNV methods, with ground truth derived from single-cell DNA sequencing. (D) Subclonal assignments of the top three methods at the bulk level. Only subclones with F1 > 0.6 are assigned.

#### NCI-N87

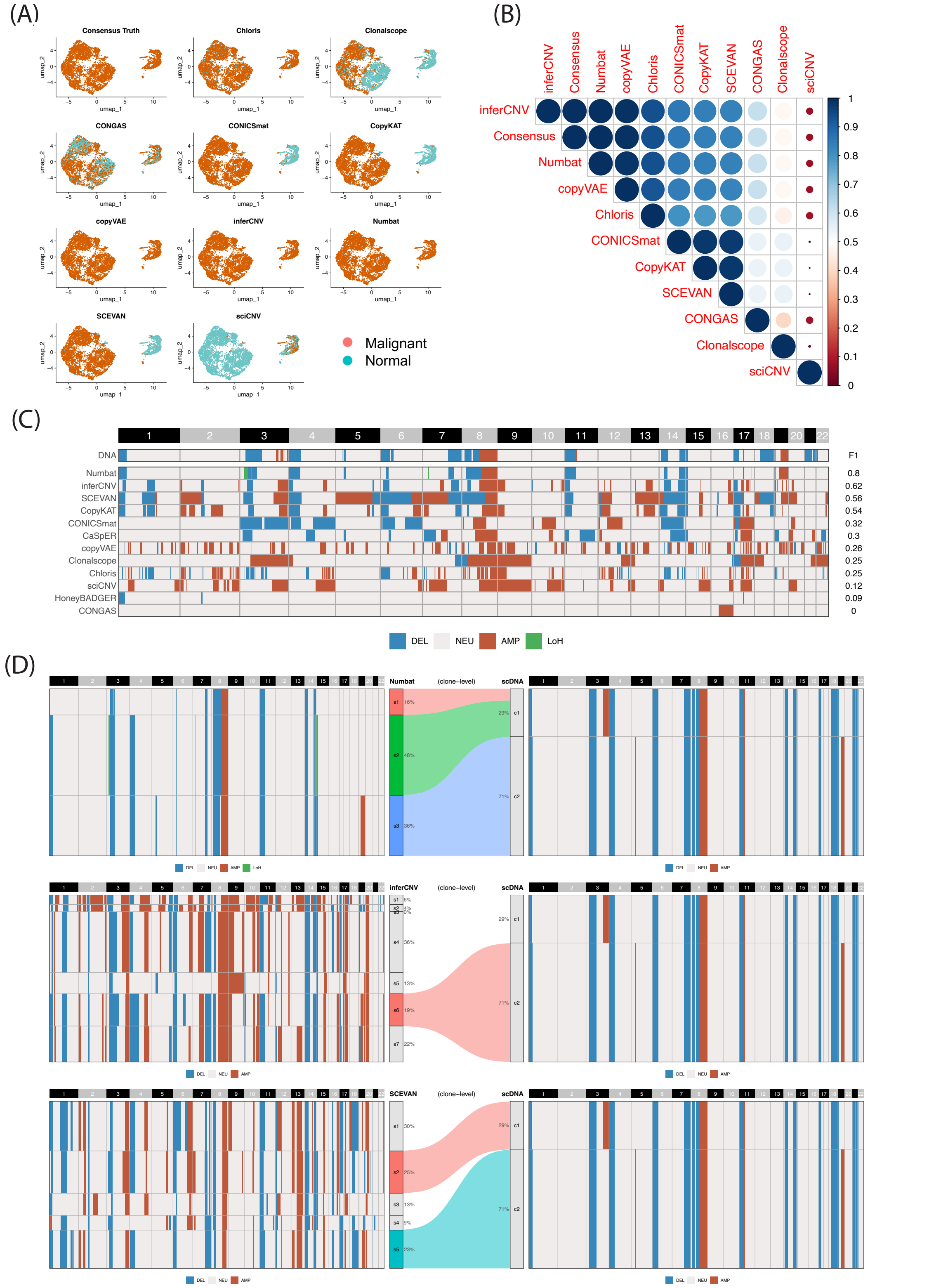


**Supplementary Figure 23: Visualization of NCI-N87 result for B1 and B2 tasks.** (A) UMAP of malignant cell classification. (B) Cross-method concordance score based on the Jaccard index. (C) Bulk-level CNV profiles inferred from scRNA-based CNV methods, with ground truth derived from single-cell DNA sequencing. (D) Subclonal assignments of the top three methods at the bulk level. Only subclones with F1 > 0.6 are assigned.

### NUGC-4

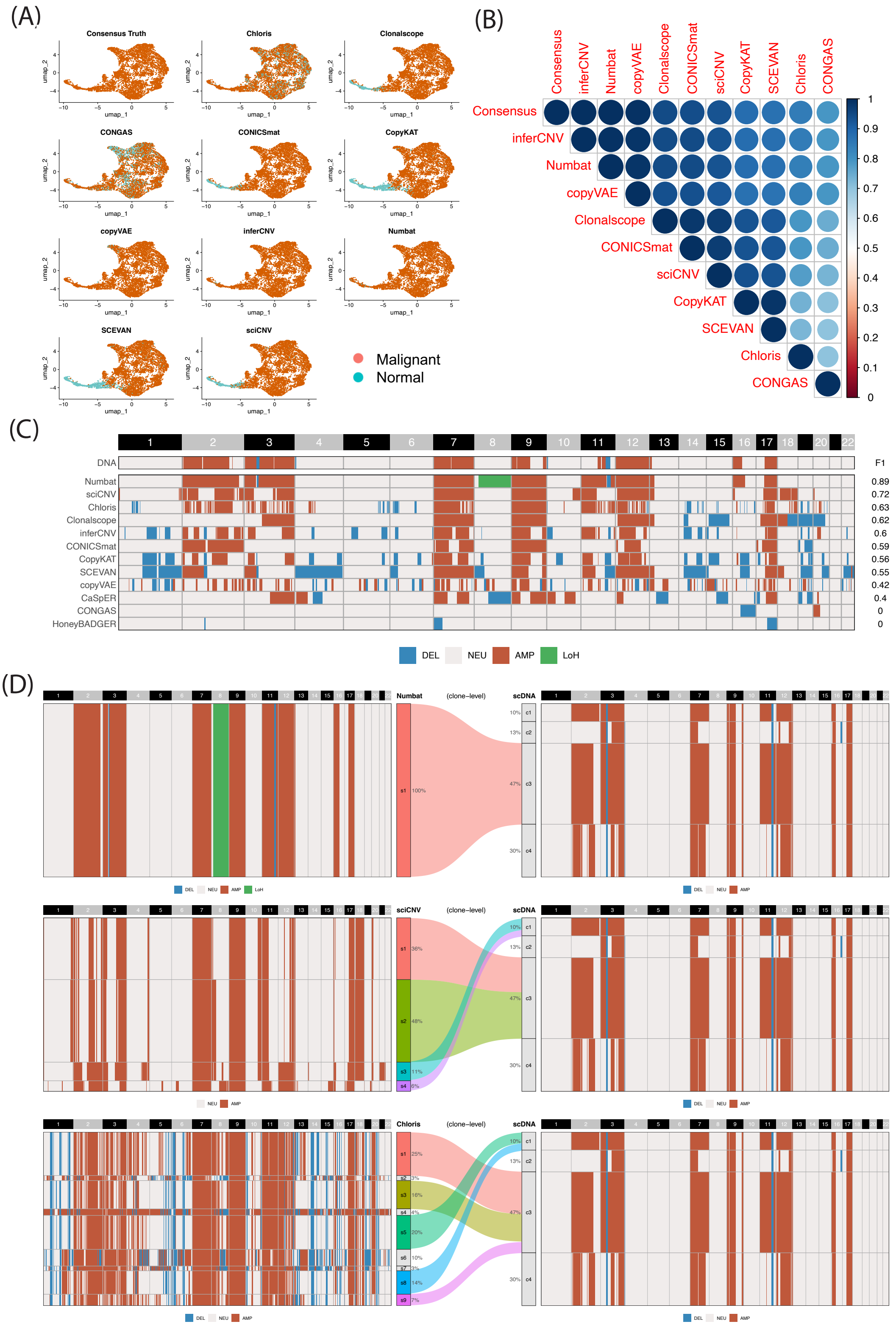


**Supplementary Figure 24: Visualization of NUGC-4 result for B1 and B2 tasks.** (A) UMAP of malignant cell classification. (B) Cross-method concordance score based on the Jaccard index. (C) Bulk-level CNV profiles inferred from scRNA-based CNV methods, with ground truth derived from single-cell DNA sequencing. (D) Subclonal assignments of the top three methods at the bulk level. Only subclones with F1 > 0.6 are assigned.

#### SNU-16

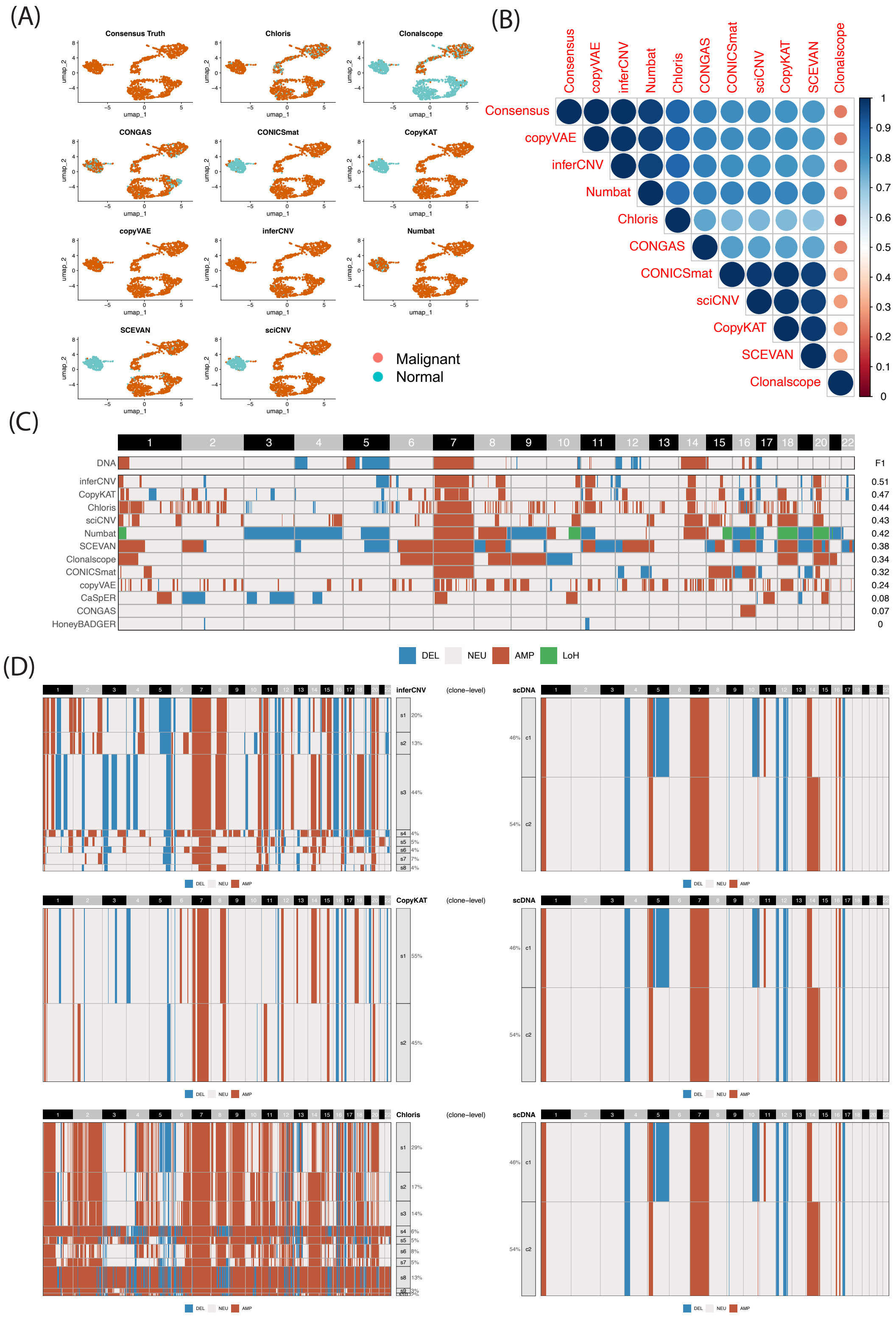


**Supplementary Figure 25: Visualization of SNU-16 result for B1 and B2 tasks.** (A) UMAP of malignant cell classification. (B) Cross-method concordance score based on the Jaccard index. (C) Bulk-level CNV profiles inferred from scRNA-based CNV methods, with ground truth derived from single-cell DNA sequencing. (D) Subclonal assignments of the top three methods at the bulk level. Only subclones with F1 > 0.6 are assigned.

#### SNU-601

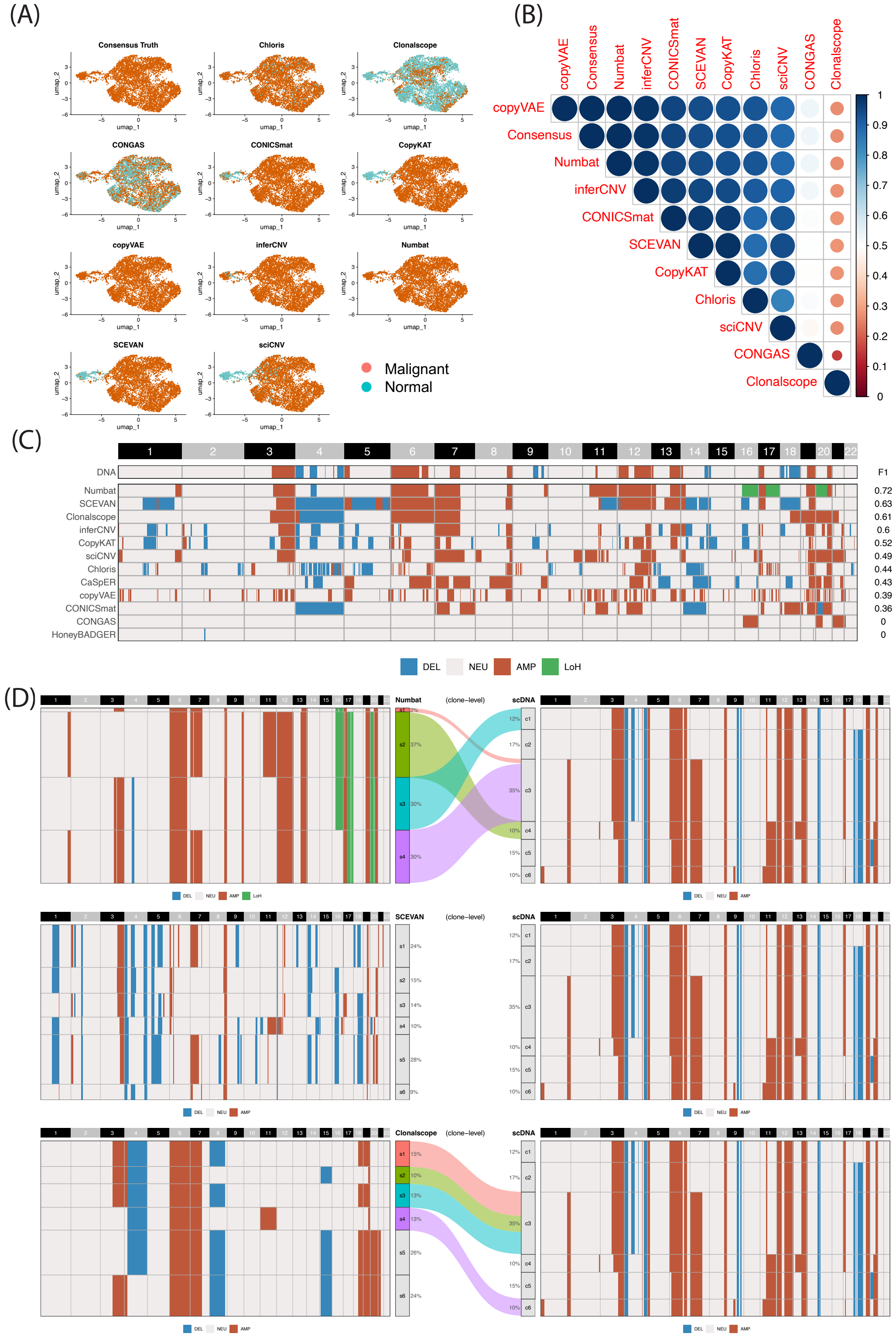


**Supplementary Figure 26: Visualization of SNU-601 result for B1 and B2 tasks.** (A) UMAP of malignant cell classification. (B) Cross-method concordance score based on the Jaccard index. (C) Bulk-level CNV profiles inferred from scRNA-based CNV methods, with ground truth derived from single-cell DNA sequencing. (D) Subclonal assignments of the top three methods at the bulk level. Only subclones with F1 > 0.6 are assigned.

### SNU-638

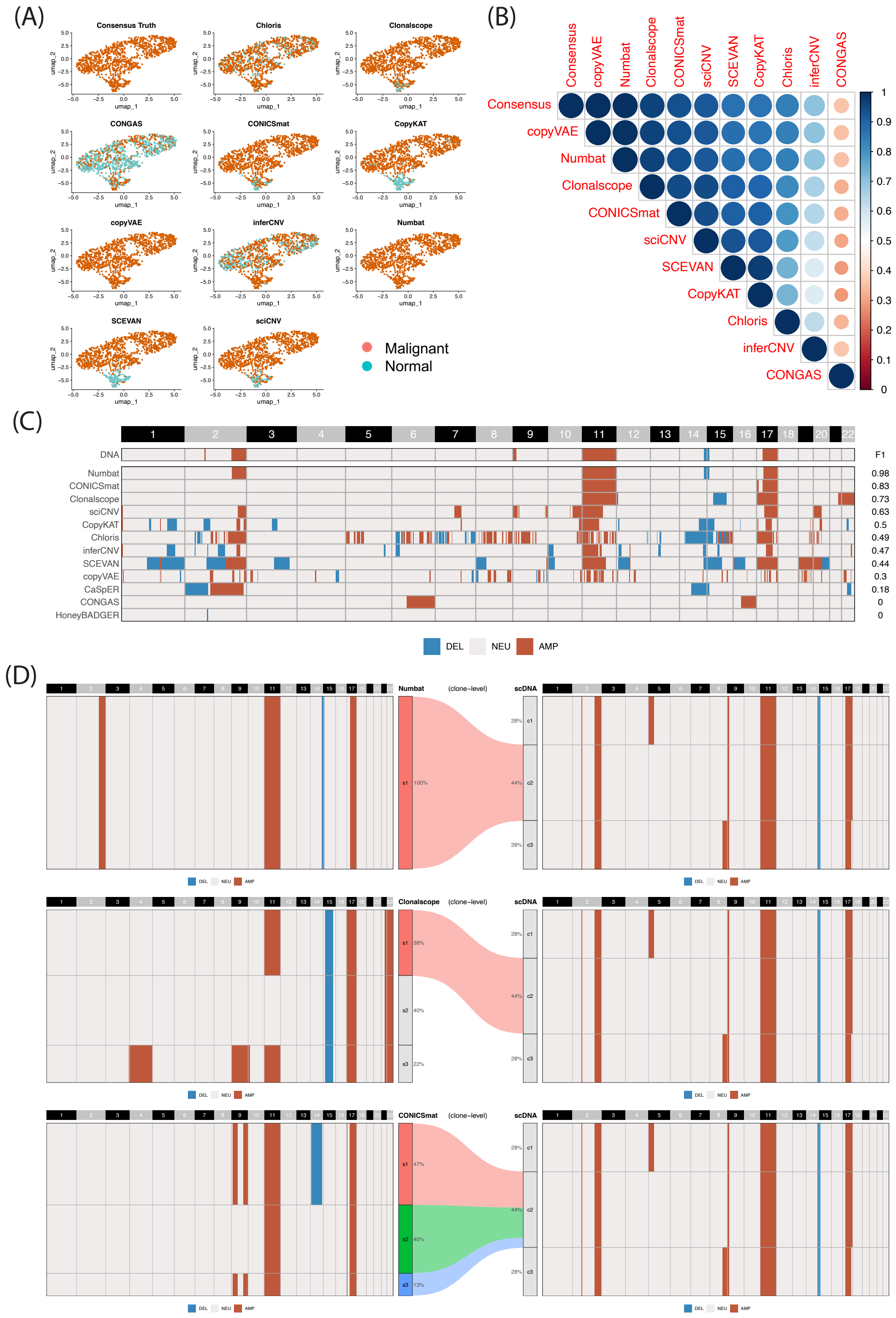


**Supplementary Figure 27: Visualization of SNU-638 result for B1 and B2 tasks.** (A) UMAP of malignant cell classification. (B) Cross-method concordance score based on the Jaccard index. (C) Bulk-level CNV profiles inferred from scRNA-based CNV methods, with ground truth derived from single-cell DNA sequencing. (D) Subclonal assignments of the top three methods at the bulk level. Only subclones with F1 > 0.6 are assigned.

### SNU-668

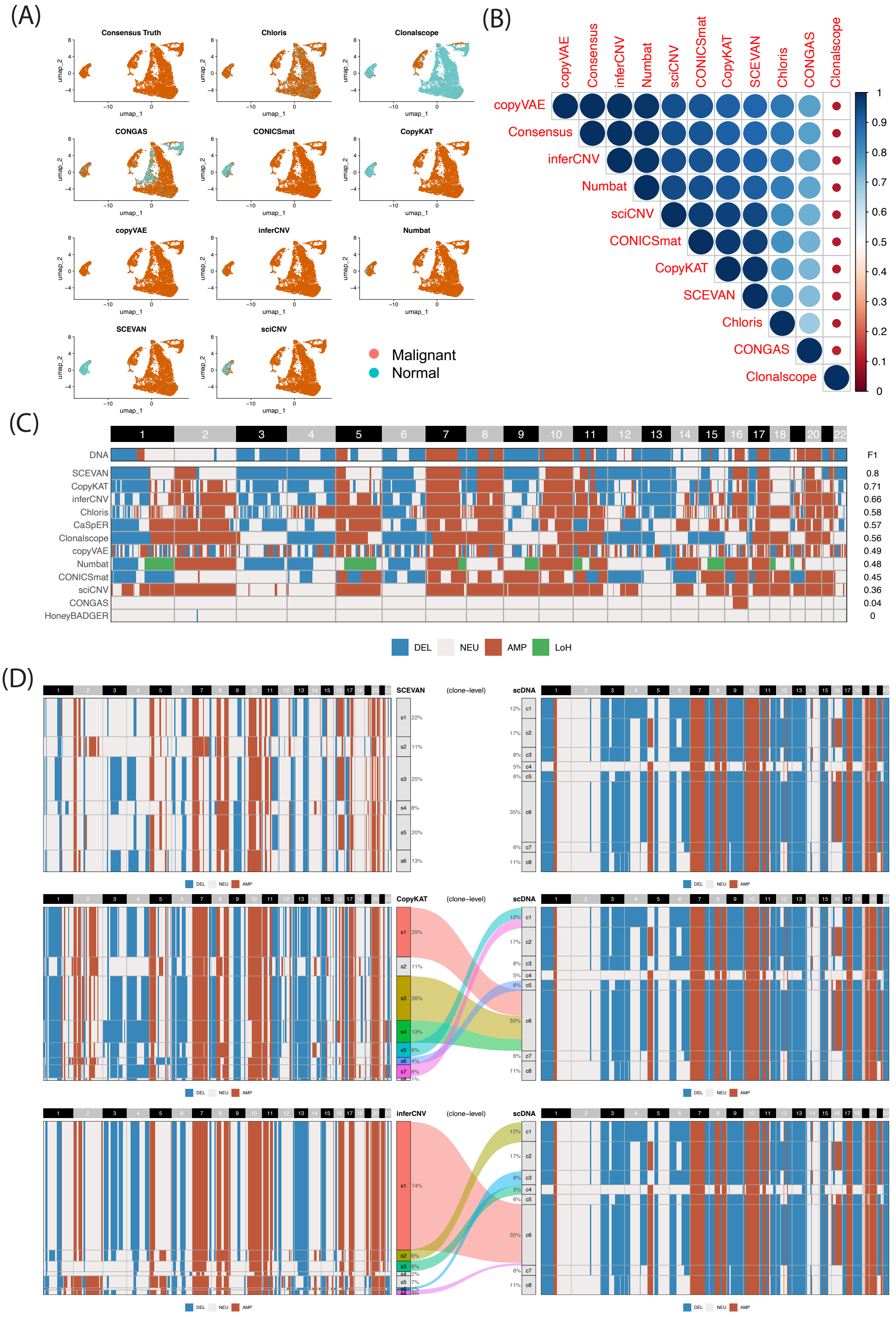


**Supplementary Figure 28: Visualization of SNU-668 result for B1 and B2 tasks.** (A) UMAP of malignant cell classification. (B) Cross-method concordance score based on the Jaccard index. (C) Bulk-level CNV profiles inferred from scRNA-based CNV methods, with ground truth derived from single-cell DNA sequencing. (D) Subclonal assignments of the top three methods at the bulk level. Only subclones with F1 > 0.6 are assigned.

### PDOs results

#### IPM-BO-46-early

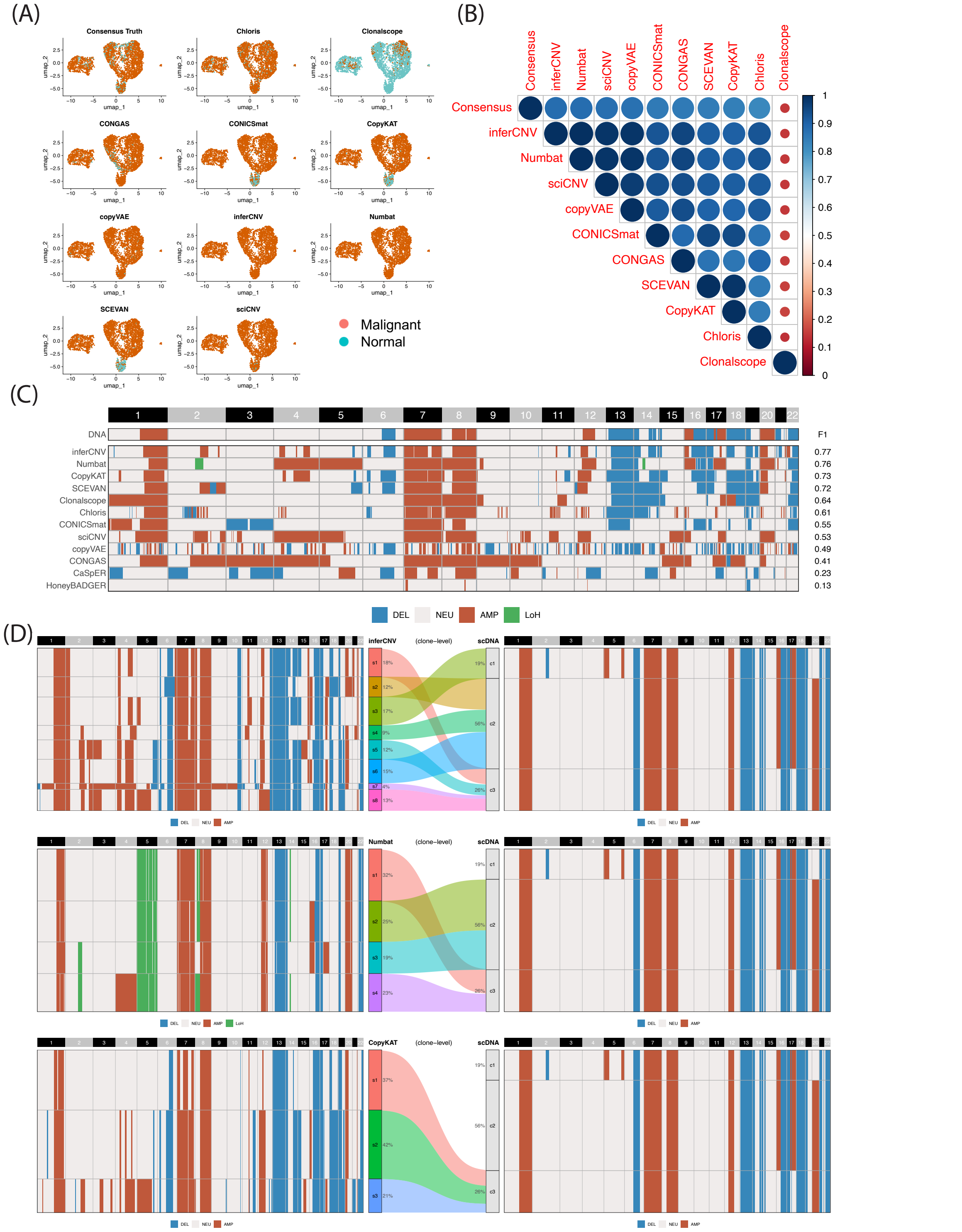


**Supplementary Figure 29: Visualization of IPM-BO-46-early result for B1 and B2 tasks.** (A) UMAP of malignant cell classification. (B) Cross-method concordance score based on the Jaccard index. (C) Bulk-level CNV profiles inferred from scRNA-based CNV methods, with ground truth derived from single-cell DNA sequencing. (D) Subclonal assignments of the top three methods at the bulk level. Only subclones with F1 > 0.6 are assigned.

#### IPM-BO-46-late

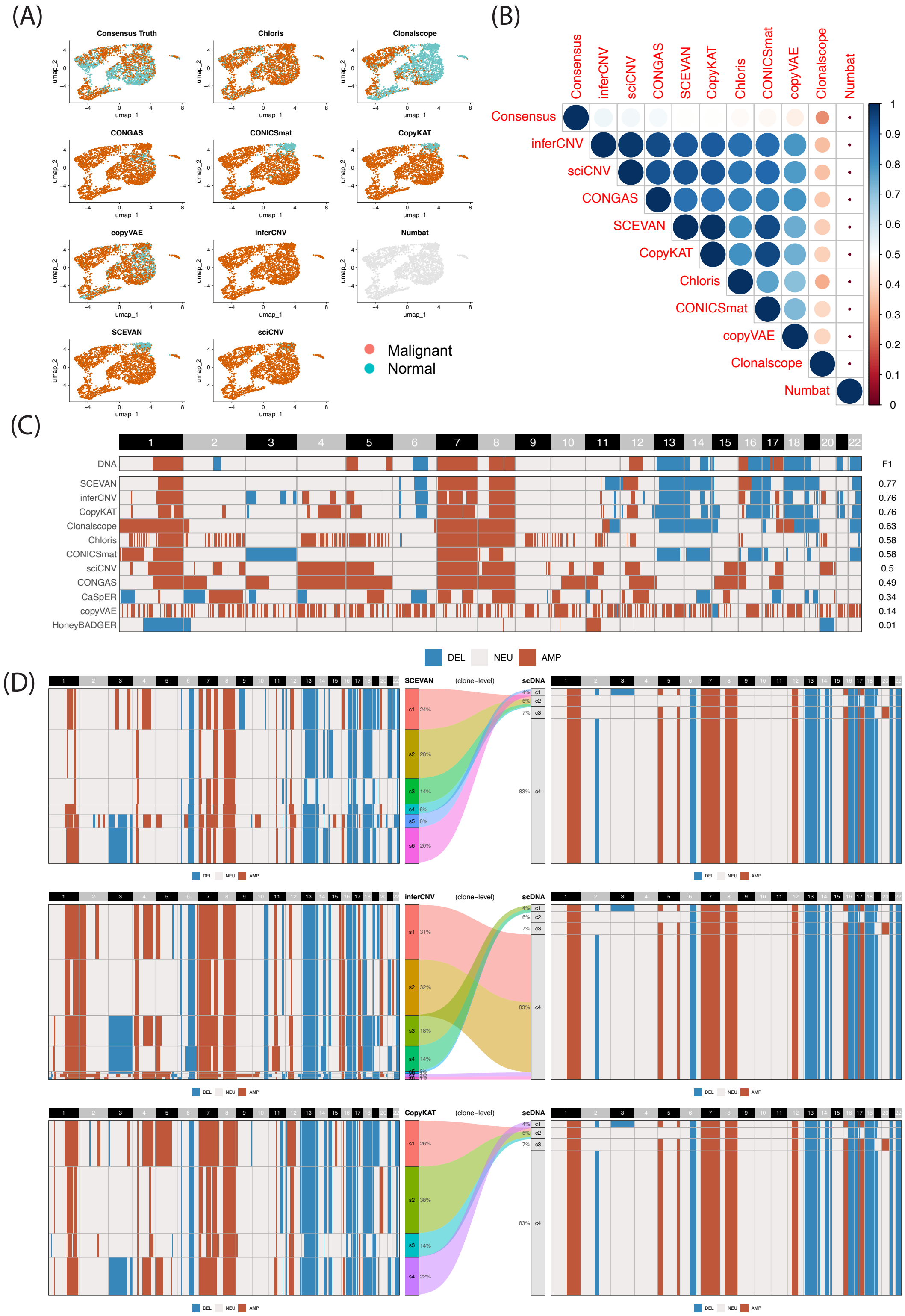


**Supplementary Figure 30: Visualization of IPM-BO-46-late result for B1 and B2 tasks.** (A) UMAP of malignant cell classification. (B) Cross-method concordance score based on the Jaccard index. (C) Bulk-level CNV profiles inferred from scRNA-based CNV methods, with ground truth derived from single-cell DNA sequencing. (D) Subclonal assignments of the top three methods at the bulk level. Only subclones with F1 > 0.6 are assigned.

#### IPM-BO-53-early

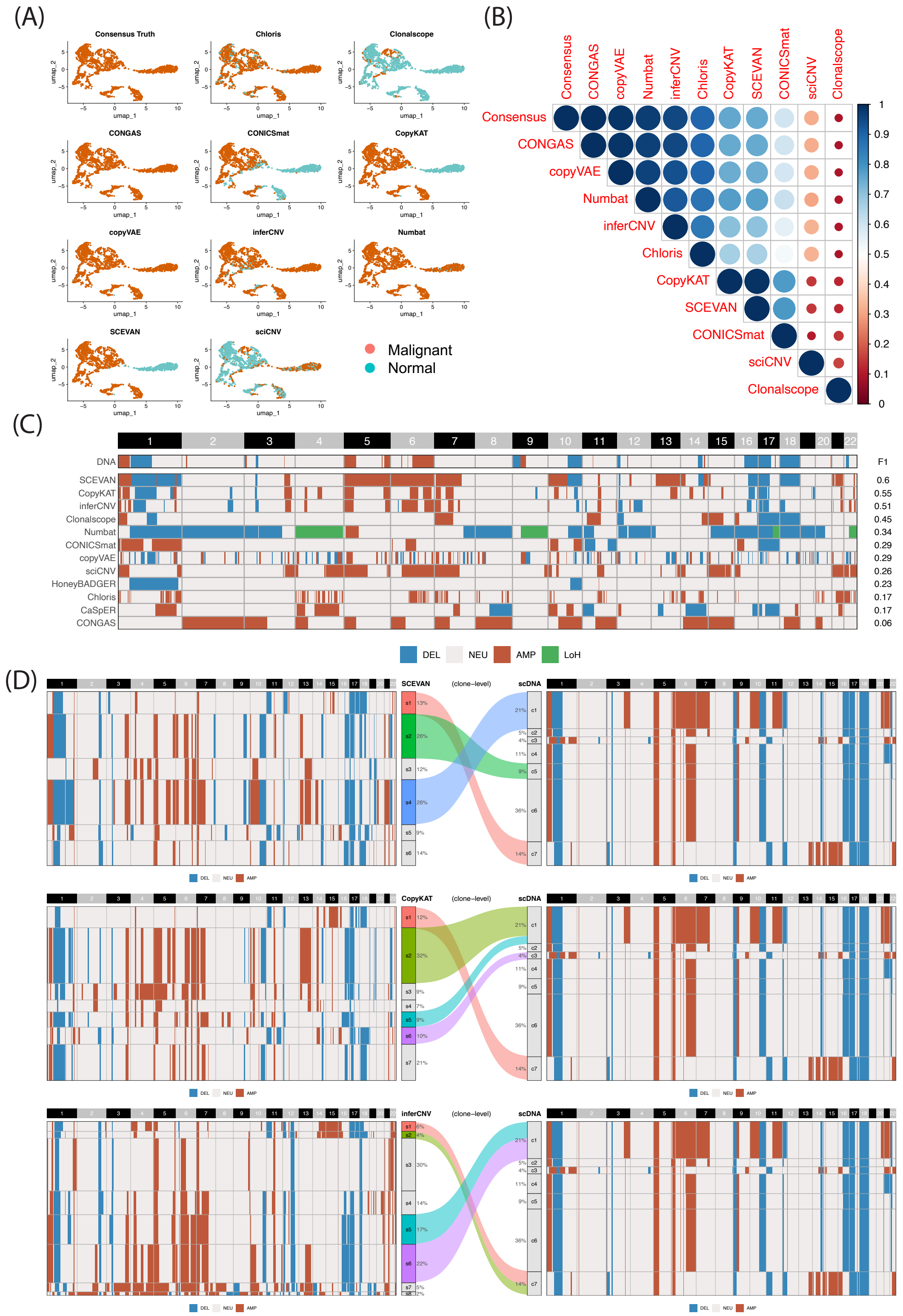


**Supplementary Figure 31: Visualization of IPM-BO-53-early result for B1 and B2 tasks.** (A) UMAP of malignant cell classification. (B) Cross-method concordance score based on the Jaccard index. (C) Bulk-level CNV profiles inferred from scRNA-based CNV methods, with ground truth derived from single-cell DNA sequencing. (D) Subclonal assignments of the top three methods at the bulk level. Only subclones with F1 > 0.6 are assigned.

### IPM-BO-53-late

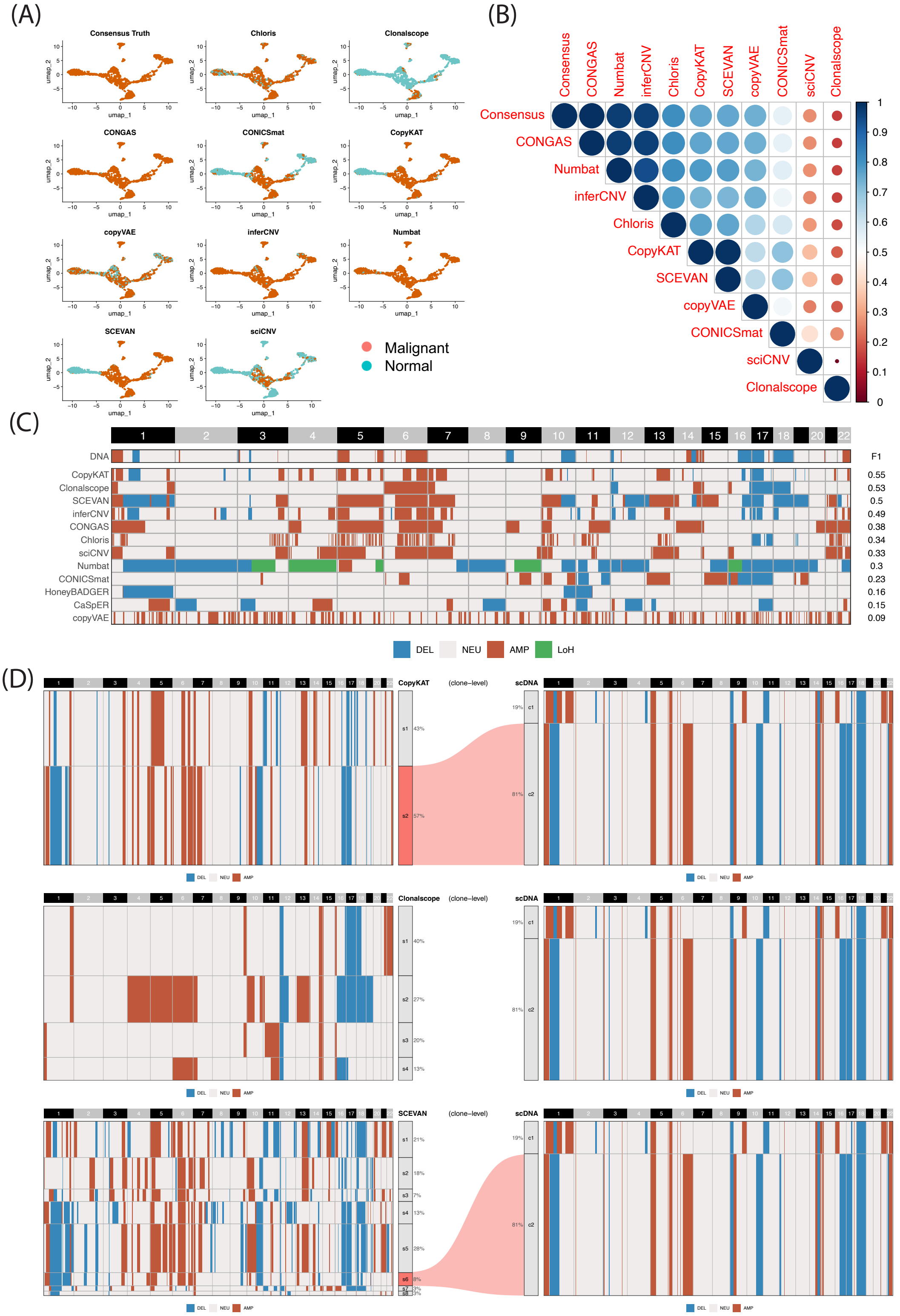


**Supplementary Figure 32: Visualization of IPM-BO-53-late result for B1 and B2 tasks.** (A) UMAP of malignant cell classification. (B) Cross-method concordance score based on the Jaccard index. (C) Bulk-level CNV profiles inferred from scRNA-based CNV methods, with ground truth derived from single-cell DNA sequencing. (D) Subclonal assignments of the top three methods at the bulk level. Only subclones with F1 > 0.6 are assigned.

#### IPM-BO-86-early

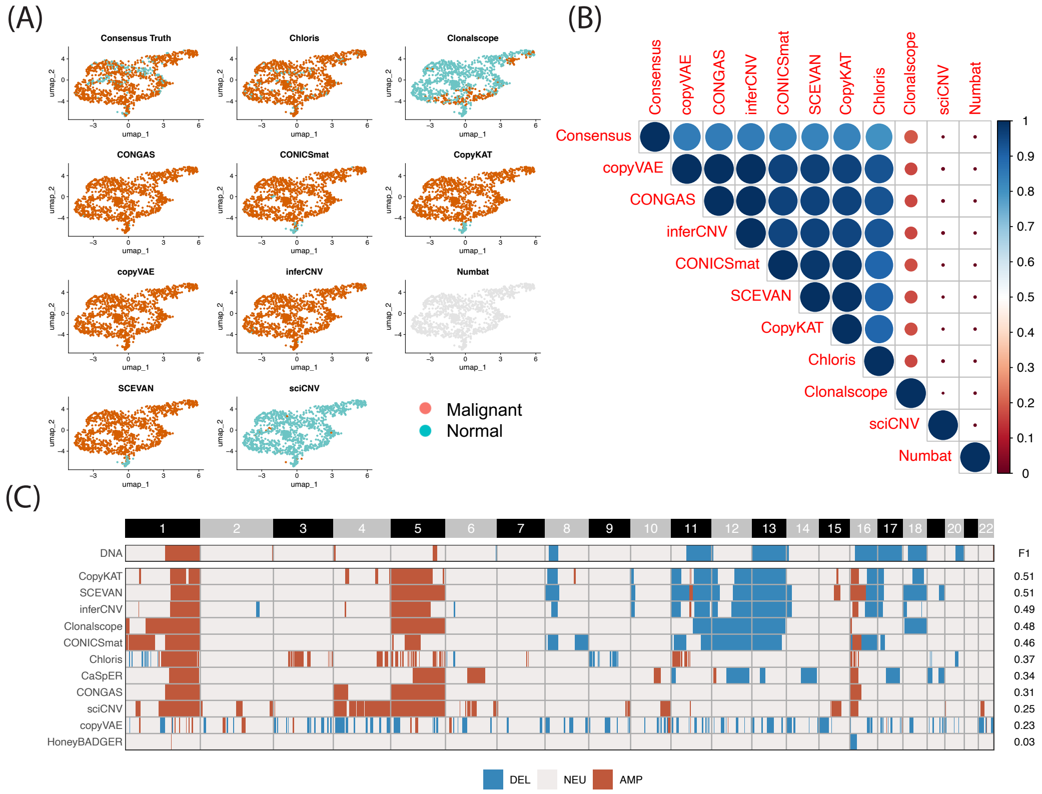


**Supplementary Figure 33: Visualization of IPM-BO-86-early result for B1 and B2 tasks.** (A) UMAP of malignant cell classification. (B) Cross-method concordance score based on the Jaccard index. (C) Bulk-level CNV profiles inferred from scRNA-based CNV methods, with ground truth derived from bulk DNA sequencing.

#### IPM-BO-86-late

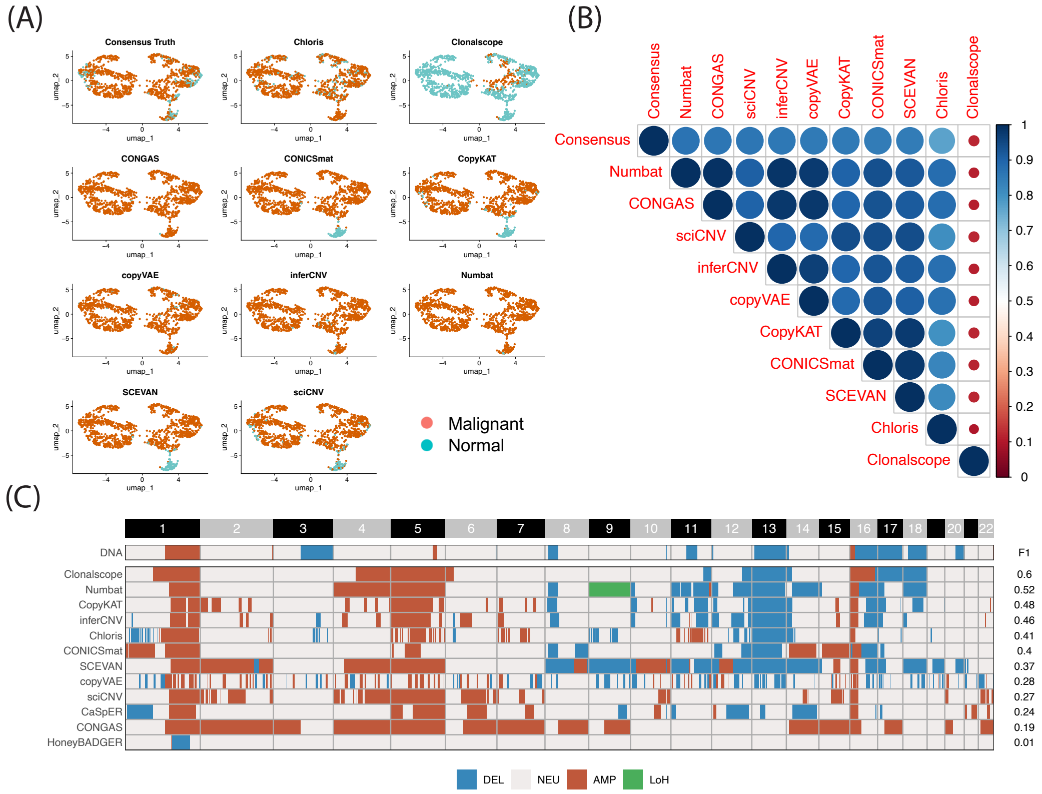


**Supplementary Figure 34: Visualization of IPM-BO-86-late result for B1 and B2 tasks.** (A) UMAP of malignant cell classification. (B) Cross-method concordance score based on the Jaccard index. (C) Bulk-level CNV profiles inferred from scRNA-based CNV methods, with ground truth derived from bulk DNA sequencing.

#### IPM-BO-139-early

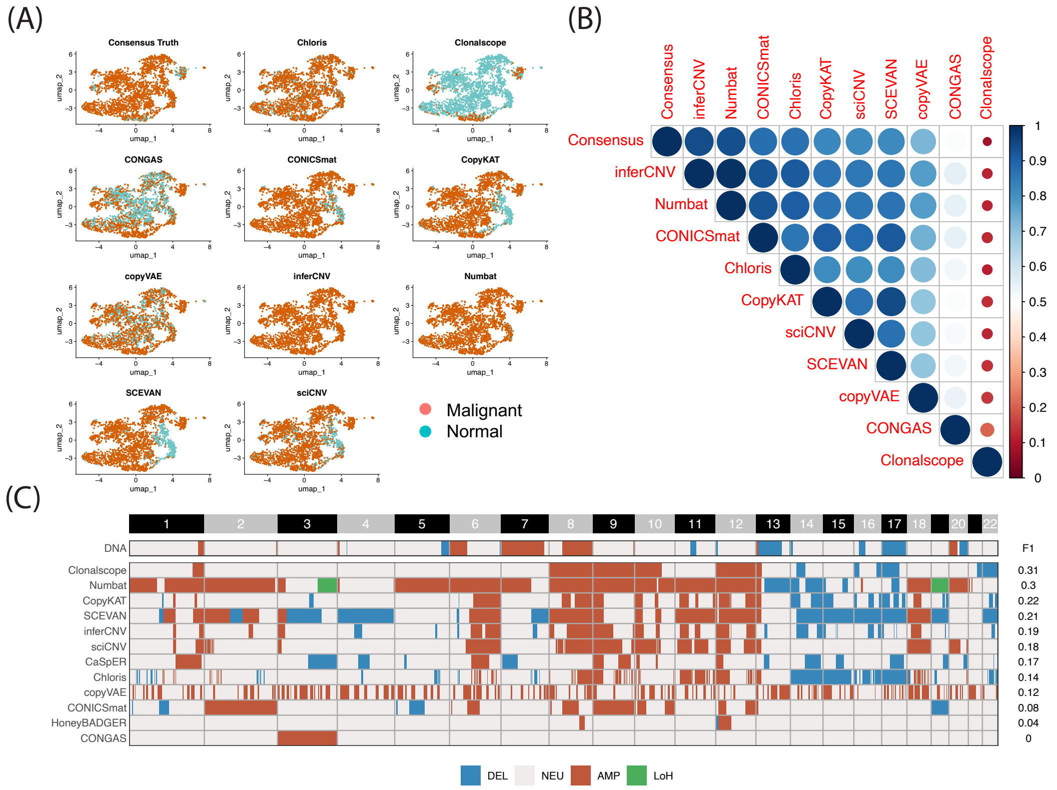


**Supplementary Figure 35: Visualization of IPM-BO--early result for B1 and B2 tasks.** (A) UMAP of malignant cell classification. (B) Cross-method concordance score based on the Jaccard index. (C) Bulk-level CNV profiles inferred from scRNA-based CNV methods, with ground truth derived from bulk DNA sequencing.

#### IPM-BO-139-late

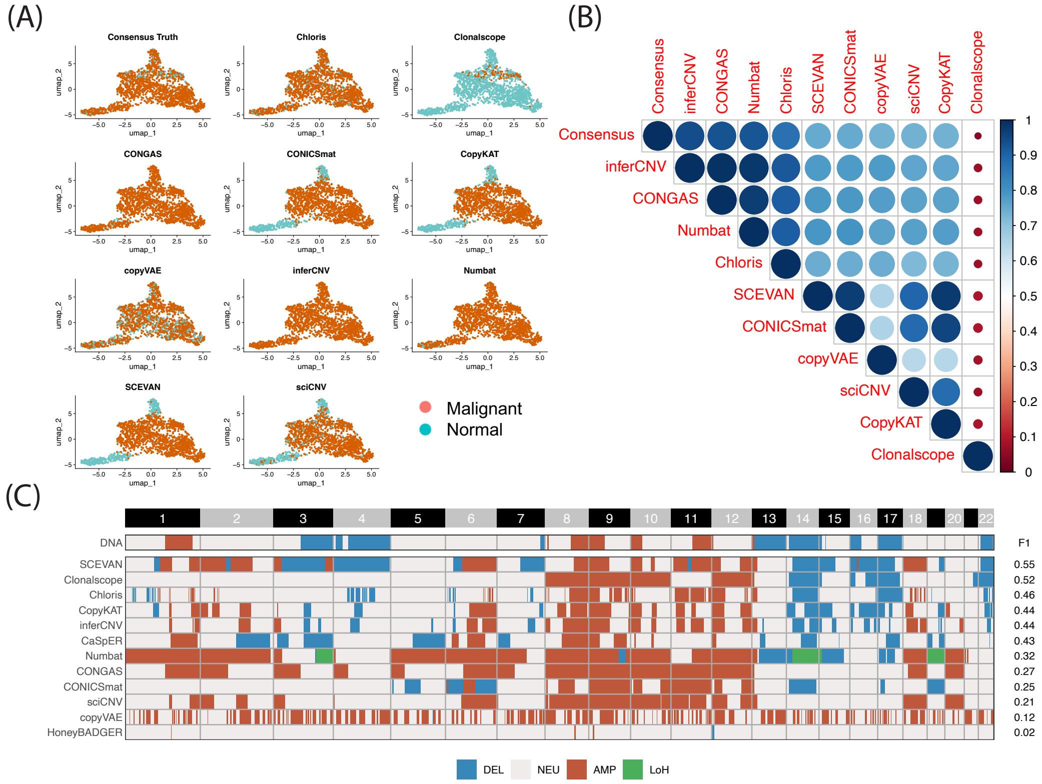


**Supplementary Figure 36: Visualization of IPM-BO-139-late result for B1 and B2 tasks.** (A) UMAP of malignant cell classification. (B) Cross-method concordance score based on the Jaccard index. (C) Bulk-level CNV profiles inferred from scRNA-based CNV methods, with ground truth derived from bulk DNA sequencing.

### De novo simulation results

#### Ground-truth CNV profile and $C_g$ distribution

(A)

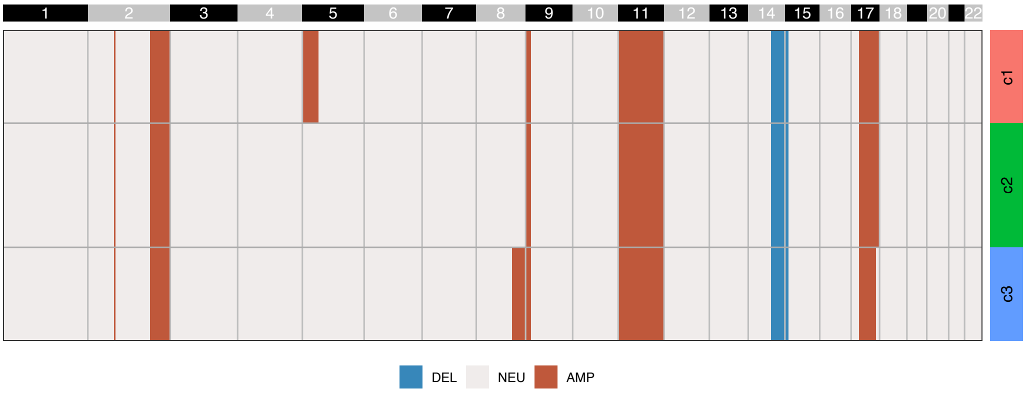


(B)

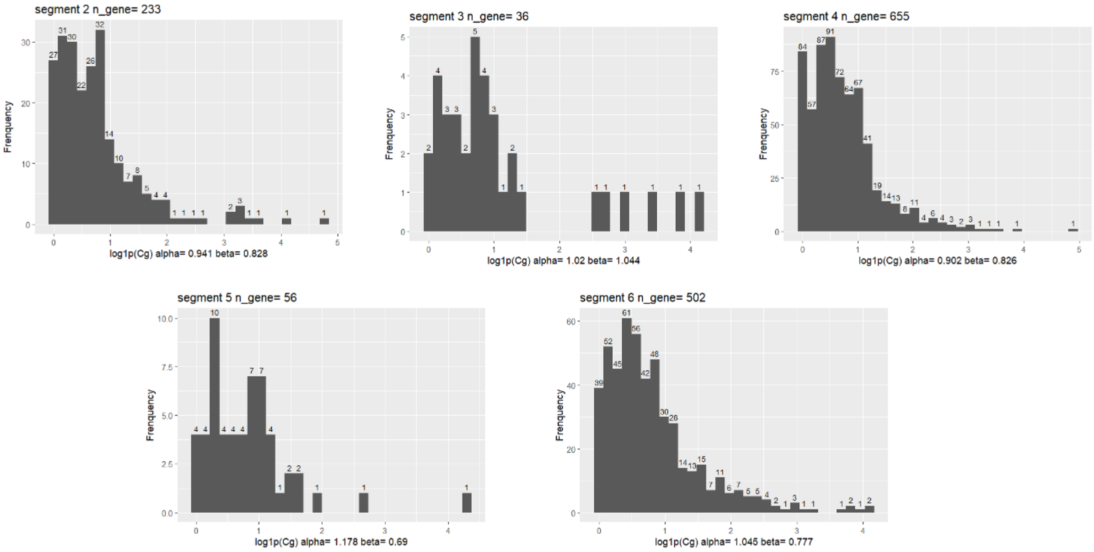


**Supplementary Figure 37: Visualization of ground-truth CNV profile and DNA-to-RNA translation strength ($C_g$) distribution.** (A) Modified CNV profile from SNU638 as ground truth for de novo simulation. The deletion segment on chromosome 14 was expanded to ensure that CNV deletion is detectable under strong-signal scenarios. (B) Empirical estimates and fitted distribution of $\log(1 + C_g)$.

#### Clonalscope

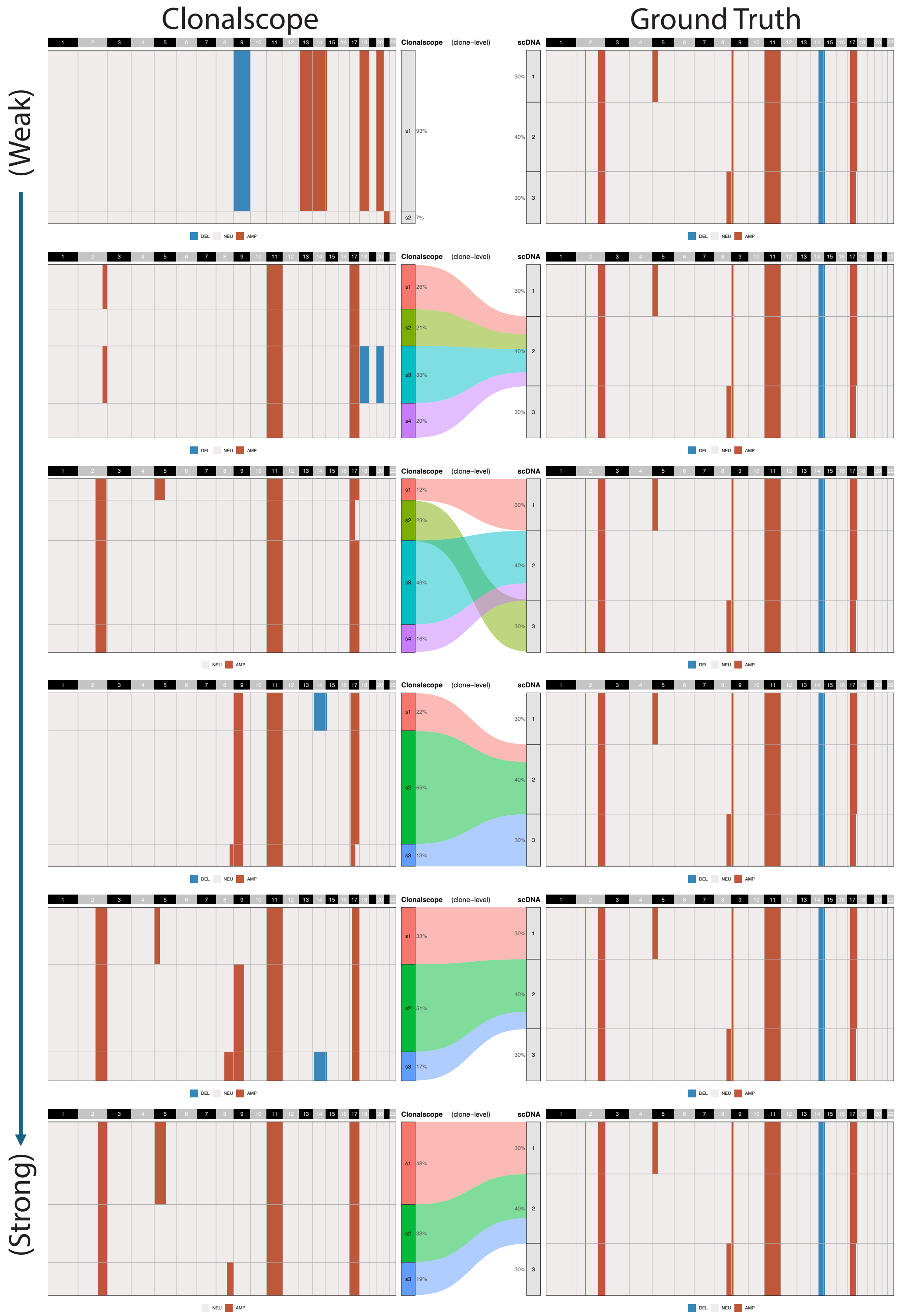


**Supplementary Figure 38: Visualization of Clonalscope on de novo simulation data.** The plots show CNV heatmaps of Clonalscope (left) and the ground truth (right). Six simulation scenarios are shown, ordered from weakest to strongest signal. Each predicted subclone is mapped to the most similar ground-truth subclone, while spurious subclones (i.e., maximum F1 score < 0.6) are excluded from mapping.

#### CopyKAT

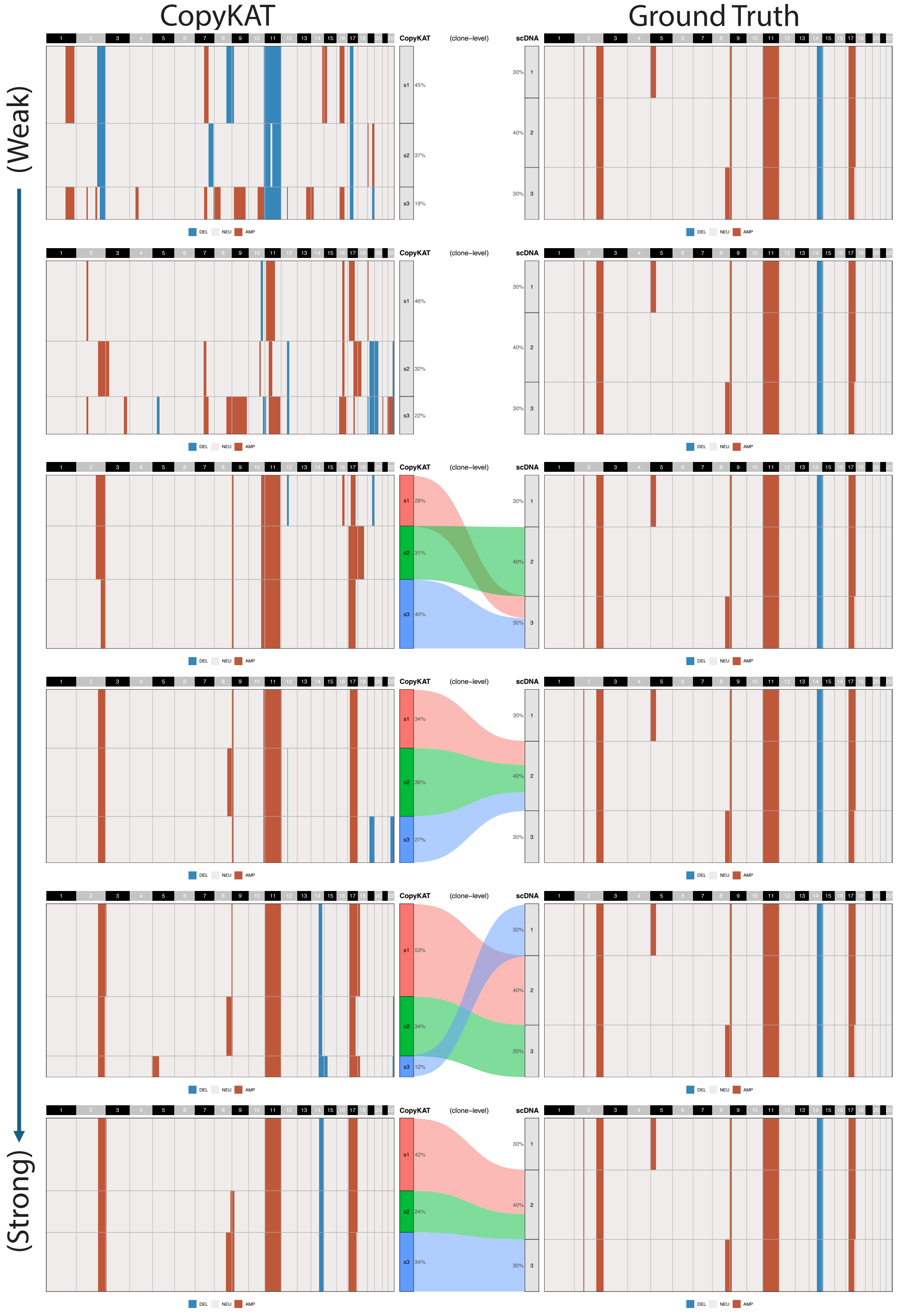


**Supplementary Figure 39: Visualization of CopyKAT on de novo simulation data.** The plots show CNV heatmaps of CopyKAT (left) and the ground truth (right). Six simulation scenarios are shown, ordered from weakest to strongest signal. Each predicted subclone is mapped to the most similar ground-truth subclone, while spurious subclones (i.e., maximum F1 score < 0.6) are excluded from mapping.

#### inferCNV

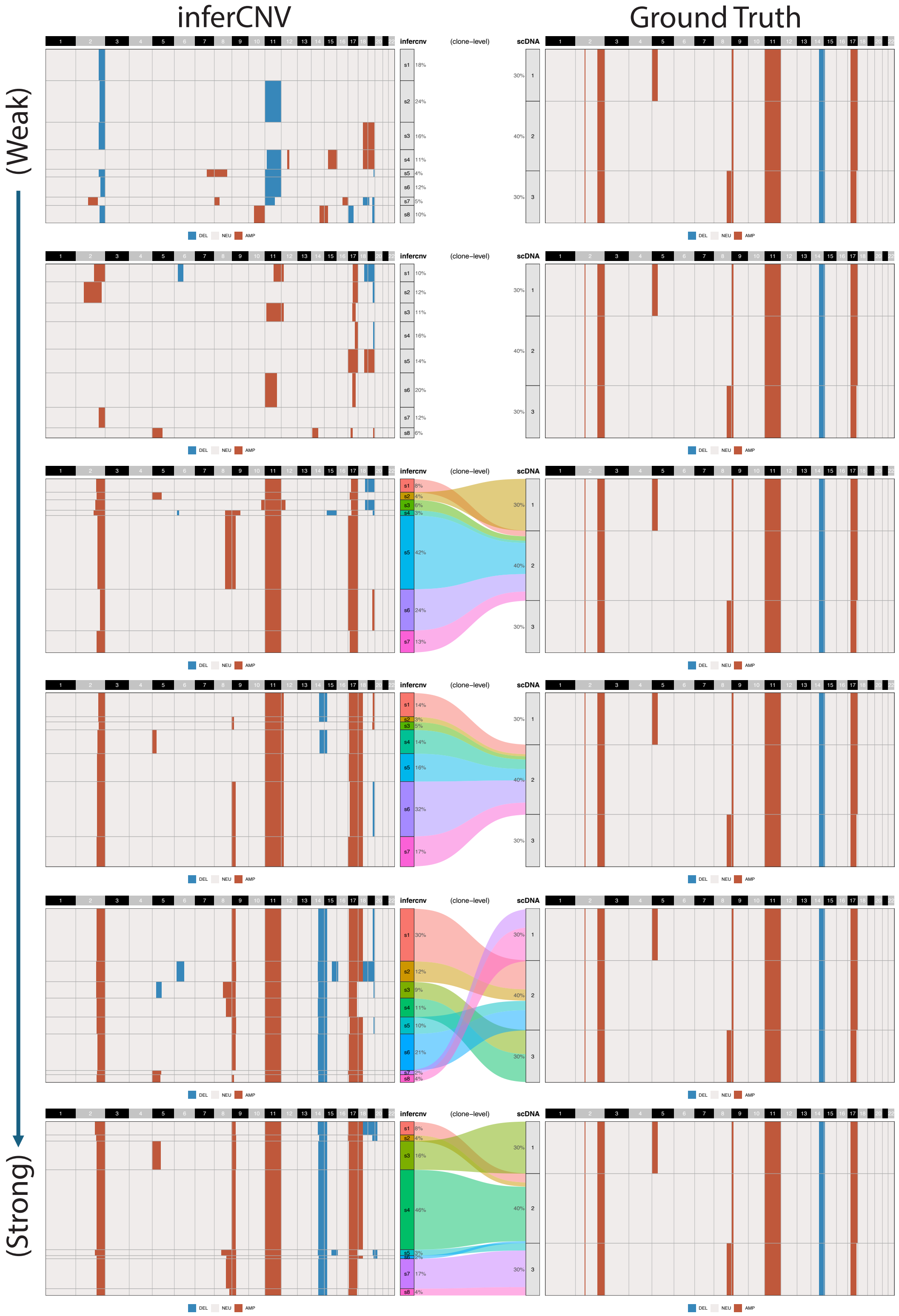


**Supplementary Figure 40: Visualization of inferCNV on de novo simulation data.** The plots show CNV heatmaps of inferCNV (left) and the ground truth (right). Six simulation scenarios are shown, ordered from weakest to strongest signal. Each predicted subclone is mapped to the most similar ground-truth subclone, while spurious subclones (i.e., maximum F1 score < 0.6) are excluded from mapping.

#### Numbat

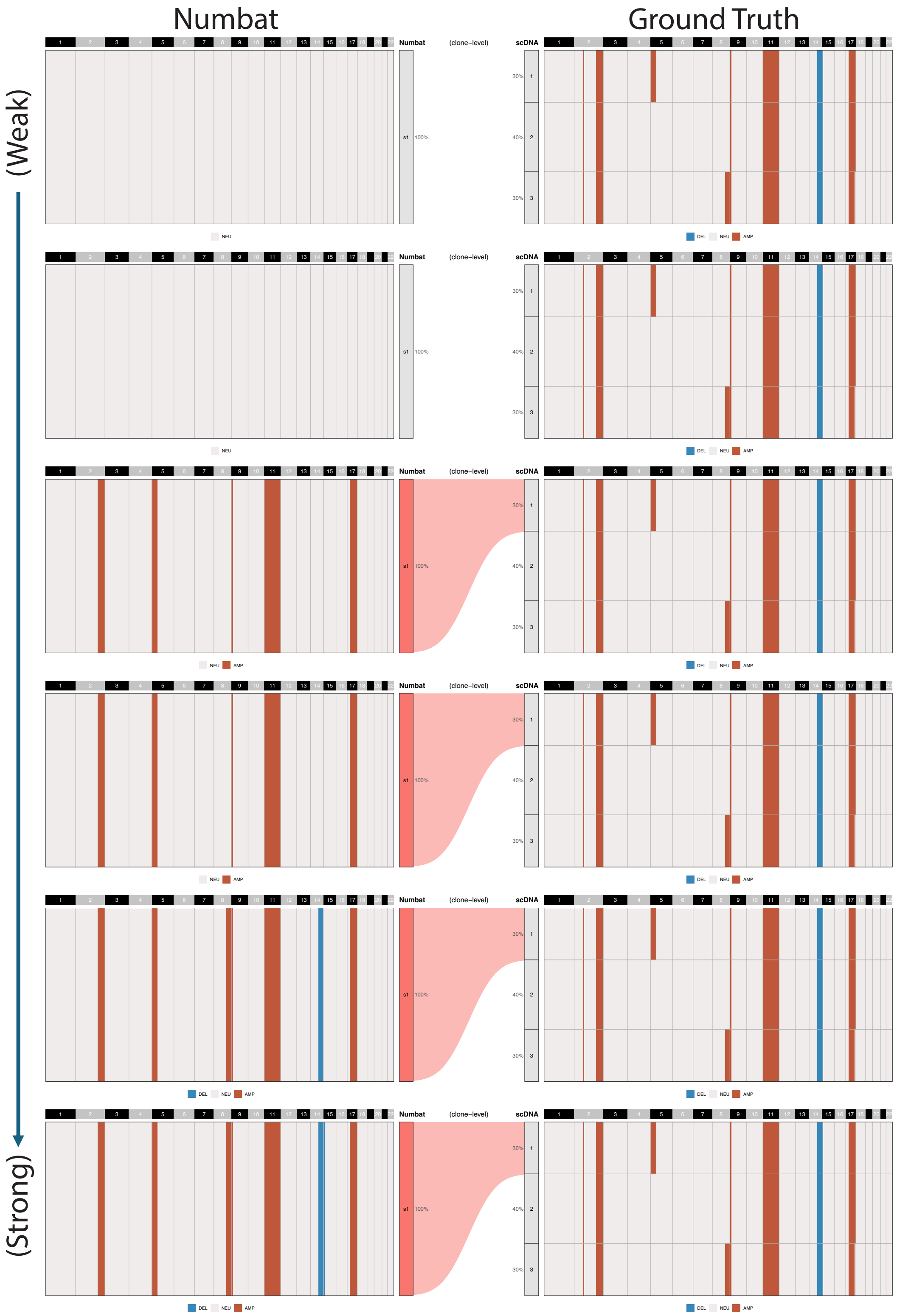


**Supplementary Figure 41: Visualization of Numbat on de novo simulation data.** The plots show CNV heatmaps of Numbat (left) and the ground truth (right). Six simulation scenarios are shown, ordered from weakest to strongest signal. Each predicted subclone is mapped to the most similar ground-truth subclone, while spurious subclones (i.e., maximum F1 score < 0.6) are excluded from mapping.

#### SCEVAN

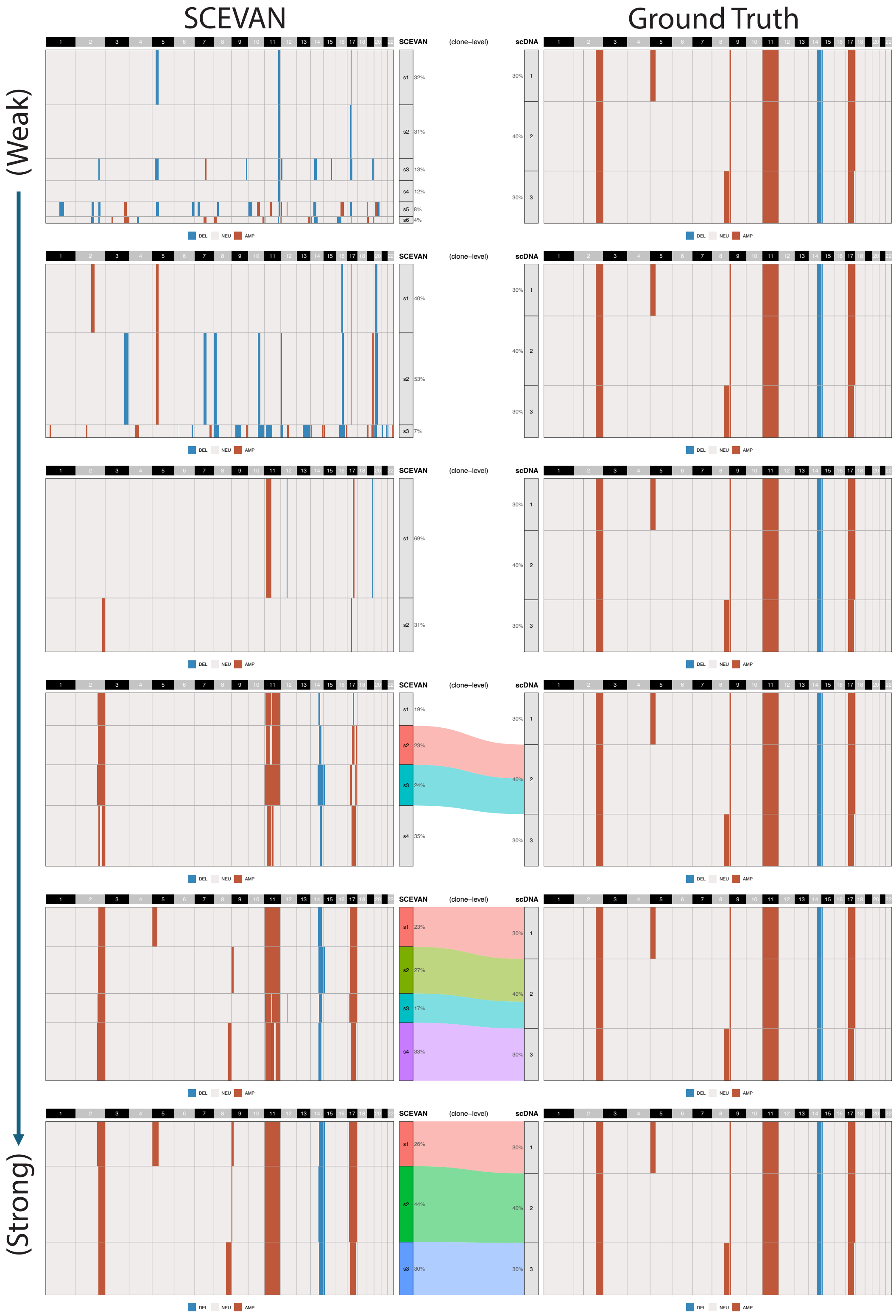


**Supplementary Figure 42: Visualization of SCEVAN on de novo simulation data.** The plots show CNV heatmaps of SCEVAN (left) and the ground truth (right). Six simulation scenarios are shown, ordered from weakest to strongest signal. Each predicted subclone is mapped to the most similar ground-truth subclone, while spurious subclones (i.e., maximum F1 score < 0.6) are excluded from mapping.

### Association between SNP-level depth and CNV inference performance

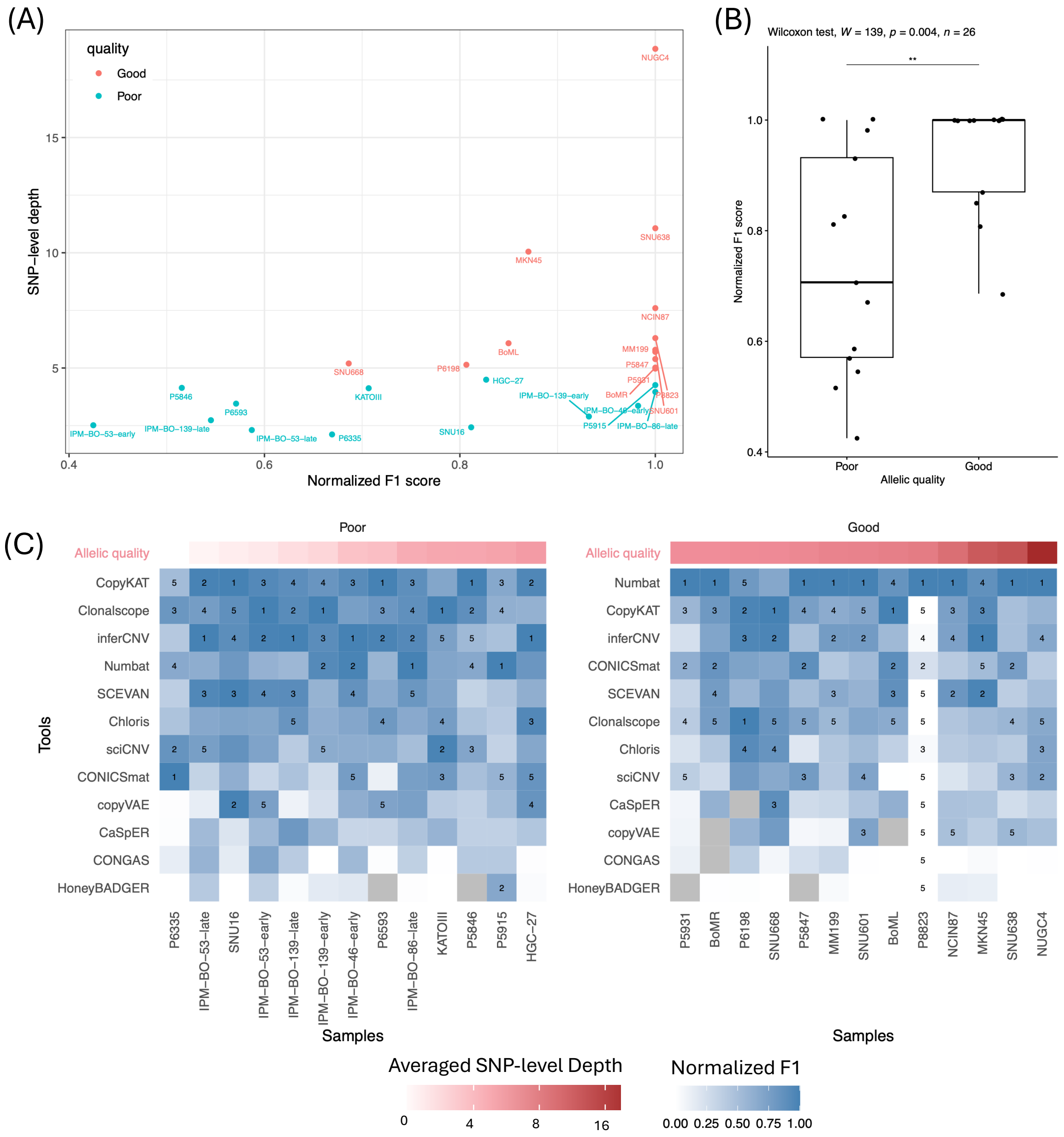

**Supplementary Figure 43: Association between SNP-level depth and CNV inference performance.** (A) Scatter plot showing the relationship between SNP-level sequencing depth and normalized F1 score for Numbat across 28 datasets. Each point represents one sample. Colors indicate groups with poor and good allelic quality, separated by an SNP-depth threshold of 5. (B) Comparison of normalized F1 scores between poor (<5) and good (≥5) SNP-depth groups using the Wilcoxon rank-sum test, demonstrating significantly improved performance under higher allelic coverage. (C) Heatmap of normalized F1 scores across all methods and samples. The top five methods for each sample are labeled. Samples are ordered by SNP-level depth to highlight performance differences across coverage conditions.

### Scalability

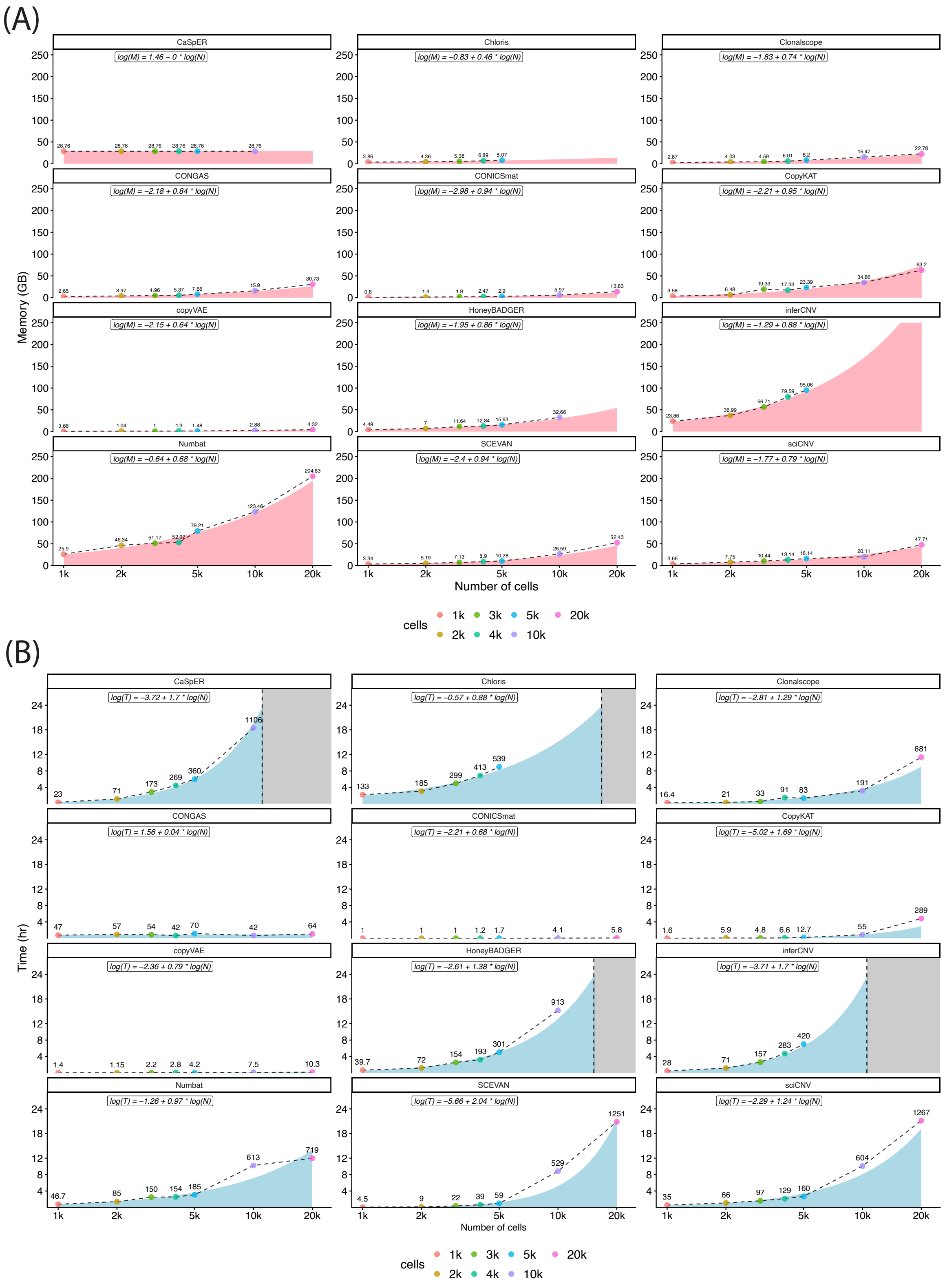


**Supplementary Figure 44: Scalability assessment of CNV inference tools.** The scalability of each CNV inference tool was assessed based on (A) runtime and (B) memory usage. Dashed lines represent observed runtime and memory usage, and the curve areas indicate values predicted by log–log regression models. The fitted regression equations are shown at the top of each panel.

#### Usability

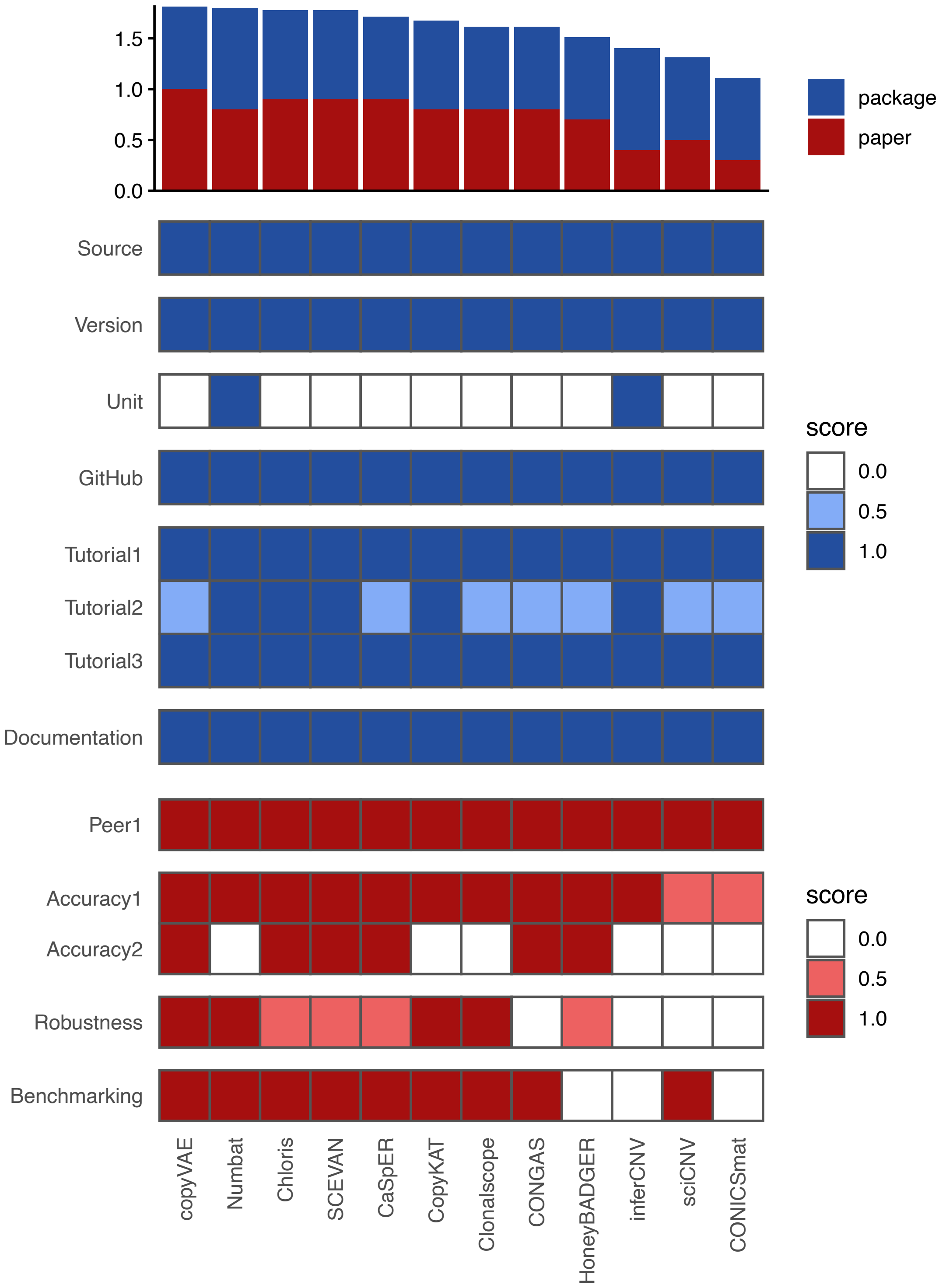


**Supplementary Figure 45: Usability assessment of CNV inference tools.** The usability of each CNV inference tools was assessed via ten categories (see Methods) that consider criteria related to the implementation of the methods (package; blue) and information included in the original publications (paper; red). Scores are visualized as a heatmap, with methods ordered by overall usability. The overall score is calculated as the sum of the average package- and paper-based usability scores and is displayed above as a bar plot.

### Robustness

#### CNV-state cutoff

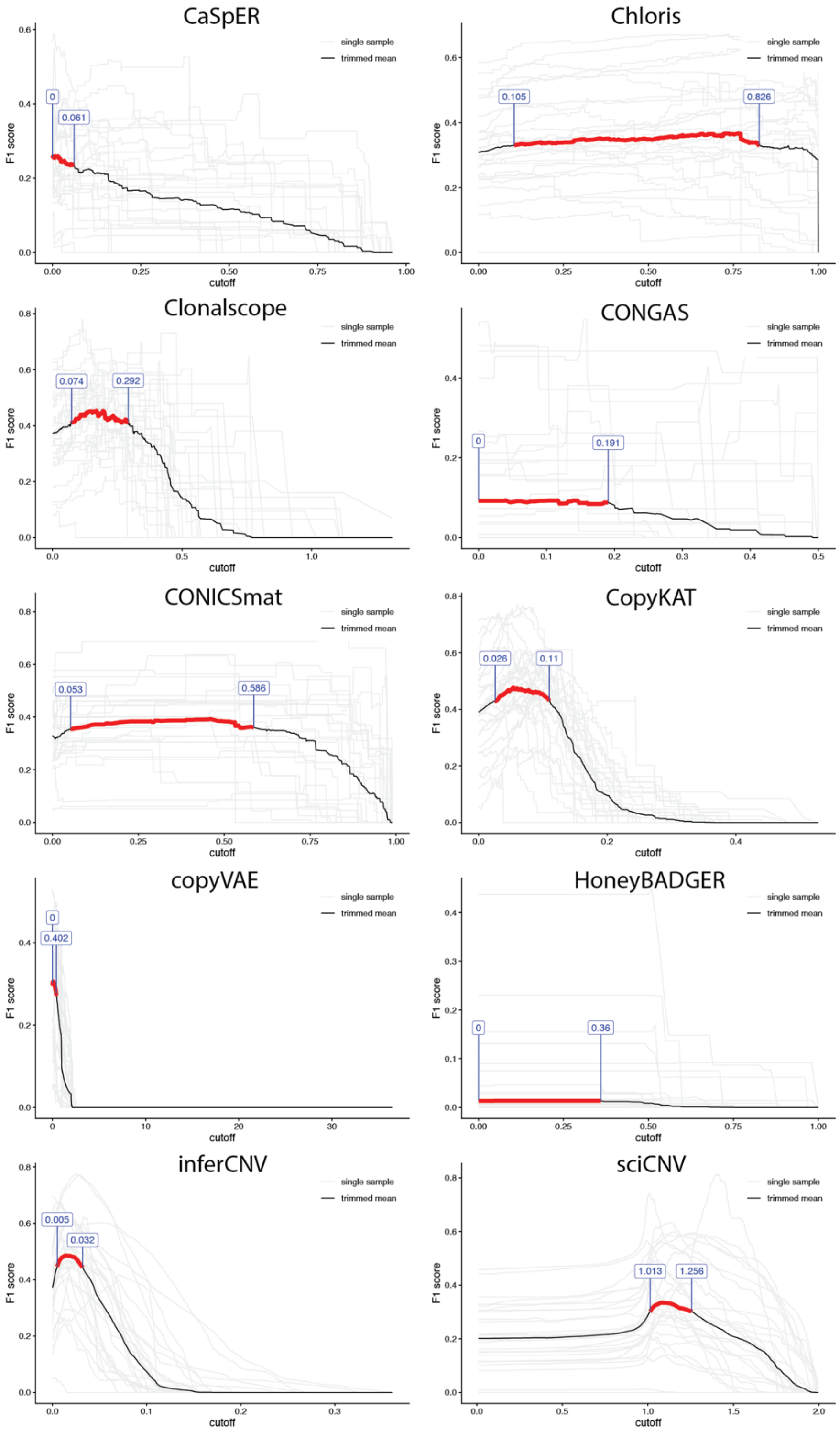


**Supplementary Figure 46: Robustness of CNV-state cutoff evaluated using trimmed mean F1 score.** Performance was summarized using a trimmed mean F1 score, computed by removing the top and bottom 25% of values and averaging the remaining 50%. The trimmed mean is shown as a solid line. Cutoff values achieving ≥90% of the maximum trimmed mean F1 score are highlighted in red as a recommended operating range.

### Validation of ground-truth CNV profiles

#### Cross-modality concordance

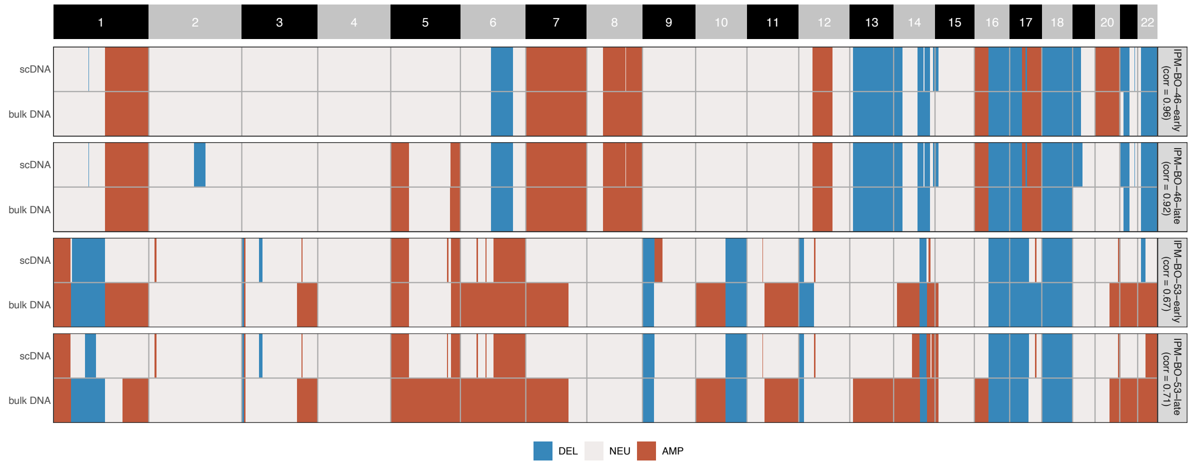


**Supplementary Figure 47: Cross-modality concordance between bulk WGS and single-cell WGS.** Among the four PDO samples (IPM-BO46-early, IPM-BO46-late, IPM-BO53-early, and IPM-BO53-late), single-cell CNV profiles are consistent with bulk CNV profiles. The correlations between bulk DNA and single-cell DNA in IPM-BO46-early and IPM-BO46-late are greater than 0.9. In IPM-BO53-early and IPM-BO53-late, single-cell CNV profiles appear slightly more conservative than bulk profiles; however, the correlations remain high, around 0.7.

### Literature validation of recurrent CNV events

(A)

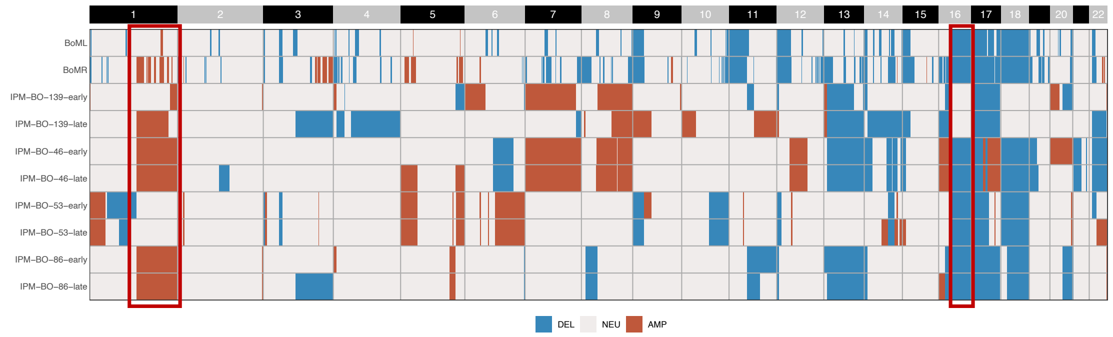


(B)

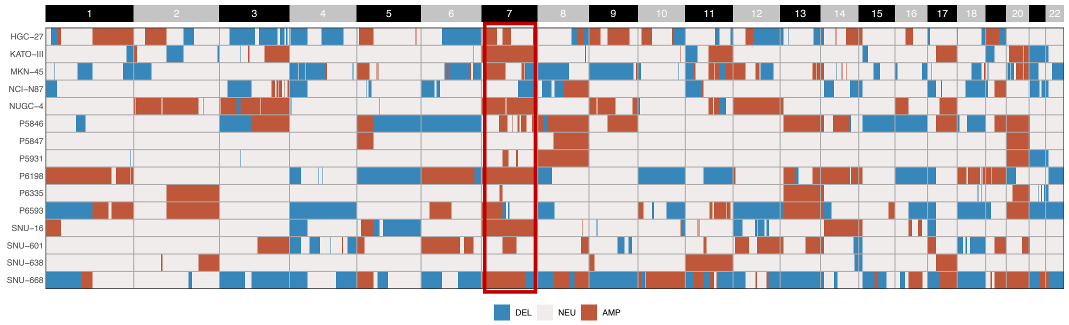


**Supplementary Figure 48: Literature validation of recurrent CNV events in breast cancer and gastric cancer.** The figures highlight recurrent CNV events observed in the analyzed samples of (A) breast cancer and (B) gastric cancer..

### Supplementary Table 1: Studies Comparison

**Supplementary Table 1: Overview and comparison of recent benchmarking studies for scRNA-seq CNV inference tools.** Comparison of recent benchmarking studies, including the number of methods, datasets, and evaluation design.

| | **Our study** | **Chen et al.[2]** | **Schmid et al.[3]** | **Song et al.[4]** |
|---|---|---|---|---|
| | | **Data and Methods** | | |
| Number of methods | 12 | 5 | 6 | 5 |
| Number of tumor samples | 28 | 7 | 20 | 14 |
| Number of cells | >100k | ~7k | ~60k | ~8k |
| # evaluation by scDNA | Primary: 3<br>Metastasis: 4<br>Cell line: 9<br>PDO: 4 | Primary: 1<br>Metastasis: 0<br>Cell line: 0<br>PDO: 0 | Primary: 4<br>Metastasis: 0<br>Cell line: 11<br>PDO: 0 | Primary: 12<br>Metastasis: 0<br>Cell line: 2<br>PDO: 0 |
| # evaluation by bulk DNA | Primary: 2<br>Metastasis: 2<br>Cell line: 0<br>PDO: 4 | Primary: 0<br>Metastasis: 0<br>Cell line: 1<br>PDO: 0 | Primary: 3<br>Metastasis: 0<br>Cell line: 2<br>PDO: 0 | Primary: 0<br>Metastasis: 0<br>Cell line: 0<br>PDO: 0 |
| Sample sources | Primary,<br>Metastasis,<br>Cell line,<br>PDO | Primary,<br>Cell line | Primary,<br>Cell line | Primary,<br>Cell line |
| Practical guideline | V | X | △ | △ |
| | | **Method Performance** | | |
| Tumor cell classification | V | X | V | V |
| Subclone construction | V | V | V | V |
| Bulk CNV inference | V | V | V | V |
| Subclone/cell CNV inference | V | X | V | V |
| De novo simulation | V | X | X | X |
| | | **Scalability & Usability** | | |
| Time benchmark | V | V | V | V |
| Memory benchmark | V | X | V | X |
| Usability | V | X | X | X |
| | | **Robustness & Sensitivity** | | |
| Sequencing depth | V | V | X | V |
| Normal reference | V | V | V | X |
| Intronic reads | V | X | X | X |
| CNV cutoff | V | X | X | X |
| V: Covered, X: Not covered, △: Partially covered | | | | |

### Supplementary Table 2: Introns Inclusion and Exclusion

**Supplementary Table 2: Comparison of sequencing metrics with and without inclusion of intronic reads.** The table summarizes key quality metrics, including "Reads Mapped Confidently to Transcriptome" and "Median Genes per Cell," for each sample under two counting strategies: exonic reads only and exonic plus intronic reads. Consistent with previous reports from 10x Genomics, inclusion of intronic reads increases the fraction of reads assigned to the transcriptome and improves gene detection sensitivity, as reflected by higher median genes per cell.

| Samples | Reads Mapped confidently to Transcriptome | | Median Gene per Cell | |
|---|---|---|---|---|
| | Exclusion | Inclusion | Exclusion | Inclusion |
| P5846 | 58.4% | 74.0% | 525 | 658 |
| P5847 | 65.7% | 78.5% | 511 | 615 |
| P5915 | 60.3% | 68.6% | 373 | 395 |
| P5931 | 61.6% | 76.7% | 682 | 1231 |
| P6198 | 60.8% | 76.3% | 2003 | 2946 |
| P6335 | 64.8% | 76.3% | 1496 | 2152 |
| P6593 | 46.4% | 63.6% | 809 | 978 |
| P8823 | 42.4% | 47.5% | 1090 | 1373 |
| **Average** | **57.55%** | **70.19%** | **936.13** | **1293.50** |
